# Examining chromatin heterogeneity through PacBio long-read sequencing of M.EcoGII methylated genomes: an m^6^A detection efficiency and calling bias correcting pipeline

**DOI:** 10.1101/2023.11.28.569045

**Authors:** Allison F. Dennis, Zhuwei Xu, David J. Clark

## Abstract

Recent studies have combined DNA methyltransferase footprinting of genomic DNA in nuclei with long-read sequencing, resulting in detailed chromatin maps for multi-kilobase stretches of genomic DNA from one cell. Theoretically, nucleosome footprints and nucleosome-depleted regions can be identified using M.EcoGII, which methylates adenines in any sequence context, providing a high-resolution map of accessible regions in each DNA molecule. Here we report PacBio long-read sequence data for budding yeast nuclei treated with M.EcoGII and a bioinformatic pipeline which corrects for three key challenges undermining this promising method. First, detection of m^6^A in individual DNA molecules by the PacBio software is inefficient, resulting in false footprints predicted by random gaps of seemingly unmethylated adenines. Second, there is a strong bias against m^6^A base calling as AT content increases. Third, occasional methylation occurs within nucleosomes, breaking up their footprints. After correcting for these issues, our pipeline calculates a correlation coefficient-based score indicating the extent of chromatin heterogeneity within the cell population for every gene. Although the population average is consistent with that derived using other techniques, we observe a wide range of heterogeneity in nucleosome positions at the single-molecule level, probably reflecting cellular chromatin dynamics.

## Introduction

The advent of long-read sequencing has opened up many new and exciting avenues of research. Recent papers in the chromatin field have described global methods to map nucleosome positions using various DNA methyltransferases to methylate exposed linker DNA between nucleosomes (1–6). The basic methylation footprinting approach was developed for single genes many years ago, involving enzymatic methylation followed by detection using bisulphite sequencing (7–12). Although this approach works well, bisulphite treatment results in significant DNA damage and, in practical terms, limits the observation of methylation along the same DNA molecule to only a few hundred base pairs. In contrast, sequencing using the Nanopore or Pacific BioSciences (PacBio) platforms facilitates the observation of methylation in multi-kilobase DNA molecules. Since the sequenced DNA molecules are original (i.e., unamplified), modified bases such as m^5^C and m^6^A can be directly detected in each DNA molecule as it is sequenced. At present, Nanopore is preferred for m^5^C (13), whereas PacBio is preferred for m^6^A (1). A major advantage of long reads is that they allow analysis of potential correlations between the methylation statuses of regions in the same molecule, such as enhancers and promoters (2).

The resolution of enzymatic methylation footprinting depends on the frequency of methylatable bases, as limited by the recognition sequence of a given methyltransferase. For example, *E. coli dam* methyltransferase methylates A in the sequence GATC, but the resolution is too low for footprinting because GATC sites are relatively scarce (about 1 in 256 bp in a genome with 50% AT). The potential resolution of the SssI methyltransferase (M.SssI), which methylates C in the sequence CG, or M.CviPI, which methylates C in the sequence GC, is much higher (about 1 in 16 bp), and that of EcoGII methyltransferase (M.EcoGII) and similar enzymes, which methylate any A (2,14), is very high indeed (about 1 in 2 bp). Consequently, M.EcoGII is the enzyme of choice for high resolution methylation footprinting (1–3).

However, we show here that the PacBio software fails to detect most of the m^6^A in completely methylated DNA at the single molecule level. Thus, the efficiency of m^6^A detection by PacBio is a critical parameter that must be accounted for when constructing chromatin maps from single cells. The issue is that the lower the detection efficiency, the more likely a false footprint will be called, because the random gaps between methylated bases average longer. We present a bioinformatic pipeline to analyse m^6^A methylation patterns in PacBio data for budding yeast which corrects for this problem. An additional issue is that we observe significant methylation within nucleosomes, which effectively breaks up the nucleosome footprint. Furthermore, m^6^A base calling decreases with increasing AT-content and with the length of poly(A) sequences. All these problems are addressed by our pipeline. While other eukaryotic genomes may contain m^6^A as a result of epigenetic modification (15), budding yeast DNA contains little or none, and is therefore a good template for M.EcoGII methylation footprinting. Since our yeast strain is haploid, a comparison of long reads for each gene indicates the extent of chromatin heterogeneity within the cell population. We observe a very wide range of heterogeneity, even though the population average is consistent with that derived using other techniques, such as MNase-seq.

## Materials and Methods

### M.EcoGII methylation of nuclei

Yeast strain YDC111 (***MATɑ*** *ade2-1 can1-100 leu2-3,112 trp1-1 ura3-1* (16)) was grown at 30°C in synthetic complete (SC) medium (2% D-glucose, 6.7 g/l yeast nitrogen base (Sunrise Science Products 1501-250) and 0.79 g/l Complete Supplement Mixture with adenine (Sunrise Science Products 1128-010) to A_600_ ∼0.2 and arrested in G1 by the addition of ɑ-factor (FDA Core Facility) to 10 μg/ml. Arrest was monitored by observance of the “shmoo” phenotype in a light microscope. After 2.5 h, the cells were harvested by filtration and stored at −80°C. Nuclei were prepared as described (17), with a few minor changes. Spheroplasts were prepared from ∼100 A_600_ units of cells in 15 ml SM Buffer (SC medium with 1 M sorbitol, 50 mM Tris-HCl pH 8.0, 20 mM 2-mercaptoethanol) by digestion with ∼19,000 units of lyticase (Sigma L-2524) at 30°C for 10 min. Digestion of the cell wall was monitored by measuring the A_600_ of 30 μl cell suspension in 1 ml 1% SDS and considered complete when the A_600_ decreased to < 10% of the initial value. Spheroplasts were collected by centrifugation in a pre-cooled Sorvall SA600 rotor (8150 g for 5 min at 4°C), washed once with 25 ml cold ST Buffer (1 M sorbitol, 50 mM Tris-HCl pH 8.0), and lysed by resuspension in 20 ml cold F Buffer (18% w/v Ficoll-PM400 (GE Healthcare 17-0300-50), 40 mM potassium phosphate, 1 mM magnesium chloride, pH 6.5; protease inhibitors (Roche 05056489001) and 5 mM 2-mercaptoethanol were added just before use). The lysate was applied to a step gradient of 15 ml cold FG Buffer (7% w/v Ficoll-PM400, 20% glycerol, 40 mM potassium phosphate, 1 mM magnesium chloride, pH 6.5, with protease inhibitors and 5 mM 2-mercaptoethanol as above) and collected using the SA600 rotor (22640 g for 20 min at 4°C).

To perform methylation, the pellet of crude nuclei was resuspended in 4.8 ml pre-warmed NEB buffer 2.1 (50 mM NaCl, 10 mM Tris-HCl, 10 mM MgCl_2_, 0.1 mg/ml BSA, pH 7.9) with 1.2 mM S-adenosylmethionine (SAM) and protease inhibitors, and divided into ten 425 µl aliquots. M.EcoGII (24 µl; 600 units at 25 units/µl; New England Biolabs M0603B-HC1) or 24 µl water was added to each of five aliquots, mixed thoroughly but gently, and incubated at 30°C for 20 min. Methylation was stopped by adding 50 μl 200 mM EDTA, 10% SDS and incubating at 25°C for 5 min. Genomic DNA was purified by addition of 27.5 μl 10% SDS, mixing, addition of 133 μl 5 M potassium acetate, followed by two extractions with an equal volume of chloroform, precipitation with 0.7 vol. isopropanol, and one wash with 75% ethanol. The purified genomic DNA was dissolved in 20 μl 10 mM Tris-HCl pH 8.0, 0.5 mg/ml RNase A and incubated at 37°C for 4 h. To verify methylation, purified genomic DNA was digested with either DpnI, which only cleaves when its recognition site (GATC) is methylated, or MboI, which recognises the same site but is blocked by methylation, and evaluated by gel electrophoresis.

Genomic DNA positive control samples (gDNA) were obtained by subjecting genomic DNA purified from yeast cells as above to three rounds of M.EcoGII treatment to maximise methylation (10-20 μg DNA in 160 μl NEB buffer 2.1 with 0.8 mM SAM, incubated with 50 to 100 units M.EcoGII at 37°C for 4 h). The DNA was purified after each round.

### PacBio SMRT sequencing

PacBio library preparation requires about 1 μg DNA. Genomic DNA samples were sheared to ∼2 kb in a total volume of 200 µl using a Covaris miniTube Clear on the Covaris M220 with shearing conditions as follows: Peak Incident Power: 8 W, Duty Factor: 20%, Cycles per Burst: 1000, Treatment time: 900 s. Sonicated DNA was purified using AMPure XP beads (Beckman-Coulter) at a 1:1 ratio, eluted in 22.5 µl AMPure Elution Buffer and checked for quality using the DNA12000 assay in an Agilent BioAnalyzer. SMRTbell libraries were prepared using PacBio barcoded adapters according to the manufacturer’s instructions. Libraries from methylated and unmethylated nuclei were pooled and loaded in a single SMRT cell in a 5:1 or 8:1 ratio to increase the number of reads from the methylated sample while generating a SMRT cell run-specific control for comparison. Data were collected using 20 h run times with a 2 h pre-immobilisation time and a pre-extension time depending on the average size of each library using a PacBio Sequel I sequencer. Two biological replicate experiments were performed (see Table S1 for a summary).

### Read quality control

We removed low quality reads from the data using the following steps: 1) Circular consensus sequencing (CCS) reads were obtained using the raw subreads and the CCS program from the PacBio SMRT Link 10.4 software with default parameters (the CCS read does not distinguish between m^6^A and A). 2) DNA molecules without a CCS read were excluded. 3) The number of subread passes and the average per base quality score for each CCS read were obtained. This quality score was used to filter the reads.

### Identification of m^6^A in PacBio sequencing data

We developed a pipeline to identify m^6^A bases in single molecules in SLURM on an HPC cluster (NIH Biowulf), after alignment to the sacCer3 version of the *S. cerevisiae* genome, with the following steps: 1) Aligned subreads derived from the same DNA molecule were grouped using the zero-mode waveguide (ZMW), which corresponds to a single DNA molecule. 2) The ipdSummary package included in the PacBio SMRT Link 10.4 software was used to predict m^6^A bases in each subread from the same DNA molecule (ZMW). 3) A single consensus read with called m^6^A bases was generated for each DNA molecule. 4) Consensus reads with a minimum average base quality score of 90 were selected for analysis (see above and Results). 5) A final bam file containing all the consensus reads, each representing a single DNA molecule, was generated. Each called m^6^A base is represented by a single base insertion at the corresponding adenine in the DNA sequence using the cigar string in the bam file. 6) The average adenine methylation was computed for each consensus read obtained from the subreads.

### Detection of m^6^A in pUC19 methylated by Dam or M.EcoGII

pUC19 was purified from a *dam-dcm-* strain of *E. coli* (NEB C2925I) to prevent endogenous DNA methylation, and linearised with SmaI. Three 350-μl reactions were prepared containing 7 μg linear pUC19 in NEB rCutSmart (rCS) buffer with 1 mM SAM and incubated for 4 h at 37°C, as follows: (1) Control; (2) +28 μl Dam at 8 U/μl (NEB M0222L); (3) +28 μl M.EcoGII at 5 U/μl (NEB M0603S). The DNA was purified using Qiagen PCR columns and quantified by measuring A_260_. Two biological replicate experiments were performed. To determine the extent of methylation at GATC sites, 0.3 μg DNA was digested with 2 U DpnI (NEB R0176S) or 5 U MboI (NEB R0147S) in 15 μl rCS buffer for 1 h at 37°C and analysed in a 1.2% agarose gel containing ethidium bromide. Bar-coded libraries were prepared from all six samples and sequenced in a PacBio Sequel IIe machine with a movie time of 10 h.

### Identification of nucleosomes and M.EcoGII-accessible regions in single DNA molecules

We developed an adjusted binomial model to identify M.EcoGII-accessible regions in each sequenced DNA molecule (i.e., regions with a relatively high density of m^6^A). These correspond to nucleosome-depleted regions (NDRs) and the linkers between nucleosomes. We predicted the expected local methylation in 25 bp windows (this window size is long enough to include sufficient adenines and short enough relative to the expected linker length) using two parameters: the number of adenines on both strands and the average methylation per read. For gDNA purified and then treated with M.EcoGII, we found that the window m^6^A methylation level is linearly related to the average methylation of the entire consensus read (see below). This linear relationship is influenced by the total number of adenines on both strands within the window (Formula 1). For a 25 bp window, we constructed ***N*** = 3 to 25 linear relationships (23 in total), to compute the expected methylation in each 25 bp window using linear regression (Table S2).

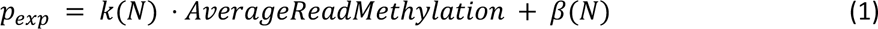

For a 25-bp window with ***n*** adenines (total count for both strands), the probability that ***X*** of ***n*** adenines are methylated should follow the binomial distribution *X*∼*B*(*n*, *p_exp_*), where *p_exp_* is the expected methylation in the window, estimated from AT content (number of adenines, ***n***) and the average read methylation, and ***β*** is a constant. Nucleosomal DNA should be associated with much lower methylation, since it is inaccessible to M.EcoGII. Accordingly, we expect that the average methylation of nucleosomal DNA will be less than the average methylation of the read, which will be less than the average methylation of accessible regions. With the null hypothesis that methylation in each 25-bp window is purely the result of random methylation efficiency, we can identify windows with lower or higher methylation than the read average using a one-tailed binomial test: For each window with ***k*** m^6^A bases, the p-value for a window to be nucleosomal is P(***X*** ≤ ***k***) and the p-value for the window to be accessible is P(***X*** ≥ ***k***). Finally, we calculated the adjusted p-value for each window within each read using the Benjamini-Hochberg procedure to minimise the family-wise error.

For data analysis using the model, each read was scanned using a 25-bp sliding window: 1) The expected local methylation in each window was calculated using Formula 1. 2) The one-tailed p-value for each window was computed to determine whether the observed methylation is higher or lower than the expected local methylation. 3) The Benjamini-Hochberg procedure was performed to calculate an adjusted p-value for the central nucleotide within each 25-bp window in each read (i.e., there is an adjusted p-value for every base pair in the read). The cumulative distribution functions of adjusted p-values for nuclei and gDNA are presented in Figure S1. Using these plots, we assigned high-confidence accessible regions using an adjusted p-value < 0.853 and high-confidence nucleosomes using an adjusted p-value < 0.918 to restrain the false-positive rate to < 1%, estimated using the gDNA control. Using these parameters, the central nucleotides in accessible windows and nucleosome windows account for only 69% (replicate 1) or 76% (replicate 2) of all nucleotides in the reads; the remainder correspond to windows with ambiguous central nucleotides (Figure S1). Most of the ambiguous nucleotides are located at the boundaries between accessible regions and nucleosomes because, as the window slides from an accessible region into a nucleosome, the m^6^A fraction decreases until it is significantly lower than random occurrence, and therefore the adjusted p-value for the central base pair in some of these windows is ambiguous. To reduce the fraction of ambiguous nucleotides, we applied the following rules: If the transition is from an accessible region to a nucleosomal region, we assigned all of the 12 nucleotides downstream of the central nucleotide in the last called accessible window as accessible (i.e., the second half of this 25-bp window); similarly, if the transition is from a nucleosome to an accessible region, we assigned all of the 12 nucleotides downstream of the central nucleotide in the last called nucleosomal window as nucleosomal. If the 12-nt extensions from nucleosomal and accessible windows overlap, the overlapping nucleotides were assigned as nucleosomal. This reduced the fraction of ambiguous nucleotides to 9% and 6% for replicates 1 and 2, respectively. Finally, any short ambiguous regions remaining that are adjacent to nucleosomal regions were assigned as nucleosomal. This re-assignment of ambiguous nucleotides as either nucleosomal or accessible reduced the fraction of ambiguous nucleotides to 7% and 4% for replicates 1 and 2, respectively. Most of the remaining ambiguous nucleotides derive from reads that have < 5% methylation. A bam file was produced to display the results of the model in IGV for visual analysis: for each molecule/read in the bam file, nucleosomes are indicated by substitutions, accessible regions are indicated as matches, ambiguous regions are labeled as deletions, and m^6^A bases are indicated as single base insertions at the corresponding adenine or thymine in the Watson strand.

### *In silico* test of the model

The median m^6^A fraction for our real data is ∼11 +/− 6%. We simulated a set of nine samples each having 100,000 25-bp windows with 60% AT content *in silico*, with per read m^6^A fractions of 5%, 10% or 15%, and nucleosomal fractions of 60%, 70% or 80%. The methylated fraction for nucleosomes was set at 1% (equivalent to 1 or 2 m^6^A within each nucleosome footprint). The m^6^A fraction in accessible regions was determined by the set values for per read m^6^A fractions and nucleosomal fractions. The test determined the fraction of windows correctly or incorrectly assigned as nucleosomal or accessible by our model. Model performance was assessed using Receiver Operating Characteristic (ROC) plots (Figure S2; Table S3).

### Nucleosome phasing relative to the TSS

To analyse the variation in nucleosome density/occupancy relative to transcription start sites (TSSs), we computed the nucleosome density for each base pair in each gene as the ratio of reads in which that base pair is nucleosomal to the read coverage (number of DNA molecules including that base pair). We selected 3,745 non-overlapping genes with a minimum gene length of 200 bp and a minimum promoter region of 200 bp for the analysis. For each gene, we computed the nucleosome density +/− 400 bp relative to the TSS for each gene, and then calculated the average nucleosome density for all 3,745 genes. TSS data were obtained from (18).

### Cell-to-cell variation in nucleosome position

We analysed cell-to-cell variation in nucleosome position by comparing nucleosome footprints on different DNA molecules belonging to the same gene, as follows: 1) Starting with the 3,745 genes identified above, we selected the 3,733 genes which had > 5 high-quality reads that include the TSS +/− 300 bp and at least one accessible region within the TSS +/− 300 bp (to exclude reads with too few m^6^A bases). 3) We extracted the 601-bp section of each read spanning the TSS +/− 300 bp region, and aligned them relative to the TSS. 4) Each 601-bp read fragment was represented as a vector of length = 601, in which accessible = 0, nucleosomal = 1 and ambiguous = 0.5, resulting in a set of vectors representing nucleosome positions for each gene in different haploid cells. 5) The correlation coefficient for every pair of vectors for each gene was calculated. For each gene, the average correlation coefficient was used as a measure of its nucleosome position heterogeneity.

## Results

M.EcoGII transfers a methyl group from SAM to the N^6^ position of adenine in double-stranded DNA (m^6^A), probably via a base-flipping mechanism (14,19,20). M.EcoGII is ideal for high-resolution methylation footprinting because of the high density of A-T base pairs in DNA and because it methylates adenines apparently regardless of their sequence context (14). We applied M.EcoGII methylation footprinting to nuclei obtained from yeast cells arrested in G1 with α-factor. Arrested cells were used to avoid potential contributions from DNA replication to accessibility. Purified nuclei were incubated with or without M.EcoGII, genomic DNA was purified and subjected to PacBio sequencing. Each experiment involved sequencing three samples: genomic DNA from M.EcoGII-treated nuclei, genomic DNA from mock-treated nuclei (negative control), and genomic DNA purified and then treated with M.EcoGII (positive control; “gDNA”) (Table S1).

### Detection of m^6^A by Single Molecule Real-Time (SMRT) sequencing

PacBio long-read SMRT sequencing provides a method for accurate detection of m^6^A in original DNA molecules (2). The sequencing platform is an array of DNA polymerases (21,22), each of which is monitored separately for the incorporation of fluorescently labeled nucleotides as it synthesizes the complementary strand of a DNA template. The fluorescence pulse begins when a nucleotide enters the polymerase active site and ends when the polymerase releases the fluorophore (23). The interpulse duration (IPD) is used to detect modified bases (24). If the polymerase encounters m^6^A in the template, the IPD is ∼5 times longer than that for unmethylated adenine, possibly because it is awaiting conversion of the favoured *cis* form of the methyl group to the *trans* form required for base pairing with the incoming thymine (24).

A SMRT sequence generated by a single polymerase pass has a large error rate (25–27). To overcome this problem, PacBio libraries are prepared by ligating hairpin adapters to the original unamplified DNA molecules, followed by denaturation to create circular single-stranded templates. These templates are sequenced continuously by strand displacement, generating a continuous very long read including multiple passes of both template strands (23,28). The long read is then split into its subreads by bioinformatic removal of intervening adapter sequences. The subreads are used to obtain a much more accurate consensus read.

### m^6^A base-calling in single DNA molecules using the subreads

We developed a pipeline designed to identify m^6^A bases in PacBio reads and then to interpret the m^6^A pattern in terms of accessible regions and nucleosome footprints at the single molecule level. The steps involved in identifying m^6^A bases are summarised in Figure 1A. The first step is to group the PacBio subreads derived from the same DNA molecule (i.e. reads from the same zero-mode waveguide (ZMW) in the PacBio device), and derive a consensus sequence for that molecule, including high-probability m^6^A bases called by the ipdSummary software.

**Figure 1.**
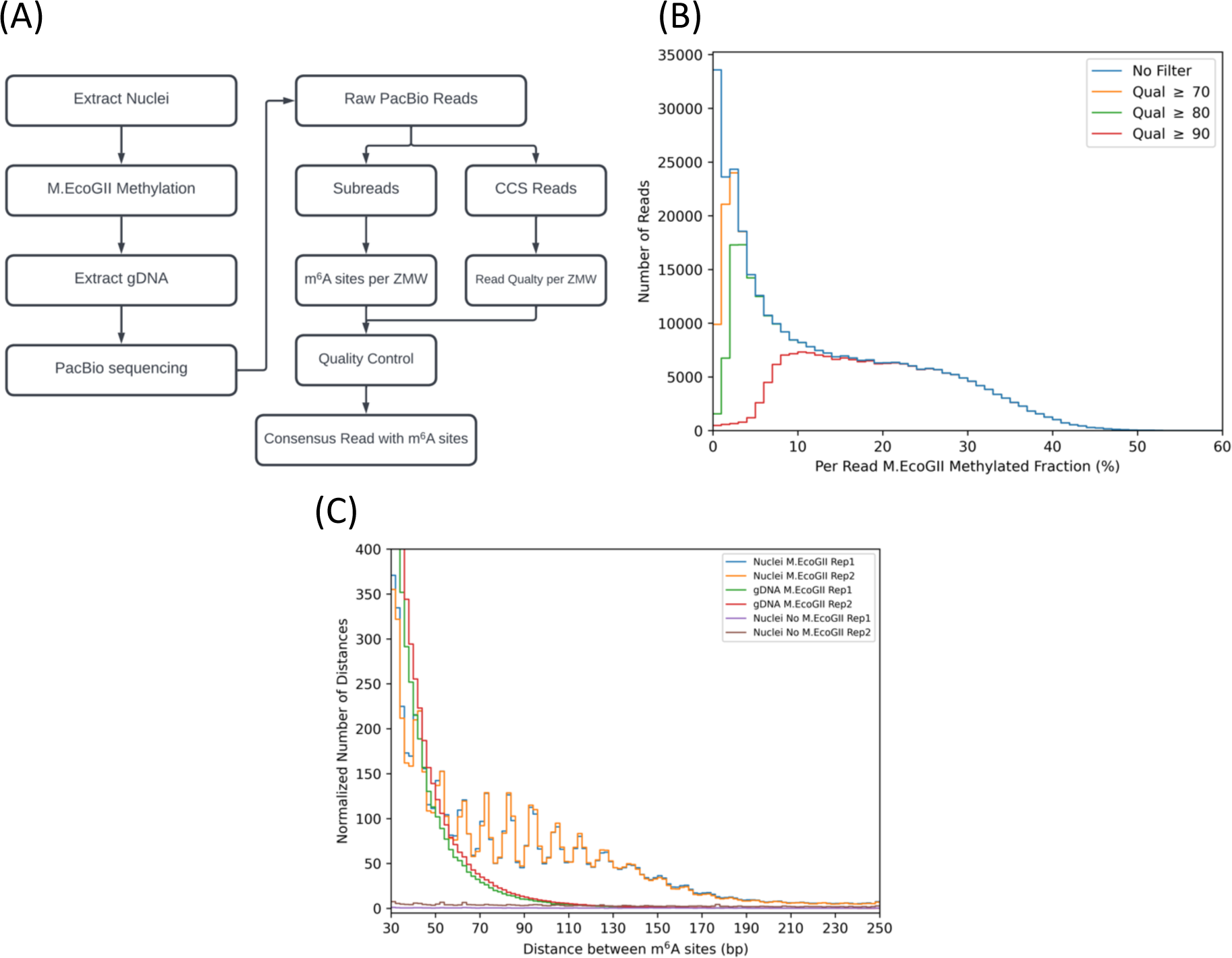
Identification of m^6^A bases in single DNA molecules. **(A)** Workflow for m^6^A identification in PacBio long reads. The zero-mode waveguide (ZMW) is a nanophotonic device for sequencing a single DNA molecule multiple times, producing repeat reads of both strands (subreads). A consensus read with accurate m^6^A base calls is obtained using the subreads. The PacBio software produces a circular consensus (CCS) read with quality scores for each base within the same DNA molecule. **(B)** Histograms of m^6^A fraction per read. Reads were filtered using different average base quality scores and the m^6^A fraction was calculated for each read. The two gDNA replicates are compared. Note that increasing the quality score effectively removes reads with low methylation. **(C)** Distribution of inter-m^6^A distances. The distance between each m^6^A and the next m^6^A was calculated for each read. The number of inter-m^6^A distances in each sample was normalised to the number of nucleotides sequenced per million. ‘Rep 1’ and ‘Rep 2’ refer to biological replicate experiments.

The next step, which is critical, is to remove poor quality consensus reads using the average quality score for each circular consensus read (CCS) generated by the PacBio pbccs protocol with default parameters (the CCS does not contain modified base information). To determine the appropriate average quality score to use as a threshold, we plotted the number of consensus reads against the fraction of methylated adenines in the consensus read for data from the combined positive control (gDNA) samples (Figure 1B). If all reads are included, a broad distribution is observed, with a high peak corresponding to reads with almost no called m^6^A bases, tailing off to a high value of ∼50% m^6^A. However, if reads with an average base quality score below 90 are filtered out, the peak of reads with little or no m^6^A is removed, revealing a broad peak ranging from ∼5% to ∼50% average read methylation. Therefore, it appears that the PacBio software cannot identify m^6^A in low-quality reads, which should, in any case, be discarded. We also confirmed that our method has a very low false-positive rate, such that the median average methylation in the negative control (mock-treated nuclei) was 0.00% for replicate 1 and 0.15% for replicate 2, using the same minimum average quality score of 90. Accordingly, we set the threshold average base quality score to 90 (Figure 1B). For the combined gDNA positive controls, the median average methylation of the high-quality reads is 19.5%, but the distribution is wide (Figure 1B), with the bottom 10% of reads having < 8.6% methylation, and the top 10% of reads having > 32.5% methylation. M.EcoGII-treated nuclei also show a wide variation in average methylation: the 10% - 90% quantile ranges from 3.5% to 25.1% with a median of 10.9% (Table S1). It is important to take this wide variation into account when interpreting the methylation pattern (see below).

A naïve approach for nucleosome identification would be to scan for m^6^A-depleted regions of about the length of DNA protected by the nucleosome (∼147 bp). We calculated the distributions of the distances between m^6^A bases in nuclei and in the gDNA positive control (Figure 1C). We found that 91% of the inter-m^6^A distances are < 30 bp in nuclei, corresponding to adenines in accessible regions. For distances > 30 bp, we observed a series of peaks with a ∼10-bp periodicity, ranging from 40 to 170 bp (Figure 1C). Such periodicity in accessibility is typical of nucleosomal DNA, suggesting that M.EcoGII occasionally methylates adenines in the minor groove on the exposed surface of the nucleosome. It should be noted that non-nucleosomal protein-DNA complexes may give a similar pattern, since 10 bp periodicity is indicative of DNA bound to a surface (29,30). We calculated that m^6^A occurs about 1.5 times on average per nucleosome footprint using the data in Figure 1C. Nevertheless, the occurrence of m^6^A within nucleosomes complicates identification of nucleosomes because the footprint is fragmented.

Unlike for nuclei, inter-m^6^A distances for gDNA show a decreasing monotonic distribution with no peaks, ranging up to ∼110 bp, suggesting that large false-positive footprints are possible (Figure 1C). These long inter-m^6^A distances in purified DNA are apparently due to low methylation levels. As the fraction of m^6^A decreases, the probability of a random gap of a given length occurring between two m^6^As increases. Since the average methylation varies widely from one DNA molecule to the next (Figure 1B), the average methylation of each read is critical for interpretation of the m^6^A pattern.

### Inefficient detection of m^6^A at the single molecule level

The median read methylation of M.EcoGII-treated and mock-treated nuclei was 9.8% and 0.0% for replicate 1, and 12.1% and 0.2% for replicate 2, respectively (Table S1). The average read methylation for the gDNA positive control replicates was 20.7% and 18.3%. Even though the gDNA samples were subjected to three rounds of methylation, they were still apparently far from completely methylated, suggesting either that M.EcoGII is unable to methylate the DNA completely, or that the ipdSummary software fails to call a large number of methylated adenines. The latter is most likely because agarose gel analysis showed that our gDNA samples were heavily digested by DpnI (which cuts GATC sites with m^6^A on both strands) and resistant to MboI (which cuts only unmethylated GATC sites), implying that the DNA is highly methylated by M.EcoGII (Figure S3). In contrast, both enzymes cut the methylated DNA from nuclei to intermediate levels, indicating that some GATC sites are fully methylated, whereas others are unmethylated - consistent with extensive protection of GATC sites within nucleosomes; DNA from the unmethylated control nuclei is resistant to DpnI and fully digested with MboI, as expected (Figure S3). Analysis of GATC site methylation using ipdSummary indicates that only ∼25% of GATC sites in the gDNA controls are methylated on both strands, and that only ∼33% of GATC sites are unmethylated (the rest are predicted as hemi-methylated which would not be cut by DpnI or MboI). Thus, the called m^6^A at GATC sites is far lower than observed by DpnI/MboI digestion, consistent with an m^6^A-calling problem.

We used plasmid DNA to determine whether the apparent low level of methylation of purified DNA reflects incomplete methylation by M.EcoGII, or failure to call m^6^A by the software (**Figure S4**). pUC19 DNA was purified from *E. coli* lacking the Dam and Dcm DNA methylases and linearised with SmaI. The DNA was then subjected to a single round of methylation by either Dam or M.EcoGII. Both Dam and M.EcoGII methylation was complete at GATC sites, as shown by complete digestion by DpnI and complete resistance to MboI (**Figure S4A**). The same samples were subjected to PacBio sequencing. We found that the predicted mean methylation of the GATC sites in pUC19 is 66+/−11% (SD) for one strand and 48+/−8% for both strands. Thus, only ∼50% digestion by DpnI is expected from m^6^A detected by PacBio sequencing (DpnI cleavage requires both strands to be methylated), which is much less than we observed (**Figure S4A**). Furthermore, even though M.EcoGII methylation results in complete digestion by DpnI, indicating that GATC sites are fully methylated, PacBio sequence analysis indicates a maximum of ∼30% methylation of each strand at GATC sites (**Figure S4B,C**). A maximum of ∼30% methylation is also observed at all other adenines (**Figure S4B,C**). We conclude that PacBio detection of m^6^A is the primary problem. It appears that m^6^A detection by PacBio is most problematic at high levels of methylation, presumably because of extensive m^6^A-induced pausing of the sequencing polymerase. Indeed, these delays result in considerably fewer subread passes for the M.EcoGII-methylated DNA than for the control and Dam-methylated pUC19 samples (**Table S1**).

Consequently, we designed our pipeline to take account of the average methylation of each molecule in order to map nucleosomes on that molecule. It is also apparent that short footprints, such as those expected from transcription factors, cannot be identified with confidence unless the methylation detection efficiency is high.

### Interpretation of the m^6^A pattern: Prediction of nucleosome positions

Given the complications described above, we opted to identify nucleosomes and M.EcoGII-accessible regions read by read. Accessible regions should be relatively highly methylated and will include the nucleosome-depleted regions (NDRs) observed at most yeast promoters (100 - 200 bp), as well as the linkers between nucleosomes, which average 15 - 20 bp in yeast. The steps in our pipeline are shown in Figure 2A. To summarise, each consensus read is scanned using a sliding 25-bp sequence window. The methylated fraction is calculated for each sequence window in the read using: [number of m^6^A in both DNA strands]/[total number of adenines (including m^6^A) in both strands]. Each window value is then compared with the average methylation of the read, given by [total m^6^A in both strands of read]/[total adenines in both strands in read]. If the window methylation is significantly higher than the average read methylation, it is called as accessible; if it is significantly lower, then it is considered protected and likely nucleosomal. This approach takes account of variation in read methylation. We discuss each step in detail below.

**Figure 2.**
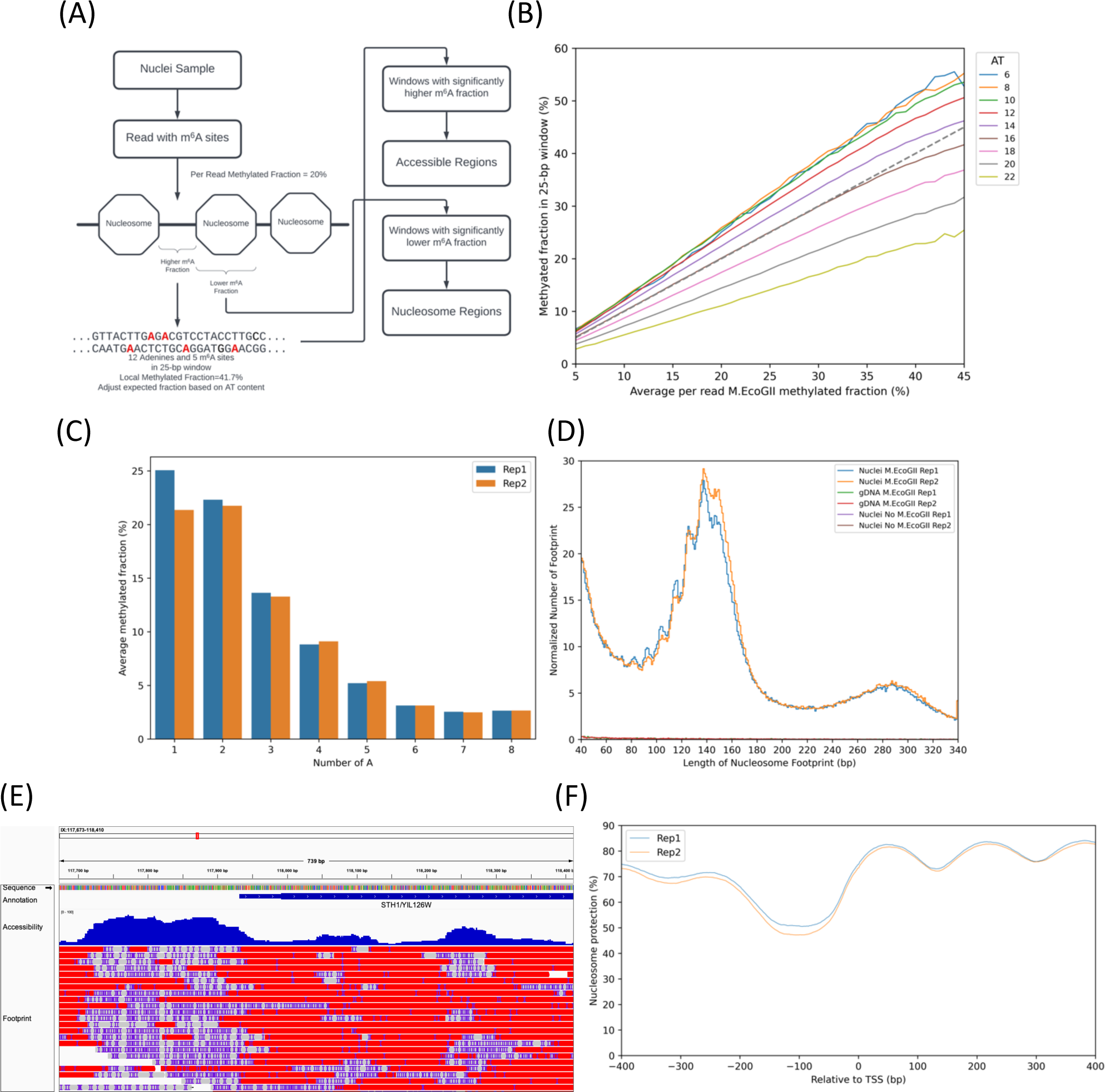
Pipeline for identifying nucleosome footprints and accessible regions in single DNA molecules. **(A)** Simplified diagram of the prediction process. The pipeline predicts accessible regions and nucleosome footprints for each read. We developed a statistical test to compare the local fraction of m^6^A in each 25 bp window with the expected m^6^A fraction based on the read average and the AT content of the window. Nucleosomal DNA should be hypo-methylated, whereas accessible regions should be hyper-methylated relative to the read average. **(B)** Linear relationship between window methylation and average read methylation as a function of window AT-content. The average methylation was calculated for each read. The 25-bp windows within each read were grouped by the number of adenines + thymines in the window sequence, and the average methylation was calculated for each group of windows with the same AT content. Data for windows from all reads with the same average read methylation were combined (rounded to 1%) and grouped by AT-content. The average methylation of each group of windows is plotted against the average read methylation. The grey dashed line corresponds to the ‘no bias’ condition, where the window average = read average. **(C)** Average fraction of m^6^A in poly(A) runs. All genomic adenines were grouped according to their location in poly(A) runs of 1 to 8 nt and the average m^6^A fraction in the gDNA samples was computed for each group. **(D)** Lengths of predicted nucleosome footprints. The major peaks at 140 - 150 bp correspond to nucleosome footprints. Some apparently fragmented nucleosome footprints appear as minor peaks < 140 bp, consistent with occasional methylation within the nucleosome, or with sub-nucleosomal particles, or other complexes (compare with Figure 1C). The number of nucleosome footprints in each sample was normalised to the number of nucleotides sequenced per million. **(E)** Snapshot of base pair level nucleosome prediction for the *YIL126W/STH1* locus in the Integrative Genomics Viewer v.2.9.2 (IGV). Called m^6^A bases are indicated as insertions (purple ‘I’), accessible (methylated) regions as matched regions (grey blocks), nucleosomes as substitutions (red blocks) and ambiguous regions as deletions (lines between blocks). **(F)** Nucleosome occupancy relative to the transcription start site (TSS) for a set of 3745 non-overlapping genes with a minimum length of 200 bp and a minimum promoter region of 200 bp. At each genome position, nucleosome protection was calculated as the percentage of reads in which base pair ‘*n*’ is nucleosomal of the total read coverage of base pair ‘*n*’.

We used the combined data from the two gDNA positive control samples to determine whether the methylation detection efficiency depends on the adenine density in a 25-bp window (Figure 2B). The dashed line indicates the condition where the window methylation is equal to the average methylation of the read. The other lines indicate data for gDNA windows with different AT contents. We observe that windows with 16 AT base pairs (64% AT) behave as expected, but GC-rich windows (< 50% AT), with only 14 or fewer AT base pairs are more methylated than expected from the read average, whereas windows with 18 to 22 AT base pairs (> 70% AT) are less methylated than expected from the read average. We conclude that methylation detection efficiency decreases with increasing AT-content. For a given AT content, the window methylation is linearly related to the average read methylation (Figure 2B). We performed linear regression for reads with an average methylation from 7% to 41% (corresponding to 5% - 99% of the reads in the gDNA samples), and obtained good linear relationships for all 25-bp windows with between 5 and 23 AT base pairs (*R^2^* > 0.99; Table S2). Since these windows account for 99.9% of the yeast genome, this linear relationship works well for almost the entire genome. In a related analysis, we found that runs of adenines on the same strand (poly(A)) are progressively less methylated as the length of the poly(A) run increases (Figure 2C). The increasing bias against m^6^A calling for sequences with higher AT-content is quite strong and should be accounted for when interpreting the local methylation pattern.

Using an adjusted binomial model based on the linear relationship between window methylation and average read methylation (Figure 2B, Table S2), we calculated the expected window methylation for the observed average read methylation to interpret the methylation pattern of each read in turn. For nuclei, methylation of nucleosomal DNA should be low relative to the calculated expected methylation, whereas methylation of accessible regions should be high relative to the calculated expected methylation (see Materials and Methods). Adjusted p-values were calculated to determine whether a window should be designated nucleosomal or accessible: threshold adjusted p-values of 0.853 and 0.918 were chosen for accessible regions and nucleosomes, respectively, to limit the false-positive rate to < 1%, estimated using the gDNA control (Figure S1).

We tested our model by simulating some simplified data with different methylated fractions and different nucleosome densities. We find that the area under the curve (AUC) values are excellent for 10% and 15% m^6^A, but not so good for 5% m^6^A (Figure S2). Therefore reads with unusually low m^6^A (5% or less) are not predicted as accurately by our model (Table S3).

Comparison of a plot of the footprint lengths predicted by our model (Figure 2D) with a similar plot of the actual inter-m^6^A distances (Figure 1C) shows that our model eliminates almost all of the false positive footprints predicted by inter-m^6^A distances for gDNA and that, in nuclei, the model also eliminates almost all of the shorter footprints > 30 bp predicted by inter-m^6^A distances. The remaining short footprints in nuclei may represent real footprints due to protection by transcription factors or other DNA-bound proteins, or residual false positive footprints. This analysis demonstrates that our pipeline resolves the problem of limited m^6^A detection efficiency. Furthermore, a major nucleosomal peak is now apparent, with sub-peaks at ∼116, 125, 137 and 146 bp, similar to the peaks observed using MNase-Exo-seq (31). Since the choice of a 25-bp sliding window was somewhat arbitrary, we tested the effect of sliding window size on predicted footprint length and predicted nucleosomal fraction (**Figure S5**): window sizes of 15, 20, 25 or 30 bp gave very similar results, indicating that sliding window size has only minor effects within the range 15-30 bp.

The output of our pipeline is an annotated bam file showing each read aligned to the yeast genome, with m^6^A bases marked, together with nucleosome positions and accessible regions predicted by our model (an example is shown in Figure 2E). Visualizing this bam file with software like IGV allows visual exploration of heterogeneity at the single molecule level and base pair resolution across the genome. The output for the whole genome is provided as processed data files at GEO.

### Cell population average nucleosome density at promoters

To confirm that our model also predicts the expected population average observed using MNase (32), we plotted the nucleosome occupancy obtained from combining the data from all reads after alignment to the transcription start site (TSSs) for a large subset of genes (3,745 genes out of 5,770) (Figure 2F). This nucleosome phasing plot shows the expected peaks corresponding to the −1, +1, +2 and +3 nucleosomes, with the expected promoter NDR between the −1 and +1 nucleosomes. The plot also provides a measure of absolute nucleosome occupancy on gene bodies in nuclei, averaging 81% for replicate 1 and 80% for replicate 2, similar to recent measurements using other methods (17,33).

### Heterogeneous nucleosome positioning around promoters

We selected 3,733 genes with more than 5 reads that include the entire 601-bp region centered on the TSS (i.e. TSS +/− 300 bp). We extracted the read fragments spanning this region for each gene, and converted the methylation pattern of each read fragment to a vector in which accessible regions were assigned a value of 0, nucleosomes were assigned a value of 1, and ambiguous regions were assigned a value of 0.5. We calculated the correlation coefficient between each pair of reads from the same gene (see Materials and Methods). The average correlation coefficient for all read pairs from the same gene was used as a measure of the heterogeneity of nucleosome positioning, such that lower values indicate increased heterogeneity. The average correlation coefficients for all 3,733 genes are listed in Table S4 and presented as a histogram in Figure 3A. Only 9 genes have an average correlation coefficient > 0.4, corresponding to relatively homogeneous nucleosome positioning, whereas 1,115 of the 3,733 genes have an average coefficient < 0.1, indicating extreme heterogeneity.

**Figure 3.**
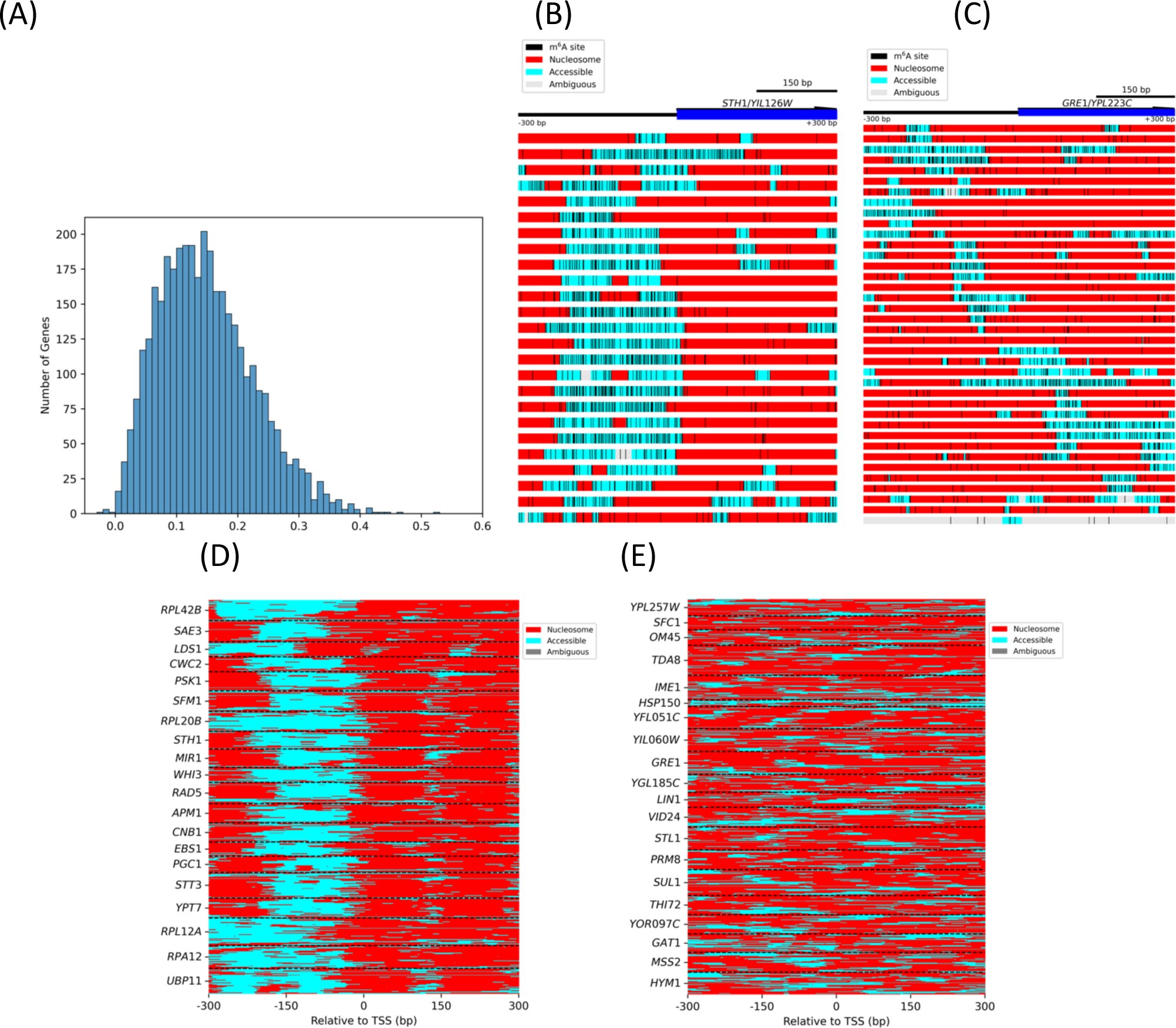
Heterogeneity in nucleosome positioning. **(A)** Histogram of correlations in nucleosome positioning on individual genes. We selected 3,733 genes with at least 300-bp separation between the gene and the next gene upstream and with at least 6 reads overlapping the TSS +/− 300 bp region. For each gene, we calculated the correlation coefficient for nucleosome positions in all the reads in the +/− 300 bp region (decreasing correlation indicates increasing heterogeneity in nucleosome positioning; see Methods). **(B)** Nucleosome positioning at the *YIL126W/STH1* locus. An example of relatively homogeneous positioning. Each block represents one read. The basepairs predicted to occupy accessible regions are indicated as cyan blocks, nucleosomes as red blocks, and ambiguous regions as grey blocks. m^6^A bases are indicated by black vertical lines. **(C)** Nucleosome positioning at the *YPL223C/GRE1* locus. This locus illustrates heterogeneous positioning. **(D)** The top 20 genes with homogeneous nucleosome positioning. Each line represents a single read, annotated as above. Reads belonging to different genes are separated by dashed lines. All of these genes have obvious promoter NDRs. **(E)** The top 20 genes with heterogeneous nucleosome positioning. These genes do not have accessible promoters and nucleosome positions vary widely from molecule to molecule.

The *YIL126W/STH1* gene is an example of relatively homogeneous nucleosome positioning (average *R* = 0.42; Figure 3B). This gene has a well-formed NDR at its promoter, and 9 of the 25 read fragments have well-positioned +1 and +2 nucleosomes clearly separated by a methylated linker. Twelve of the 25 read fragments exhibit protected regions much larger than a nucleosome, which may correspond to +1/+2 dinucleosomes, since there is no linker in between (34–36). The *YPL223C/GRE1* gene is an example of extremely heterogeneous nucleosome positioning (Figure 3C; average *R* = −0.002; Table S4); the 38 read fragments differ markedly from one another (only 2 read fragments show a typical NDR and a linker between the +1 and +2 nucleosomes). The nucleosome positions are also very different from read to read, often with poorly defined linkers.

More generally, the twenty genes with the most homogeneous nucleosome positioning and the twenty genes with the most heterogeneous positioning are compared in Figure 3D and Figure 3E, respectively (and shown in detail in Figures S6 and S7). The homogeneous genes generally exhibit clear promoter NDRs with well defined +1 and +2 nucleosomes downstream of their promoters; many read fragments from those genes show nucleosome positions shifted relative to one another (Figure 3D). In contrast, the heterogeneous genes do not have an obvious promoter NDR, and nucleosome positioning appears random with respect to the TSS (Figure 3E), consistent with a role for the NDR in organising phased nucleosomal arrays (reviewed by (37)).

In conclusion, our analysis demonstrates that nucleosome positioning is generally heterogeneous at the single gene level. This is also true at the single cell level, since each read is derived from a single cell. This is the case even though the cell population average indicates that nucleosomes form phased arrays relative to the TSS, as expected from MNase-seq data (Figure 2F).

## Discussion

We have presented a bioinformatic pipeline that utilises a probability model to resolve interpretative problems associated with PacBio long-read methylation footprinting experiments. Our M.EcoGII footprinting data illustrate several challenges that we had to address before the data could be interpreted in terms of nucleosomes and accessible regions: apparent relatively low methylation levels in the genomic DNA positive control samples, a wide range in the fraction of m^6^A from one DNA molecule to the next, and a strong local bias against methylation of AT-rich sequences and poly(A) runs. Furthermore, in nuclei, there is a high probability of observing a single m^6^A base within a nucleosome, breaking up the expected ∼147 bp footprint.

Despite enzymatic detection of m^6^A indicating near complete methylation of protein-free DNA by M.EcoGII, the median methylation level called using PacBio sequence data did not exceed 27%. We used an older PacBio Sequel I machine for the yeast studies, but we observe the same issue with our pUC19 data, which were obtained using a Sequel IIe machine. Another study (2) using Sequel II also reported relatively low % methylation per PacBio read in their *Drosophila* samples (gDNA: 16% m^6^A, ranging from 6 to 27%; nuclei: 11%, ranging from 3 to 21%). These observations suggest that M.EcoGII might be unable to methylate all adenines in a DNA molecule. However, a previous study reported almost complete methylation of DNA by M.EcoGII, using liquid chromatography and mass spectrometry to measure the m^6^A/A ratio in M.EcoGII-treated DNA hydrolysed to its constituent nucleosides (14). We demonstrated that the GATC sites in pUC19 can be completely methylated by M.EcoGII, and that ipdSummary fails to detect most of the m^6^A in fully methylated pUC19. We observed that PacBio sequencing is much more efficient in detecting methylated GATC sites in a background of unmethylated DNA, suggesting that high levels of flanking m^6^A interfere with m^6^A-calling (accounting for the reduced detection efficiency for m^6^A in poly(A) sequences), perhaps because cumulative local delays confound the ipdSummary algorithm. Inefficient m^6^A base calling may also reflect selection for high-confidence m^6^A in ipdSummary, ignoring low-confidence m^6^A calls, some of which may be true positive calls. Most importantly, our pipeline effectively compensates for low m^6^A-calling efficiency.

To correct for m6A-calling efficiency and determine the accessibility at each base pair, our pipeline utilises a probability model to compare the local m^6^A level in 25-bp sliding windows along each DNA molecule with that expected from the average m^6^A for the entire molecule. Relatively high m^6^A fractions indicate accessible regions, whereas relatively low m^6^A fractions indicate protected regions, presumed to be nucleosomes. This per read approach adjusts for the wide range of methylation of individual molecules within the same sample. The probability model also takes account of the observed bias against methylation in local windows of higher AT-content and longer poly(A) runs. The occasional m^6^A within a nucleosomal footprint, which in a naive approach would break up its footprint, is ignored by the model, because the local m^6^A fraction is still below the read threshold for calling an accessible region. We note that short footprints, potentially attributable to transcription factors or other DNA-bound proteins, will be unreliable until m^6^A base calling is improved.

We used our long-read sequence data to investigate heterogeneity in chromatin structure around the TSSs of yeast genes. In contrast to the phased nucleosome arrays observed downstream of the TSS for the average gene at the population level, we find that nucleosome positioning on a specific gene in individual cells is generally highly variable, with only a small fraction of genes showing a similar ordered nucleosomal array in each cell. Though perhaps surprising, this degree of heterogeneity is consistent with single-gene MNase studies using monomer extension (16,38,39) and with genome-wide MNase-seq studies (40), both of which show multiple overlapping alternative nucleosome positions for each nucleosome in the array, which can only be accounted for by cell-to-cell heterogeneity (sometimes called “fuzzy” positioning). It seems likely that heterogeneity reflects chromatin dynamics, as nucleosomes are shifted or displaced by ATP-dependent chromatin remodelers and polymerases during transcription, DNA replication and repair (37). The nucleosome spacing enzymes may play an important role in generating heterogeneity, because nucleosomes are spaced with different linker lengths, according to the (10*n* +5) bp rule (41), resulting in alternative positions, and because of interference between nucleosome arrays, depending on the distance between phasing barriers (42).

In pioneering work, Nanopore long-read sequencing has been used to map chromatin structure in yeast (1,4). Wang et al. (4) treated yeast spheroplasts (which presumably lysed in the reaction buffer) with M.CviPI, which methylates C in GC sequences, to obtain detailed chromatin maps, although their resolution is limited by the genomic distribution of GC sequences. To maximise resolution, Shipony et al. (1) treated yeast nuclei with three different DNA methylases (M.EcoGII, M.SssI and M.CviPI) and then detected both the m^6^A and m^5^C bases in each DNA molecule (1). They used the Nanopore modified base caller to calculate the methylation probability for a fixed window using a Bayesian probability model to overcome the noisy signal. They reported some unmappable regions, some fully methylated reads and, surprisingly, high methylation at the nucleosome dyad. However, we did not observe any of these in our nuclei samples. These issues may reflect the difficulty of calling both m^6^A and m^5^C in Nanopore sequences. Generally, Nanopore methylation calls may be less accurate than PacBio because the template molecule is sequenced only once as it passes through the pore, whereas PacBio sequences are derived from multiple reads of the same molecule.

In principle, the use of PacBio sequencing to detect m^6^A introduced by M.EcoGII or similar enzymes improves the resolution of methylation footprinting still further, although inefficient m^6^A-calling limits the actual resolution at this point in time. In an elegant study (2), PacBio sequencing was used to detect m^6^A in *Drosophila* and human K562 DNA from nuclei treated with Hia5, an enzyme similar to M.EcoGII. NDRs and linkers were defined by long and short m^6^A clusters, respectively; nucleosomes were then defined by the unmethylated regions between these clusters, which ranged from 80 - 250 bp in length. However, since yeast linkers (15 - 20 bp) are shorter than those of *Drosophila* (30 - 45 bp), simple detection of m^6^A clusters to define linkers does not work well for yeast.

We believe that our approach and pipeline represent a significant improvement on earlier methods. We note that new computational pipelines are being developed using neural net approaches (43) (https://github.com/fiberseq/fibertools-rs and https://github.com/RamaniLab/SAMOSA-ChAAT). Our pipeline should be useful for interpreting other PacBio methylation footprinting studies. The bam file output is convenient for use in the IGV viewer, visualising both called m^6^A bases and interpretation of the methylation pattern as accessible regions and nucleosomes in individual DNA molecules.

## Data availability

The PacBio sequence data and processed files for viewing the data are available at the GEO database under accession number GSE243114. The code for m^6^A identification from raw subreads and the prediction of nucleosome locations is available at Github and FigShare.

Nucleosome Prediction: https://github.com/zhuweix/AdenineFootprinter

m6A prediction: https://github.com/zhuweix/MethyladenosineFinder

**Reviewer links:** To review GEO accession GSE243114:

Go to https://www.ncbi.nlm.nih.gov/geo/query/acc.cgi?acc=GSE243114

Enter token uvwjcukujrovton into the box.

m6A prediction (MethyladenosineFinder)

DOI (The DOI becomes active when the item is published.)

10.6084/m9.figshare.22194700

Private link (Note: Do not reference this link in papers. Use the public DOI.)

https://figshare.com/s/9a192db8187c465a914b

Footprint prediction (AdenineFootprinter)

DOI (The DOI becomes active when the item is published.)

10.6084/m9.figshare.22194712

Private link (Note: Do not reference this link in papers. Use the public DOI.)

https://figshare.com/s/6e19a2225094ebff4bda

## Acknowledgements

We thank the NCI CCR Sequencing Facility and the NICHD Molecular Genomics Core for PacBio library preparation and sequencing (Tianwei Li and James Iben). We thank Alan Hinnebusch, Michael Lichten and Carl Wu for much helpful discussion. We thank Peter Eriksson for useful comments on the manuscript. This study utilised the high-performance computational capabilities of the Biowulf Linux cluster at the National Institutes of Health (NIH).

