## Supplementary Figures S1-S7 for "Examining chromatin heterogeneity through PacBio long-read sequencing of M.EcoGII methylated genomes: an m^6^A detection efficiency and calling bias correcting pipeline"

### Examining chromatin heterogeneity through PacBio long-read sequencing of M.EcoGII methylated genomes: an m$^6$A detection efficiency and calling bias correcting pipeline

Allison F. Dennis, Zhuwei Xu and David J. Clark

Division of Developmental Biology,
*Eunice Kennedy-Shriver* National Institute of Child Health and Human Development,
National Institutes of Health,
Bethesda MD 20892, USA.

(A) (B)

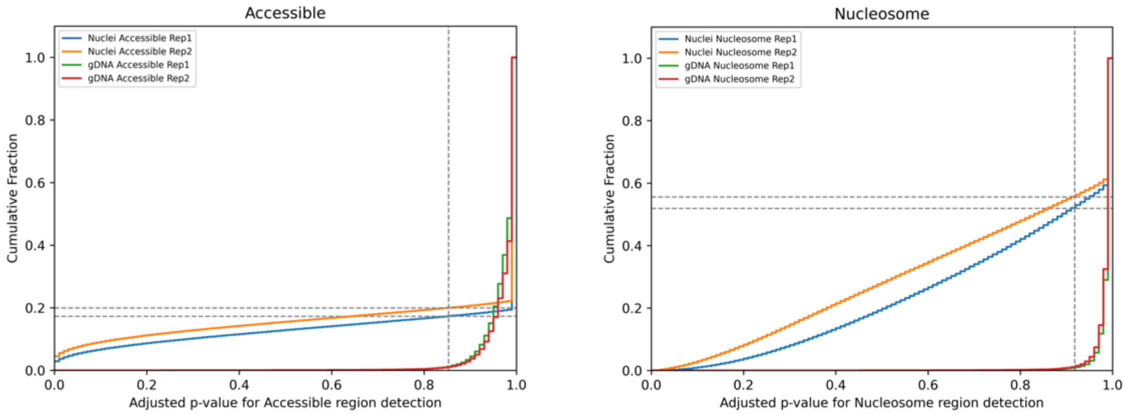

**Figure S1.** Selection of adjusted p-value for high-confidence accessible regions and nucleosome regions. Using our model, we computed the cumulative distribution function (CDF) of the fraction of each read predicted as **(A)** accessible or **(B)** nucleosomal, as a function of the threshold adjusted p-value obtained after the Benjamini-Hochberg procedure. The CDF for the gDNA positive control reflects the false-positive rate in our analysis, while the CDF for M.EcoGII-treated nuclei estimates the fraction of the genome with high-confidence accessible regions or nucleosomes at various threshold adjusted p-values. The adjusted p-value cut-off is indicated by a vertical dashed line, and the corresponding fractions of high-quality accessible/nucleosomal regions are indicated by horizontal dashed lines. For accessible regions, an adjusted p-value = 0.853 was selected to ensure that the false positive rate (CDF in gDNA samples) is < 1%; we expect high-confidence accessible regions to account for 17% and 20% of the genome in the two replicates, respectively. For nucleosomes, an adjusted p-value = 0.918 was selected to ensure the false positive rate (CDF in gDNA samples) is < 1%; we expect high-confidence nucleosomes to occupy 52% and 56% of the genome in the two replicates, respectively. Therefore, 31% of 25-bp windows in replicate 1 and 24% of windows in replicate 2 have an ambiguous central nucleotide (i.e., not assigned to an accessible region or a nucleosome). We reduced the fraction of ambiguous central nucleotides to 7% and 4% of all nucleotides in the reads for replicates 1 and 2, respectively (see Materials and Methods for details).

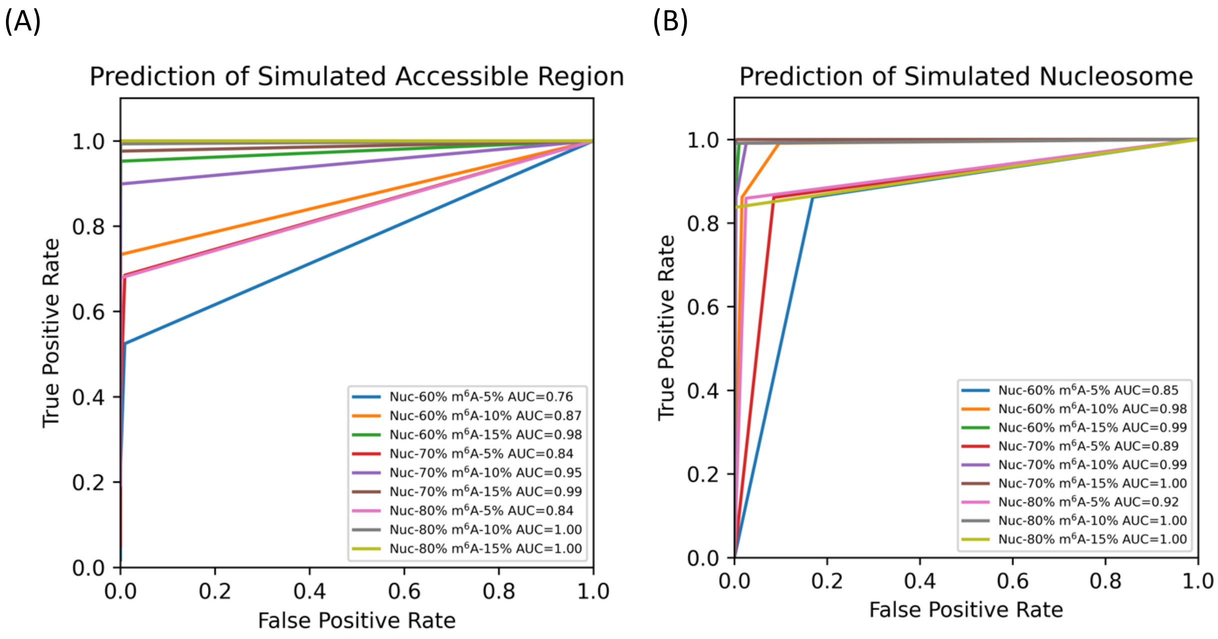

**Figure S2.** *In silico* test of our nucleosome footprint prediction model. The experimental data for nuclei have a median $m^6A$ fraction per read of ~11% (standard deviation ~6%). We tested our footprint detection model *in silico* using a range of average $m^6A$ fractions per read (5, 10 and 15%) and a range of nucleosome-protected fractions (60, 70 and 80% nucleosomal) using simulated data. The methylated fraction for nucleosomes was set at 1% (equivalent to 1 or 2 $m^6A$ within each nucleosome footprint). The $m^6A$ fraction in accessible regions was determined by the set values for $m^6A$ fractions per read and nucleosomal fractions. For each simulation, 100,000 25-bp windows with 60% AT were generated. Model performance was assessed using ROC plots. The test determined the fraction of windows correctly or incorrectly assigned by our model as (**A**) accessible regions, or (**B**) nucleosomal regions. Each point in the ROC curve indicates the true positive rate (TPR) and false positive rate (FPR) for a given adjusted p-value. The test shows good ROC curves for the detection of accessible regions in reads with 10% and 15% $m^6A$. The fraction of false-positive windows is generally low, but true-positive windows may be incorrectly assigned in reads with low $m^6A$ (5%).

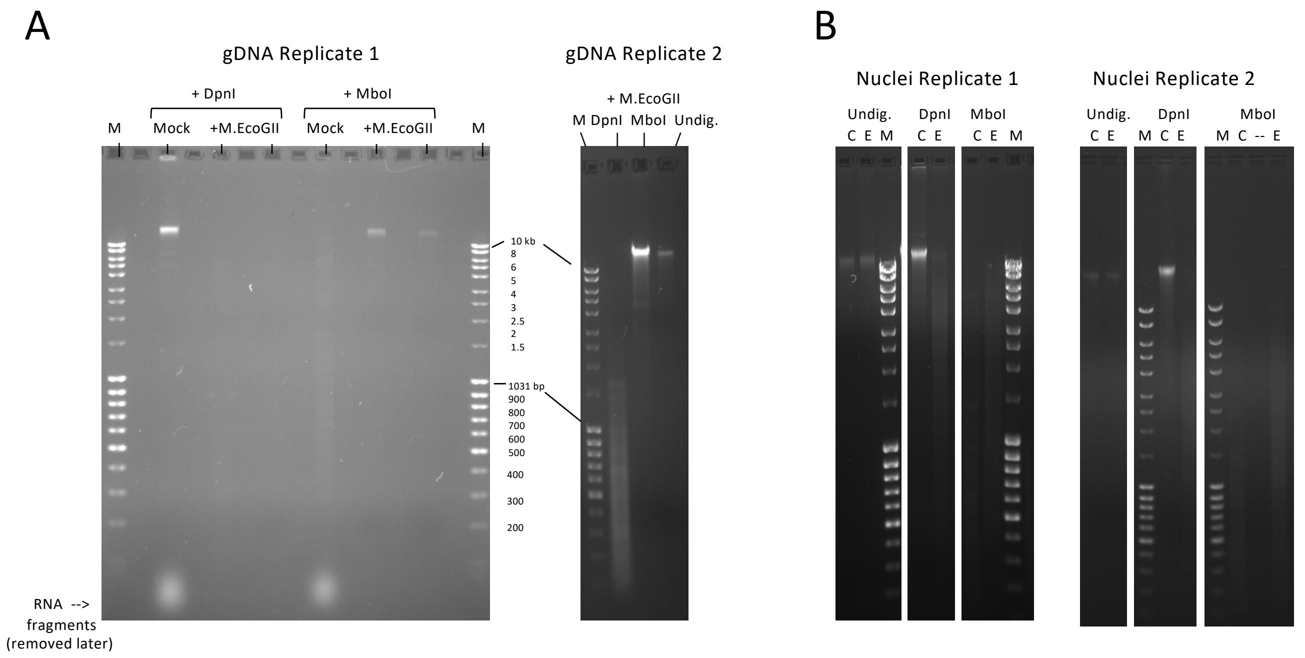

**Figure S3.** Estimation of the extent of methylation at GATC sites: Test digests with DpnI and MboI. DpnI cuts at GATC only if the 'A' on both strands is methylated; MboI cuts at GATC only if the 'A' on both strands is unmethylated. M.EcoGII-treated and mock-treated samples (125 ng) were digested for 1 h at 37°C in CutSmart buffer (NEB) with DpnI (10 units; NEB) or MboI (2.5 units; NEB) as indicated and analysed by electrophoresis in agarose gels containing ethidium bromide. 'Undig.': undigested DNA. Marker, 'M': MassRuler DNA Ladder Mix (Thermo-Fisher SM403). **(A)** gDNA replicates. Purified genomic DNA was methylated in vitro by M.EcoGII, purified and then digested with DpnI or MboI. (**B**) Nuclei replicates. Nuclei were methylated with M.EcoGII ('E') or mock-methylated (control 'C'), the DNA was purified, and digested with DpnI or MboI. Note: the lanes shown for Replicate 1 all come from the same gel (additional sample lanes, represented by the gaps, were removed for clarity); the same is true for Replicate 2.

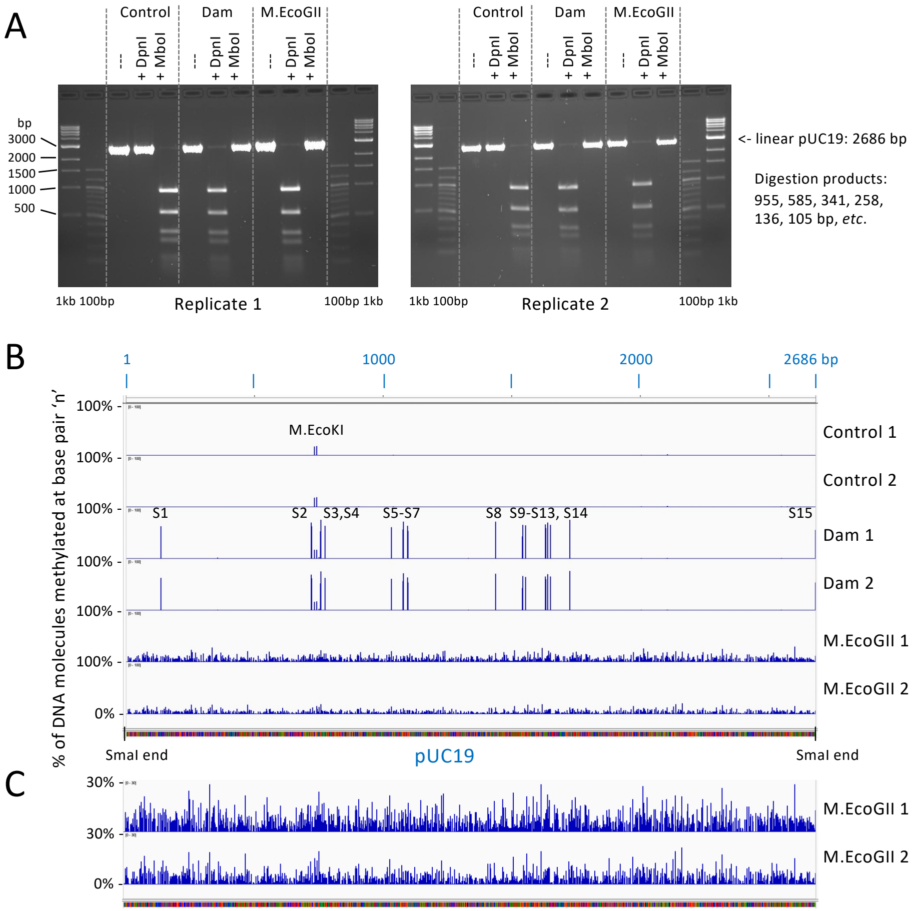

**Figure S4.** Detection of m6A in plasmid pUC19 methylated *in vitro* using Dam or M.EcoGII. pUC19 was purified from an *E. coli dam- dcm-* strain lacking the Dam and Dcm DNA methylases, but does have M.EcoKI, which methylates adenine in the sequence 'GCAC-$N_6$-GTT' (one site in pUC19). pUC19 linearised with SmaI was methylated by Dam or M.EcoGII. **(A)** Agarose gel analysis of DpnI and MboI digests of unmethylated pUC19 (control) and pUC19 methylated by Dam or M.EcoGII. DpnI cuts only if both strands of a GATC site are methylated; MboI cuts only if neither strand in a GATC site is methylated. Markers: NEB 1-kb and 100-bp ladders. **(B)** Detection of m6A in the same pUC19 samples by PacBio long-read sequencing (IGV viewer). Dam methylation (GATC) sites: S1 to S15. The population average methylation for each 'A' and 'T' is shown. M.EcoKI: adenine methylation at the 'A' and the penultimate 'T' in 'GCAC-$N_6$-GTT'. **(C)** The M.EcoGII data from 'B' after adjustment to a range of 0-30% average methylation.

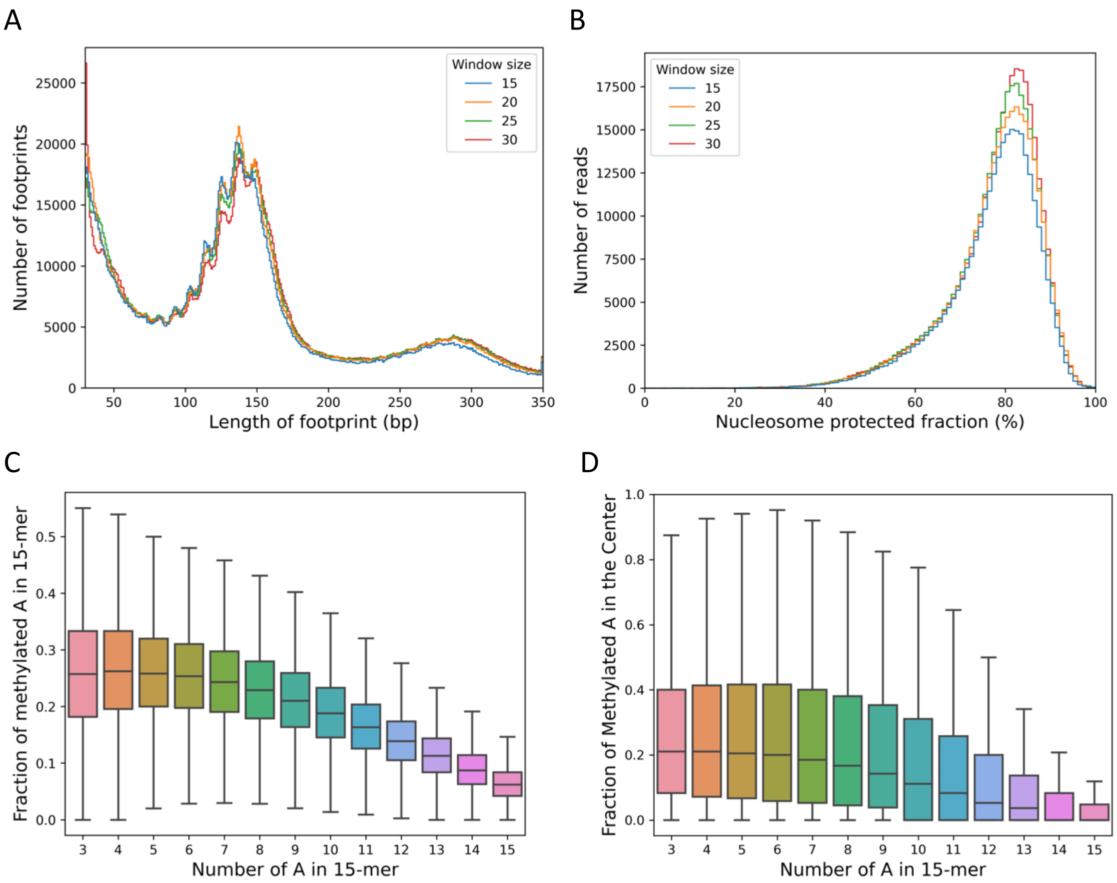

**Figure S5.** Supplementary test of the model. **(A) The impact of sliding window size on footprint length.** We ran our pipeline using different sliding window sizes (15, 20, 25 or 30 bp), and calculated the length distribution of the detected footprints in the M.EcoGII methylated nuclei sample for the combined replicates. **(B) The impact of sliding window size on nucleosome protection.** We performed the same test as in (A) and calculated the fraction of nucleosomal regions within each read as the total length of footprints within one read / read length. **(C) Effect of AT content on m$^6$A detection in 15-mers in M.EcoGII-treated gDNA.** For each 15-mer with 3-15 adenines (counting both strands), we calculated the average fraction of adenines in both strands that are methylated in the combined replicates. We grouped the 15-mers by number of As. The figure shows a decrease in m$^6$A as AT content increases. **(D) Effect of AT content on detection of methylation of the central adenine in 15-mers in M.EcoGII-treated gDNA.** We performed a similar analysis to that in (C), but examined only the fraction of methylation calls for the central adenine in each 15-mer.

### Supplementary Figure 6. Nucleosome positioning on the 20 most homogeneous genes.

Nucleosome positioning on the 20 most homogeneous genes. The genes are ordered by homogeneity. Each block below the gene locus represents one read. Accessible regions are indicated by cyan boxes, nucleosomes as red boxes, and ambiguous regions by grey boxes. Called $m^6A$ bases are indicated by vertical black lines.

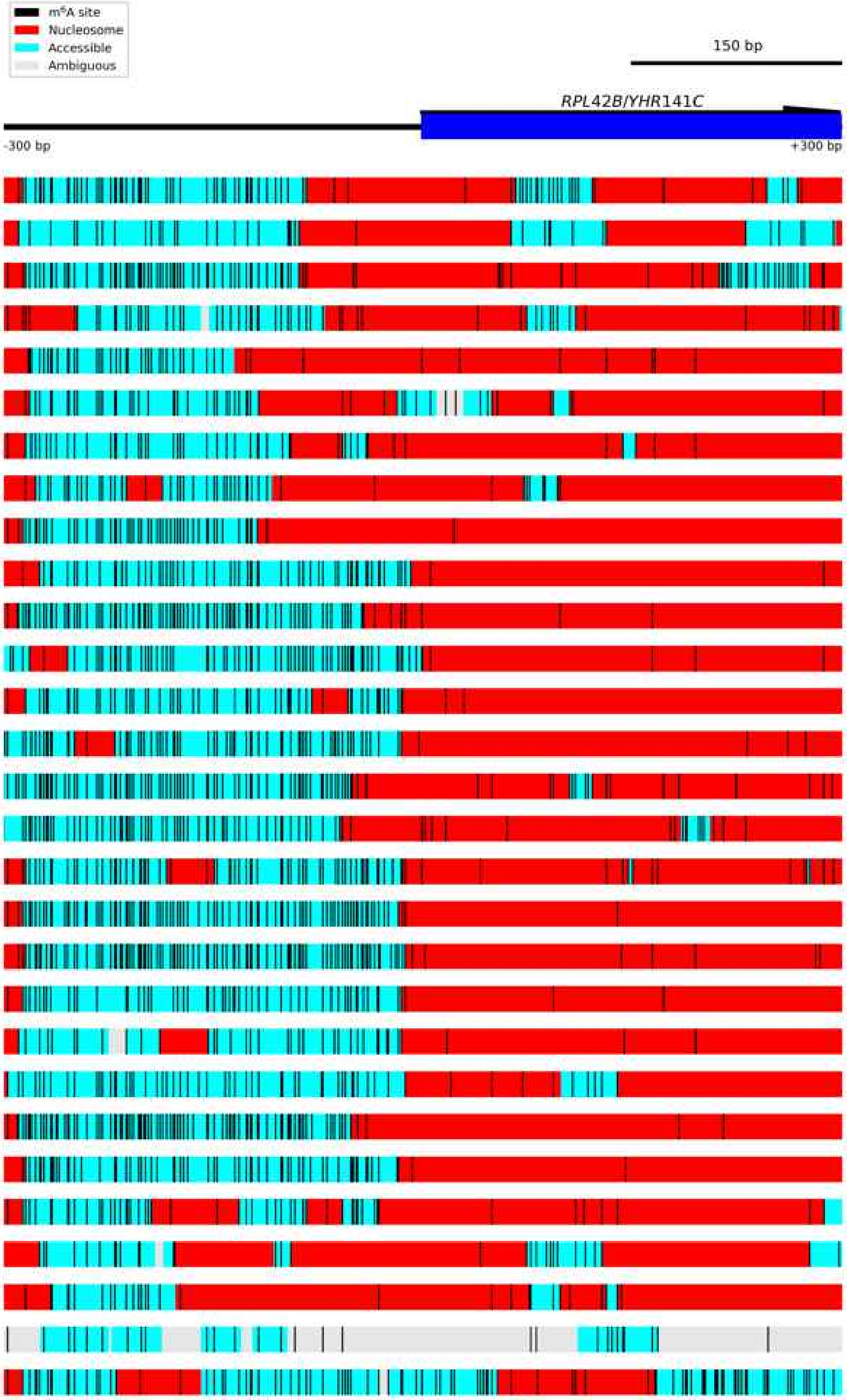

Figure S6.1 **Diagram of *RPL42B/YHR141C*.**

The *RPL42B/YHR141C* locus was an example of homogeneous nucleosome positioning. Each block below the gene locus represents one read. Accessible regions are indicated by cyan boxes, nucleosomes as red boxes, and ambiguous regions by grey boxes. Called $m^6A$ bases are indicated by vertical black lines.

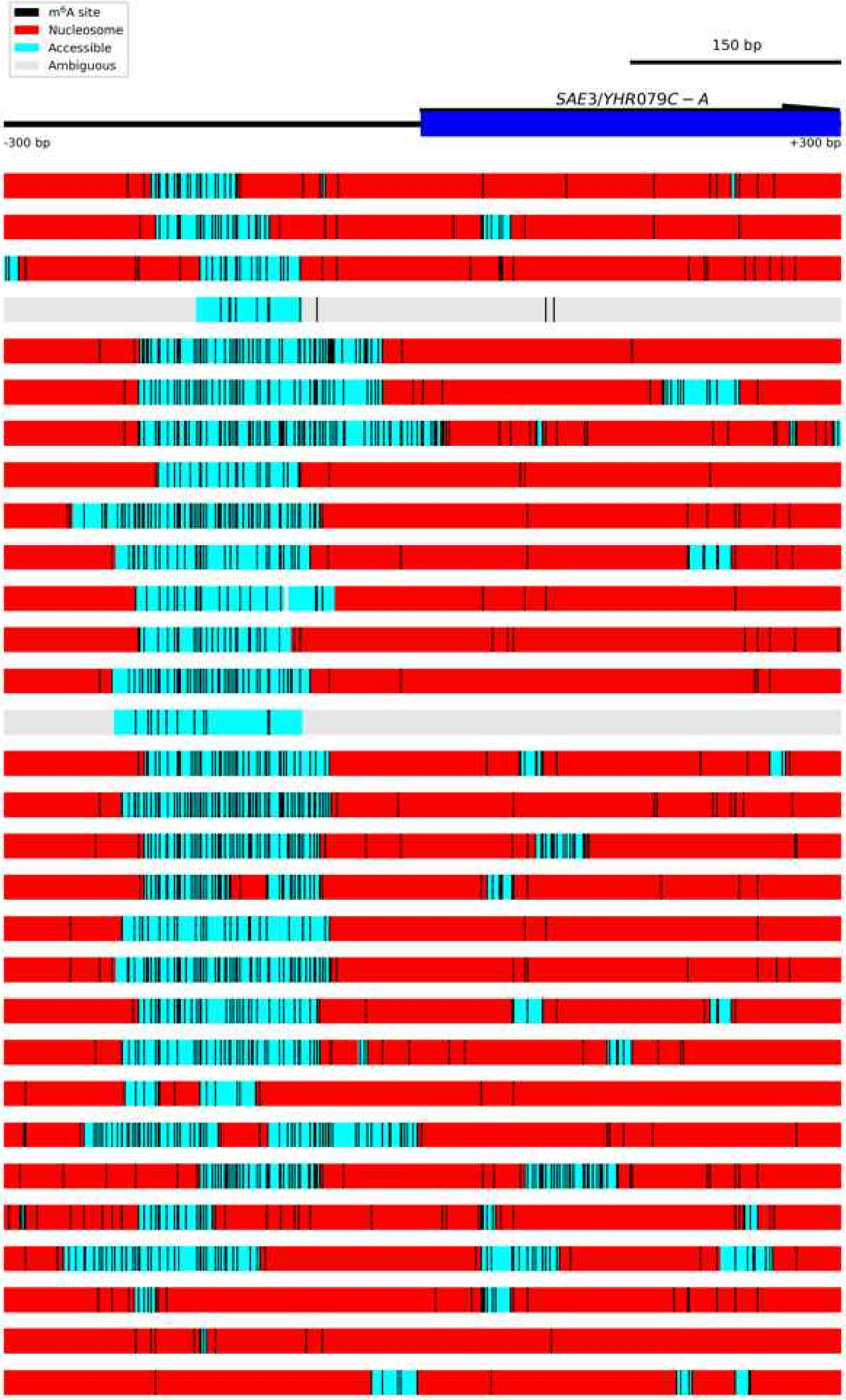

Figure S6.2 **Diagram of *SAE3/YHR079C-A*.**

The *SAE3/YHR079C-A* locus was an example of homogeneous nucleosome positioning. Each block below the gene locus represents one read. Accessible regions are indicated by cyan boxes, nucleosomes as red boxes, and ambiguous regions by grey boxes. Called m$^6$A bases are indicated by vertical black lines.

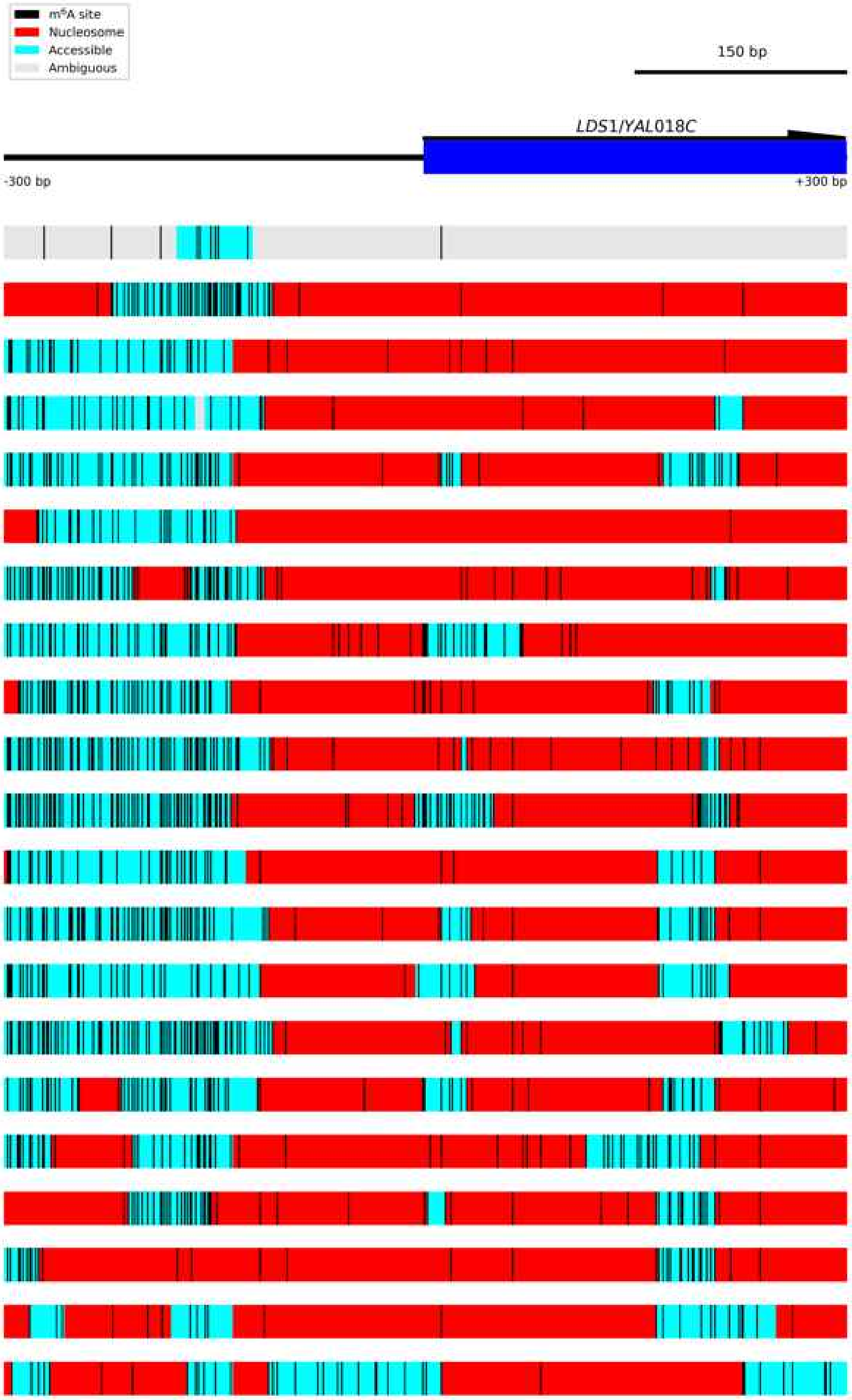

Figure S6.3 **Diagram of *LDS1/YAL018C*.**
The *LDS1/YAL018C* locus was an example of homogeneous nucleosome positioning. Each block below the gene locus represents one read. Accessible regions are indicated by cyan boxes, nucleosomes as red boxes, and ambiguous regions by grey boxes. Called m$^6$A bases are indicated by vertical black lines.

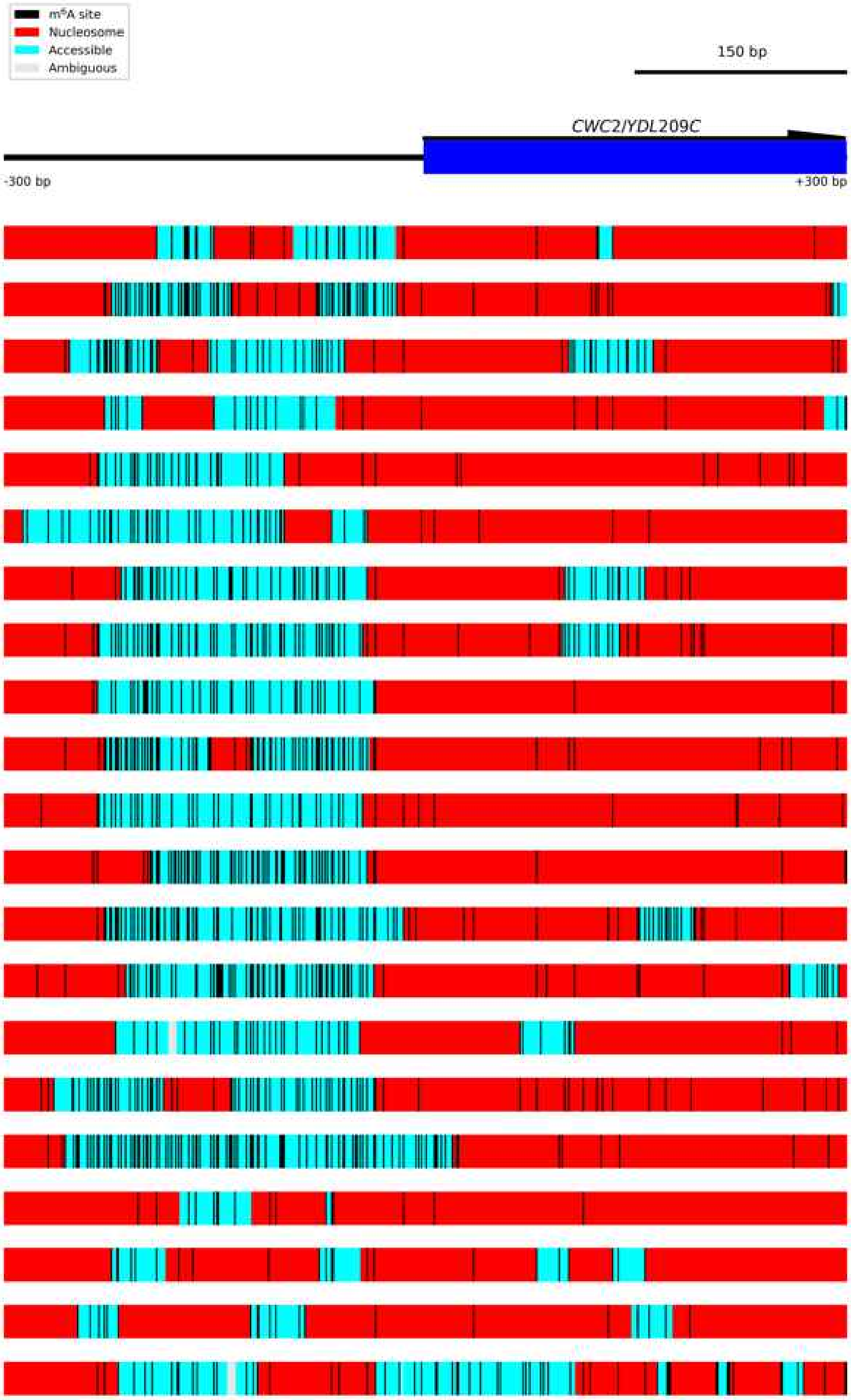

Figure S6.4 **Diagram of *CWC2/YDL209C*.**
The *CWC2/YDL209C* locus was an example of homogeneous nucleosome positioning. Each block below the gene locus represents one read. Accessible regions are indicated by cyan boxes, nucleosomes as red boxes, and ambiguous regions by grey boxes. Called $m^6A$ bases are indicated by vertical black lines.

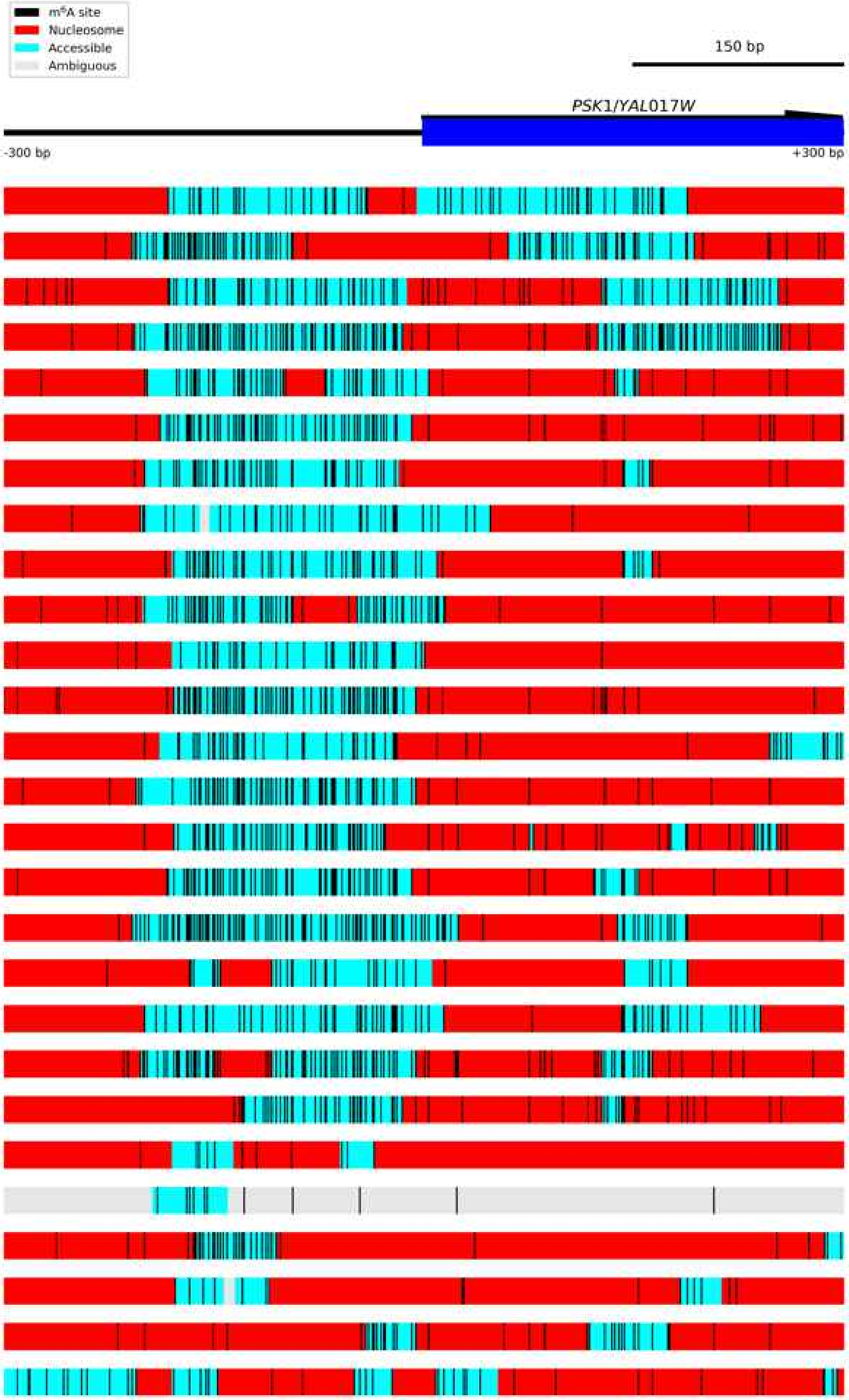

Figure S6.5 **Diagram of *PSK1/YAL017W*.**
The *PSK1/YAL017W* locus was an example of homogeneous nucleosome positioning. Each block below the gene locus represents one read. Accessible regions are indicated by cyan boxes, nucleosomes as red boxes, and ambiguous regions by grey boxes. Called $m^6A$ bases are indicated by vertical black lines.

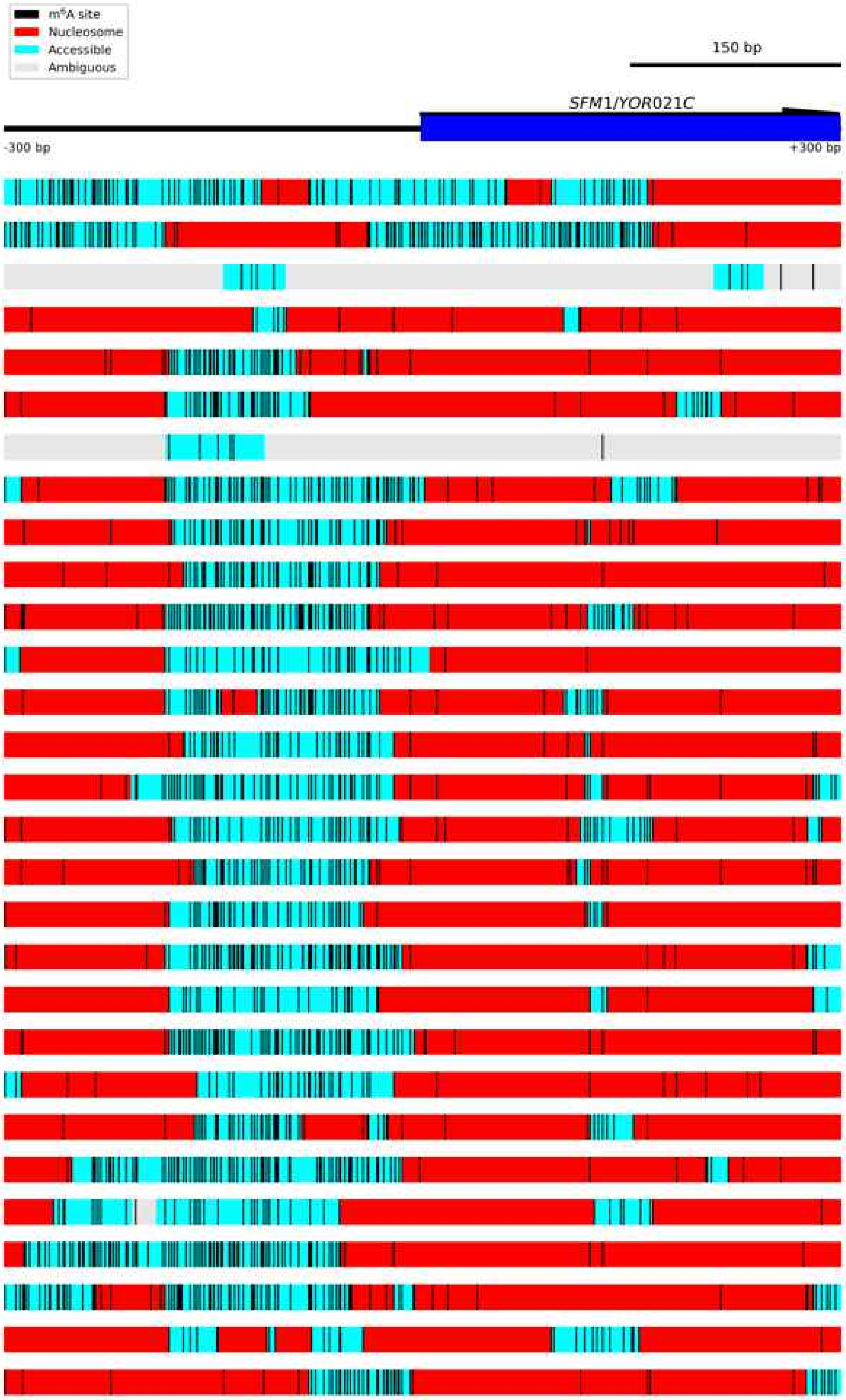

Figure S6.6 **Diagram of *SFM1/YOR021C*.**
The *SFM1/YOR021C* locus was an example of homogeneous nucleosome positioning. Each block below the gene locus represents one read. Accessible regions are indicated by cyan boxes, nucleosomes as red boxes, and ambiguous regions by grey boxes. Called $m^6A$ bases are indicated by vertical black lines.

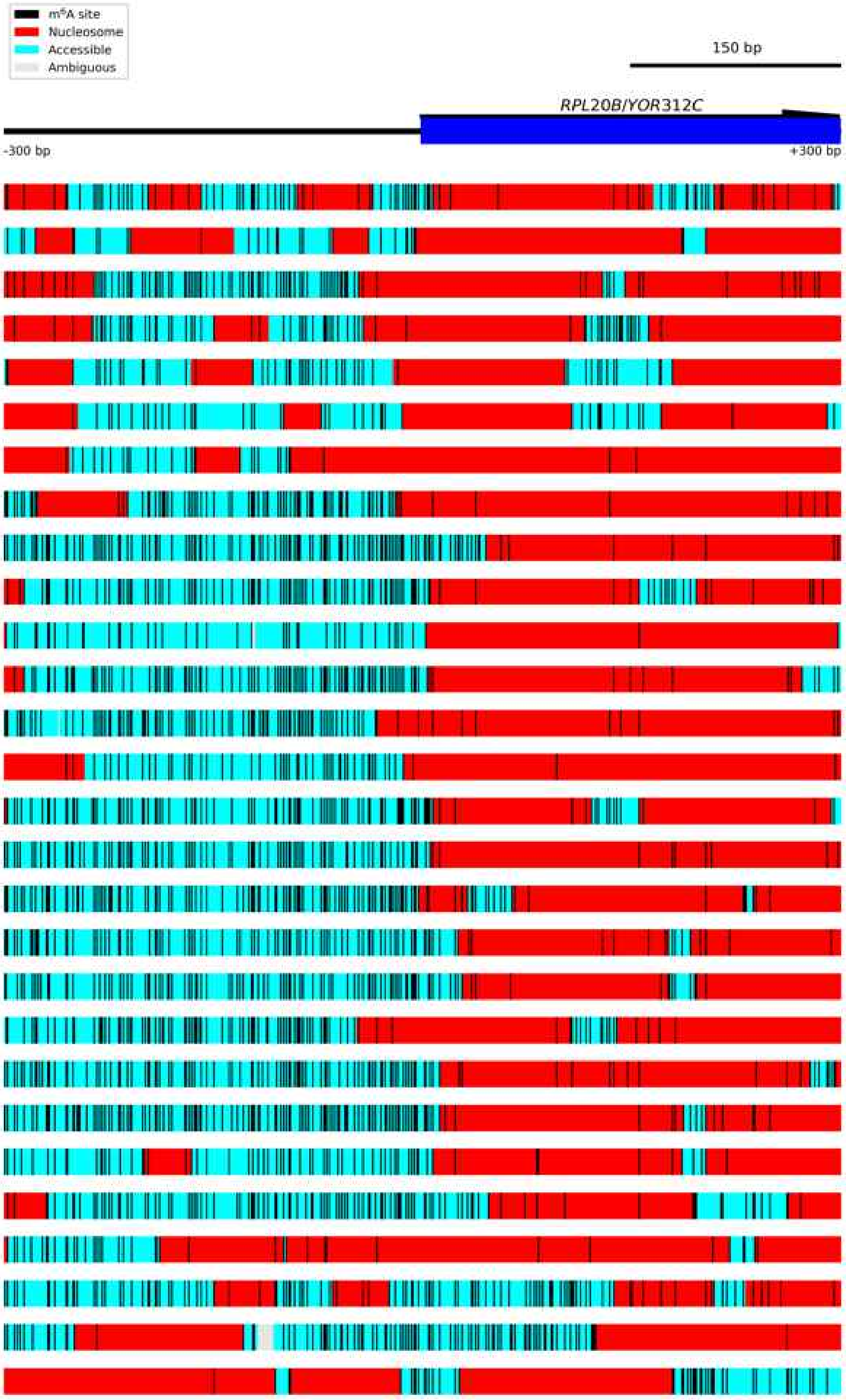

Figure S6.7 **Diagram of *RPL20B/YOR312C*.**
The *RPL20B/YOR312C* locus was an example of homogeneous nucleosome positioning. Each block below the gene locus represents one read. Accessible regions are indicated by cyan boxes, nucleosomes as red boxes, and ambiguous regions by grey boxes. Called m$^6$A bases are indicated by vertical black lines.

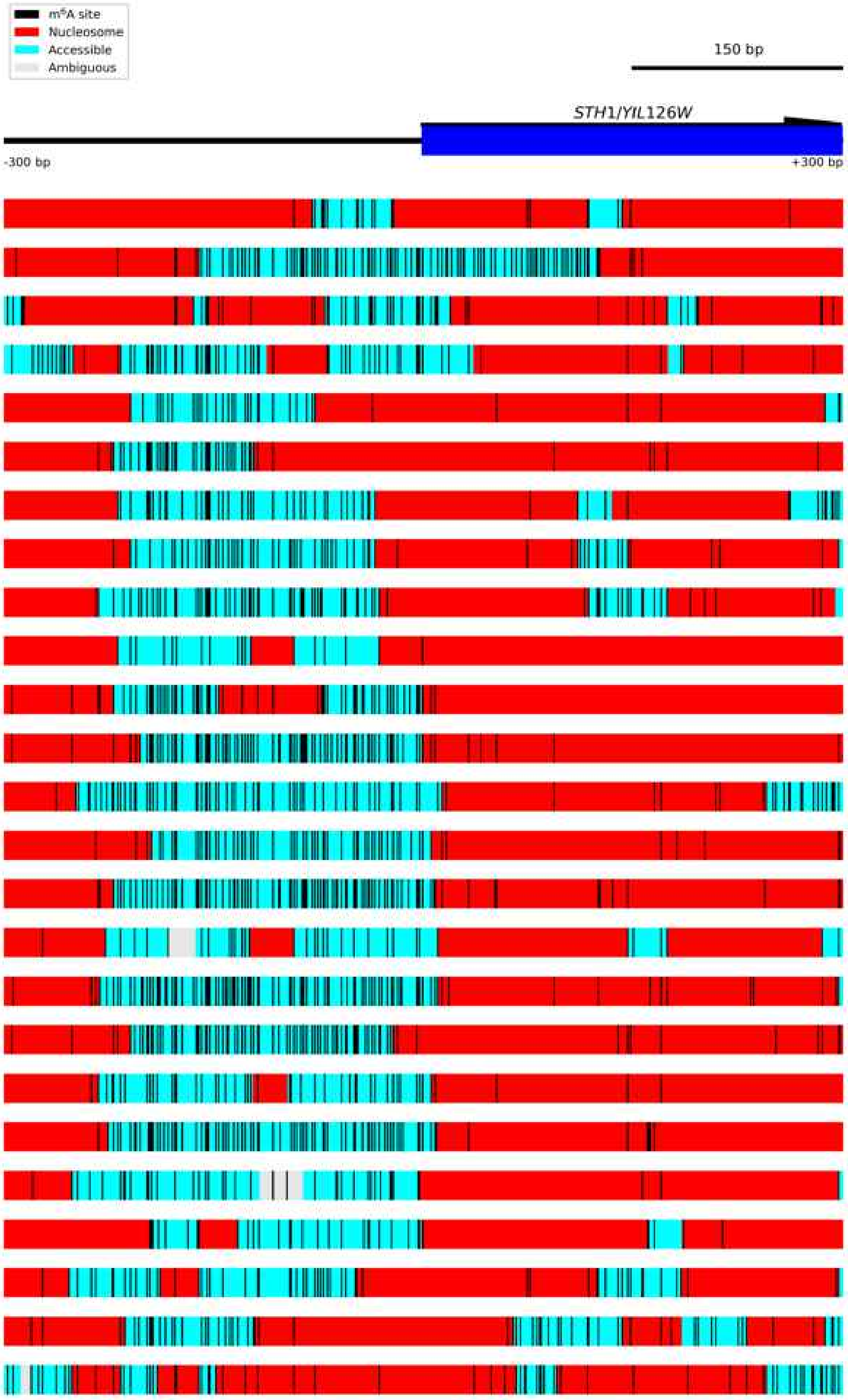

Figure S6.8 **Diagram of *STH1/YIL126W*.**

The *STH1/YIL126W* locus was an example of homogeneous nucleosome positioning. Each block below the gene locus represents one read. Accessible regions are indicated by cyan boxes, nucleosomes as red boxes, and ambiguous regions by grey boxes. Called $m^6A$ bases are indicated by vertical black lines.

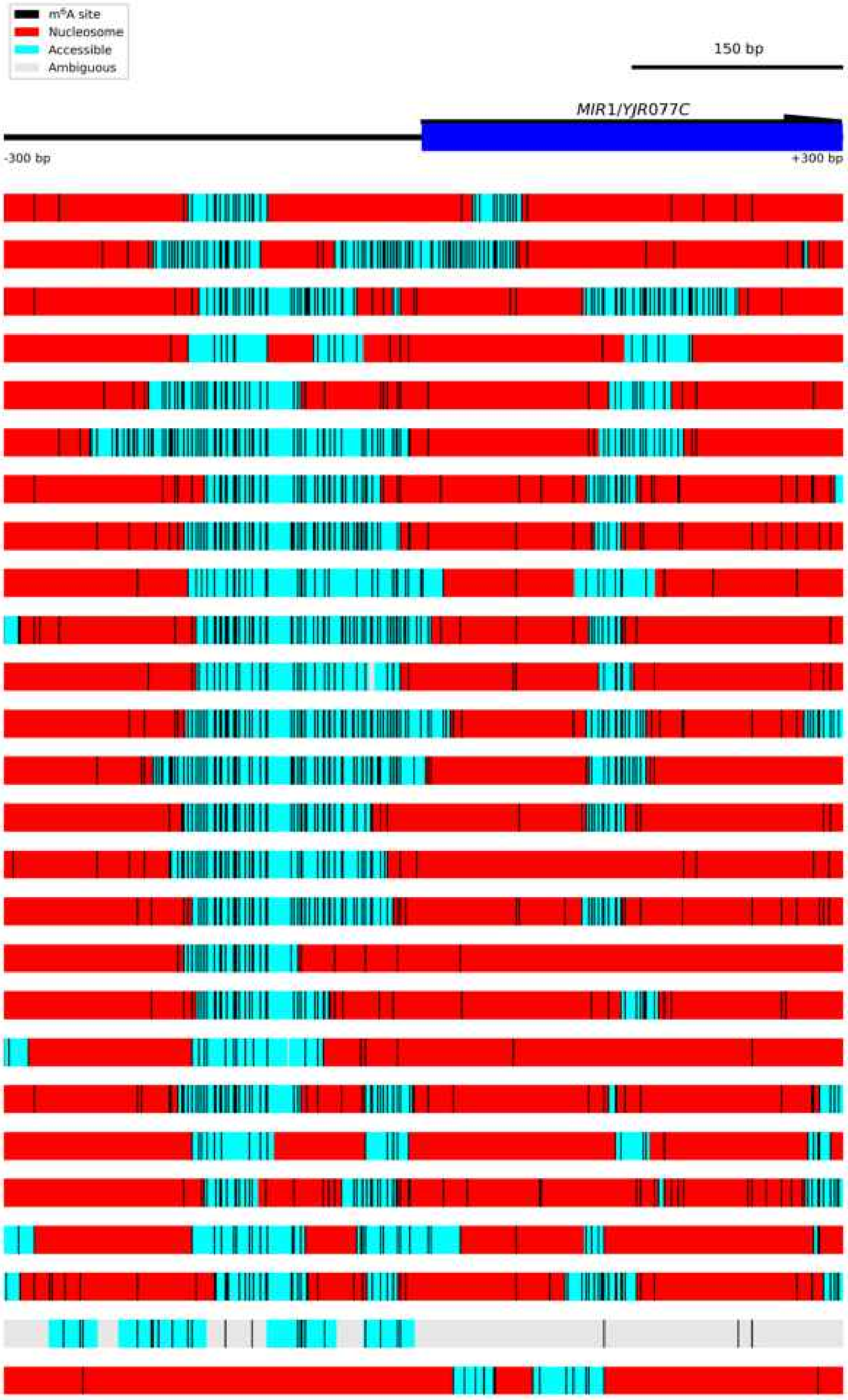

Figure S6.9 **Diagram of *MIR1/YJR077C*.**

The *MIR1/YJR077C* locus was an example of homogeneous nucleosome positioning. Each block below the gene locus represents one read. Accessible regions are indicated by cyan boxes, nucleosomes as red boxes, and ambiguous regions by grey boxes. Called m$^6$A bases are indicated by vertical black lines.

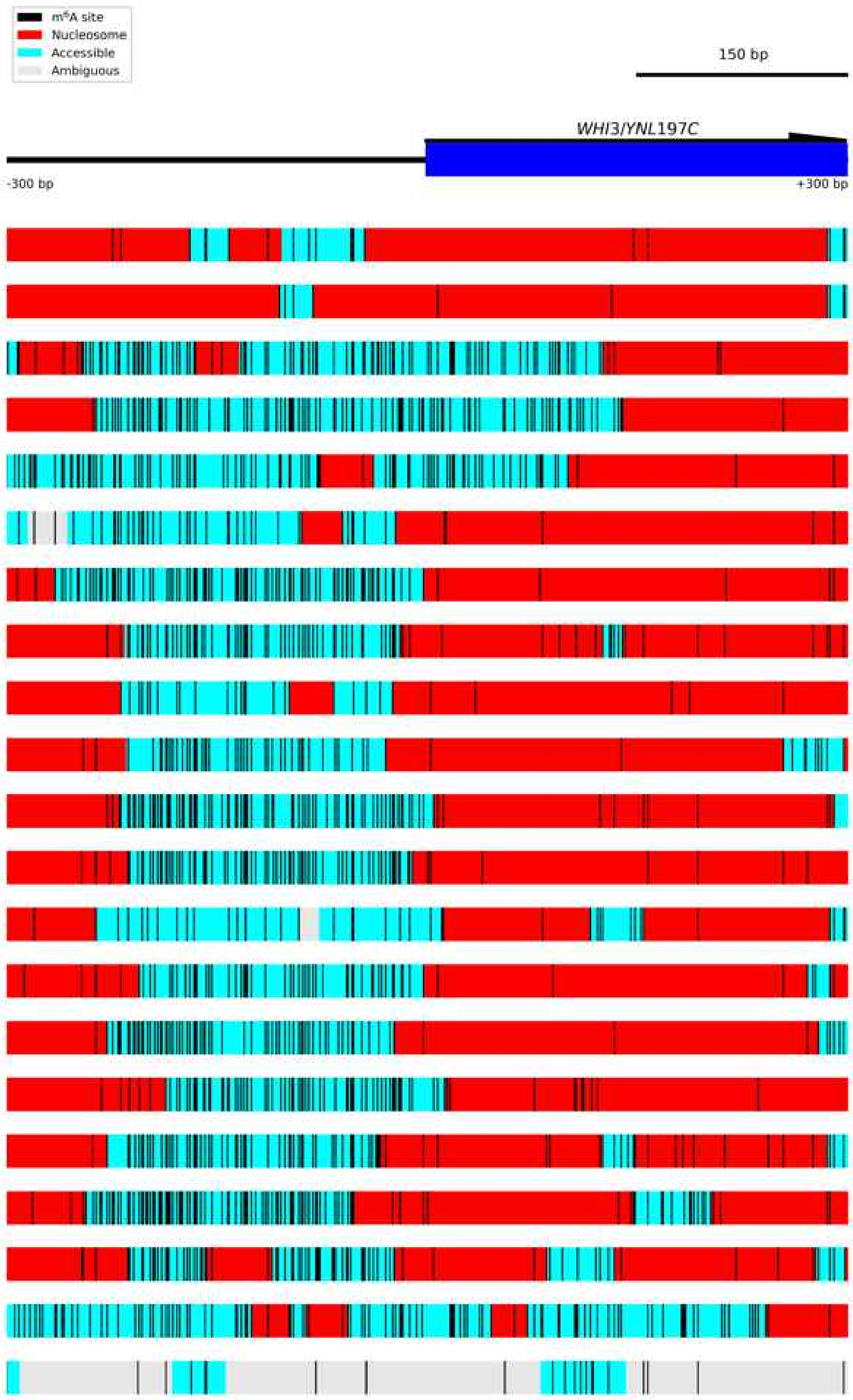

Figure S6.10 **Diagram of *WHI3/YNL197C*.**
The *WHI3/YNL197C* locus was an example of homogeneous nucleosome positioning. Each block below the gene locus represents one read. Accessible regions are indicated by cyan boxes, nucleosomes as red boxes, and ambiguous regions by grey boxes. Called m$^6$A bases are indicated by vertical black lines.

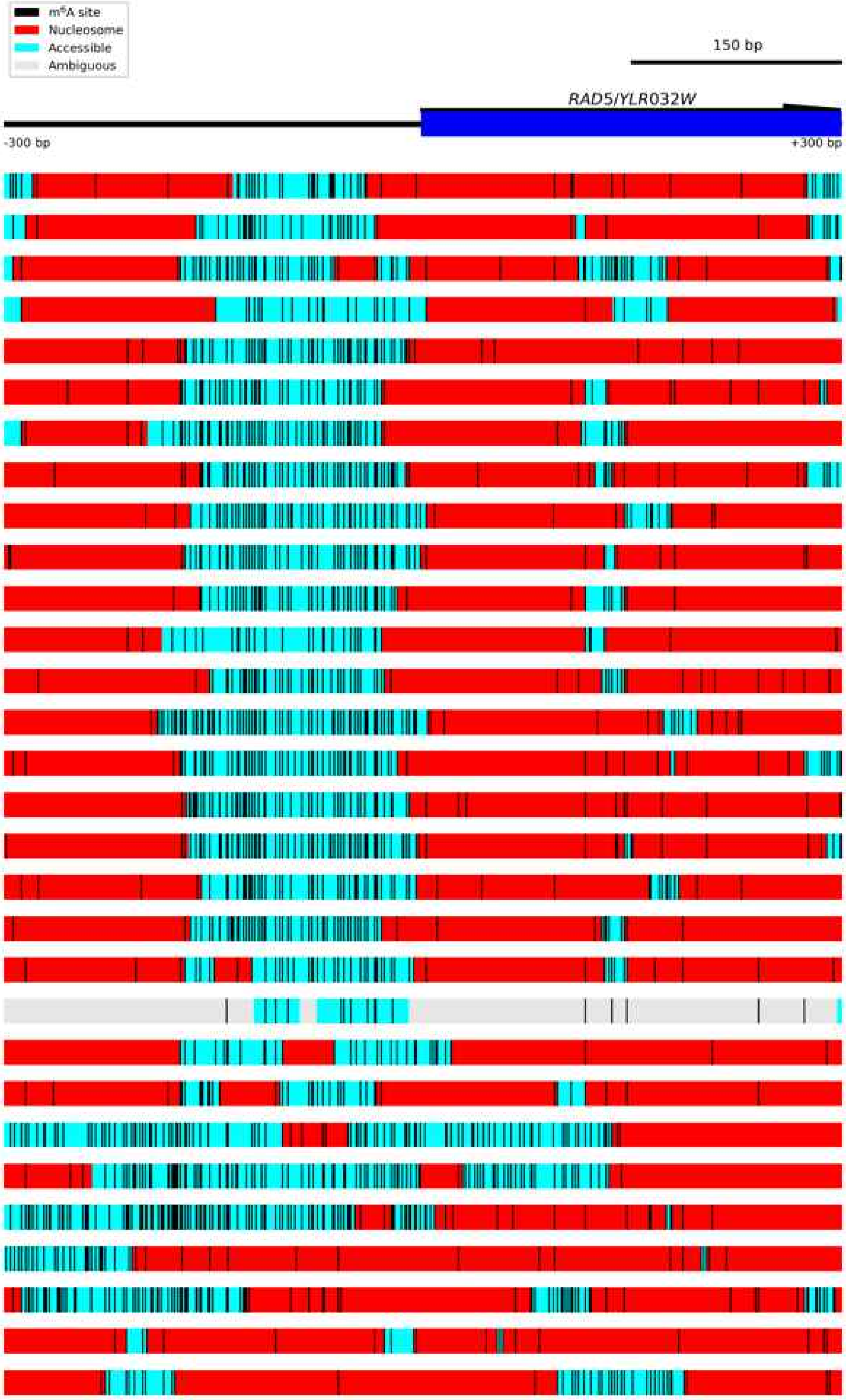

Figure S6.11 **Diagram of *RAD5/YLR032W*.**
The *RAD5/YLR032W* locus was an example of homogeneous nucleosome positioning. Each block below the gene locus represents one read. Accessible regions are indicated by cyan boxes, nucleosomes as red boxes, and ambiguous regions by grey boxes. Called m$^6$A bases are indicated by vertical black lines.

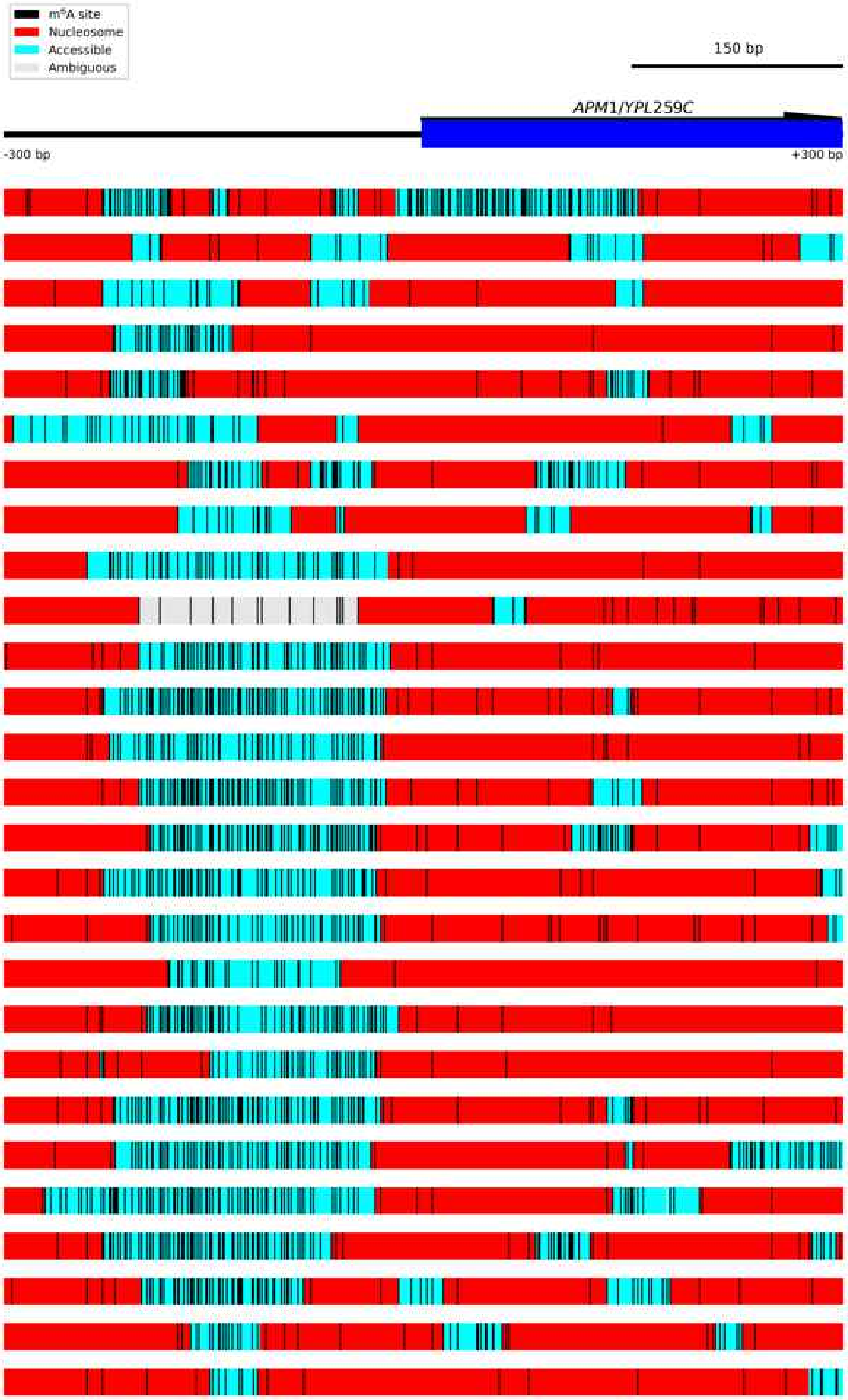

Figure S6.12 **Diagram of *APM1/YPL259C*.**
The *APM1/YPL259C* locus was an example of homogeneous nucleosome positioning. Each block below the gene locus represents one read. Accessible regions are indicated by cyan boxes, nucleosomes as red boxes, and ambiguous regions by grey boxes. Called $m^6A$ bases are indicated by vertical black lines.

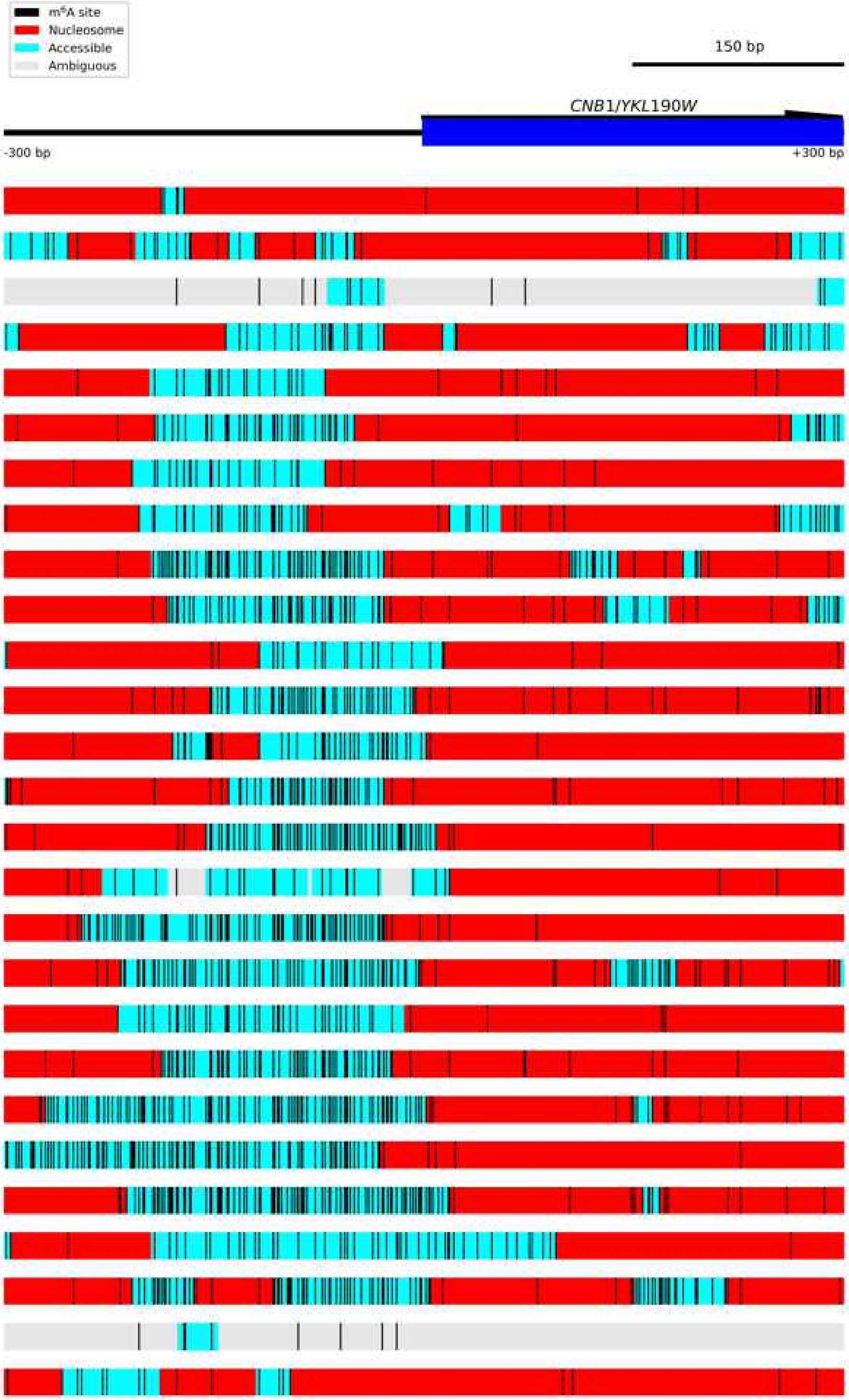

Figure S6.13 **Diagram of *CNB1/YKL190W*.**
The *CNB1/YKL190W* locus was an example of homogeneous nucleosome positioning. Each block below the gene locus represents one read. Accessible regions are indicated by cyan boxes, nucleosomes as red boxes, and ambiguous regions by grey boxes. Called $m^6A$ bases are indicated by vertical black lines.

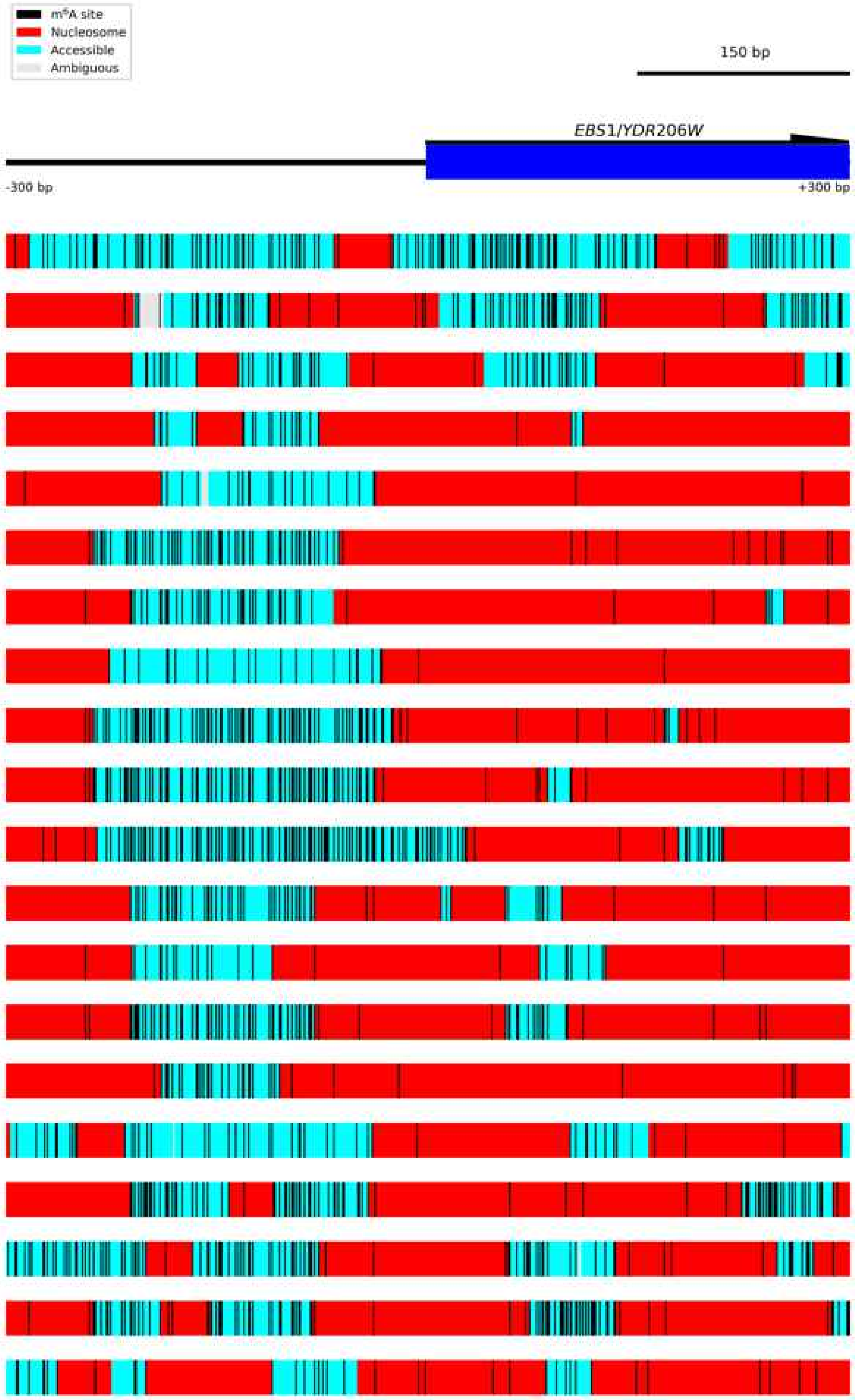

Figure S6.14 **Diagram of *EBS1/YDR206W*.**
The *EBS1/YDR206W* locus was an example of homogeneous nucleosome positioning. Each block below the gene locus represents one read. Accessible regions are indicated by cyan boxes, nucleosomes as red boxes, and ambiguous regions by grey boxes. Called $m^6A$ bases are indicated by vertical black lines.

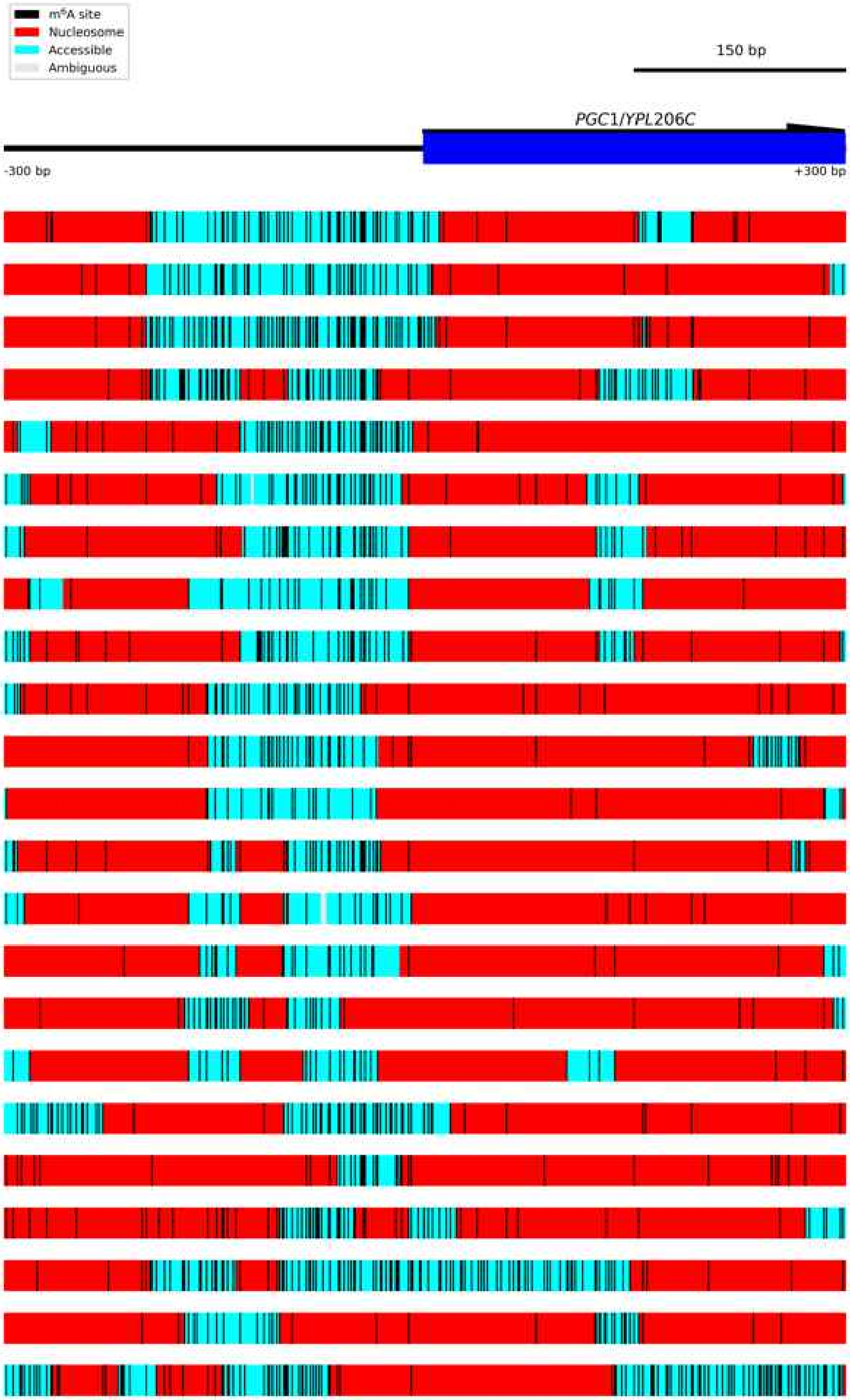

Figure S6.15 **Diagram of *PGC1/YPL206C*.**
The *PGC1/YPL206C* locus was an example of homogeneous nucleosome positioning. Each block below the gene locus represents one read. Accessible regions are indicated by cyan boxes, nucleosomes as red boxes, and ambiguous regions by grey boxes. Called $m^6A$ bases are indicated by vertical black lines.

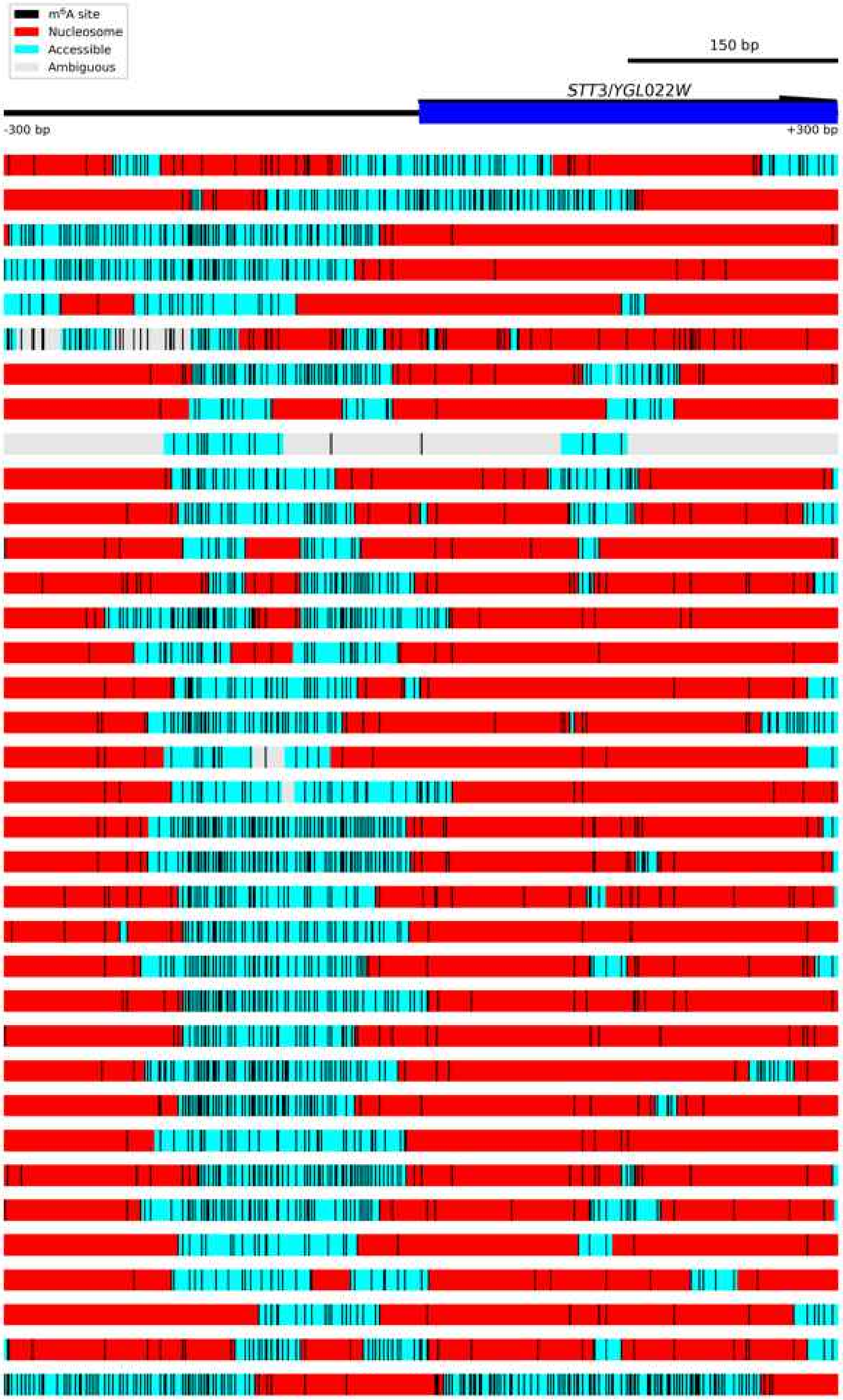

Figure S6.16 **Diagram of *STT3/YGL022W*.**
The *STT3/YGL022W* locus was an example of homogeneous nucleosome positioning. Each block below the gene locus represents one read. Accessible regions are indicated by cyan boxes, nucleosomes as red boxes, and ambiguous regions by grey boxes. Called $m^6A$ bases are indicated by vertical black lines.

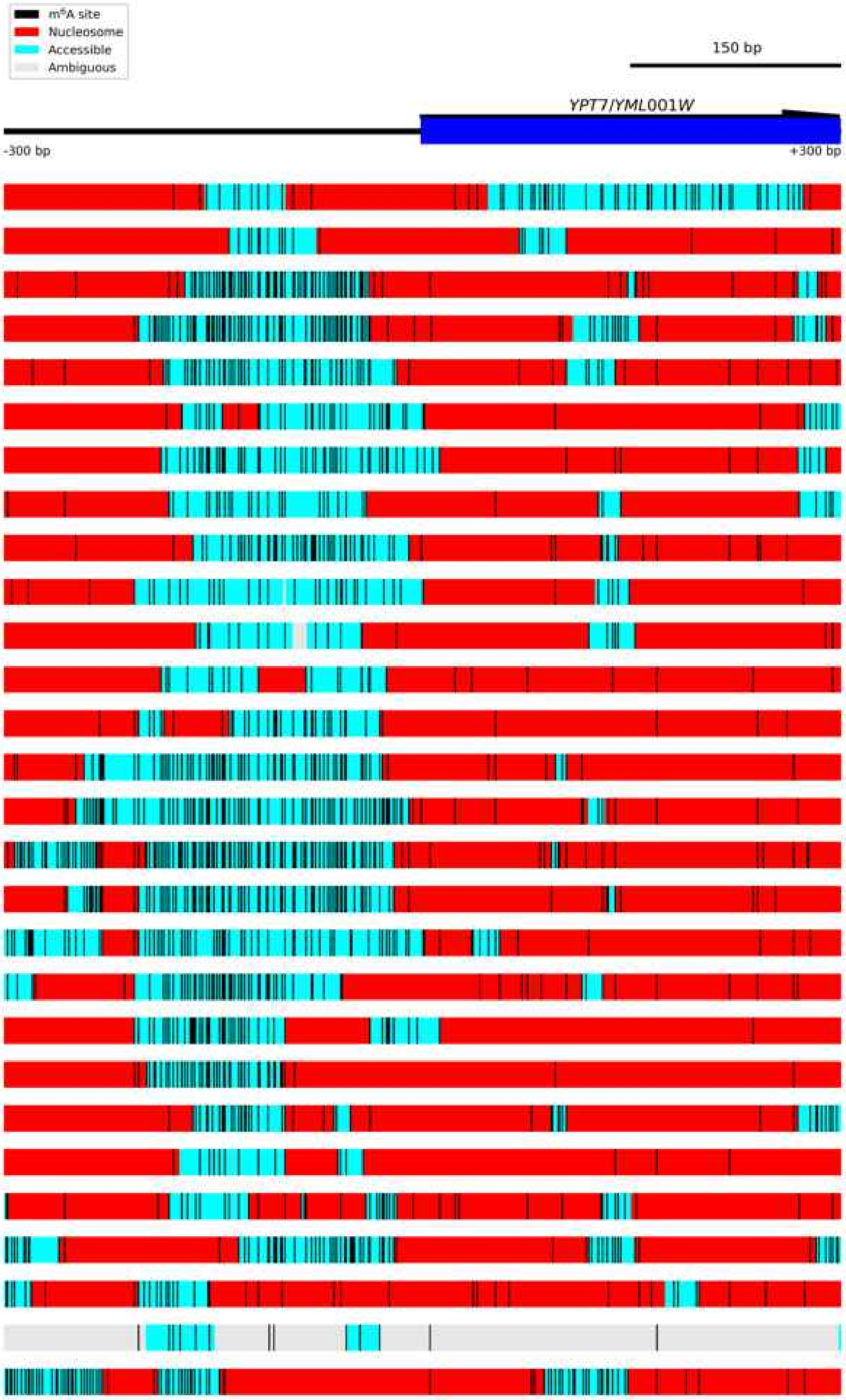

Figure S6.17 **Diagram of *YPT7/YML001W*.**
The *YPT7/YML001W* locus was an example of homogeneous nucleosome positioning. Each block below the gene locus represents one read. Accessible regions are indicated by cyan boxes, nucleosomes as red boxes, and ambiguous regions by grey boxes. Called $m^6A$ bases are indicated by vertical black lines.

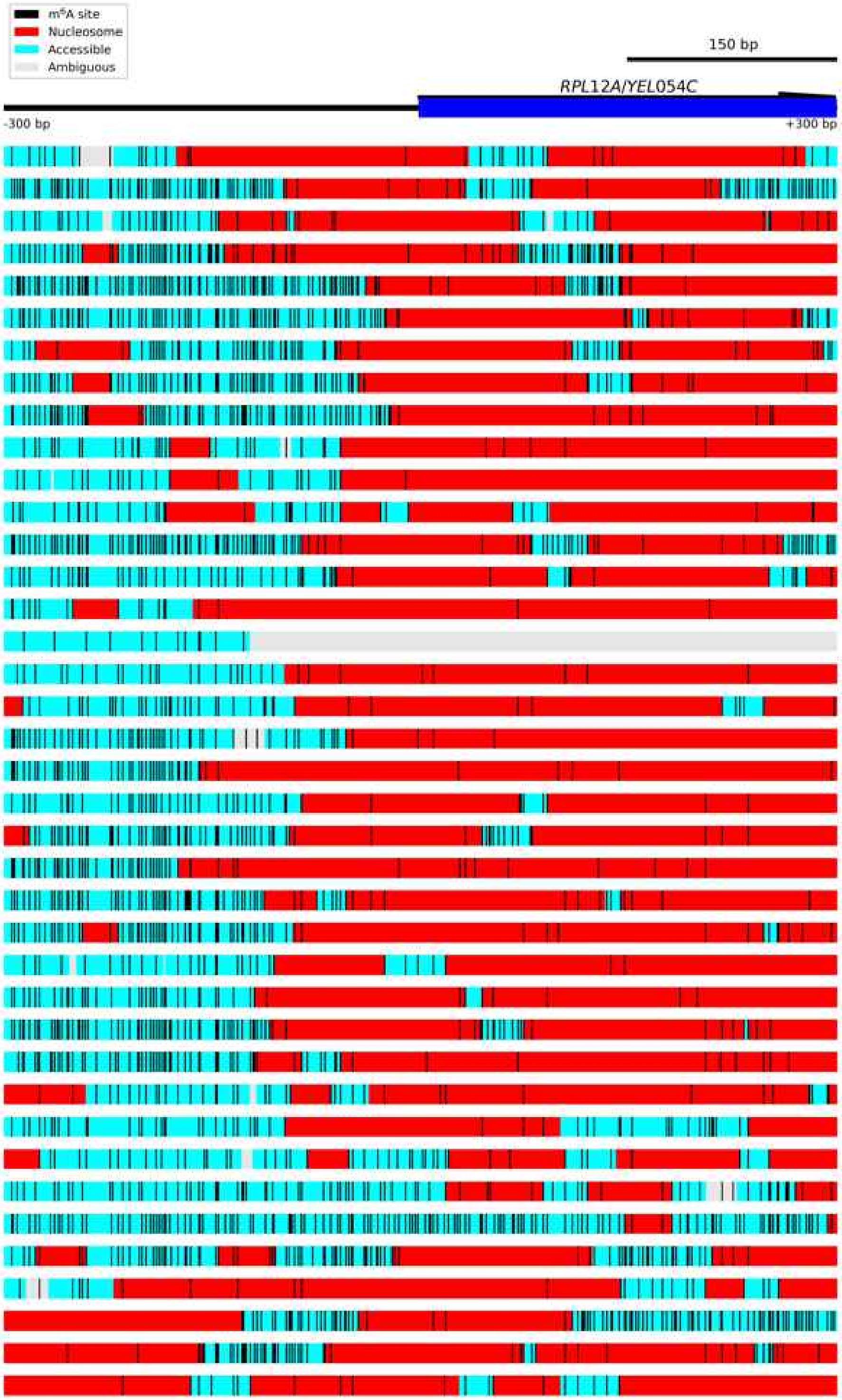

Figure S6.18 **Diagram of *RPL12A/YEL054C*.**
The *RPL12A/YEL054C* locus was an example of homogeneous nucleosome positioning. Each block below the gene locus represents one read. Accessible regions are indicated by cyan boxes, nucleosomes as red boxes, and ambiguous regions by grey boxes. Called m$^6$A bases are indicated by vertical black lines.

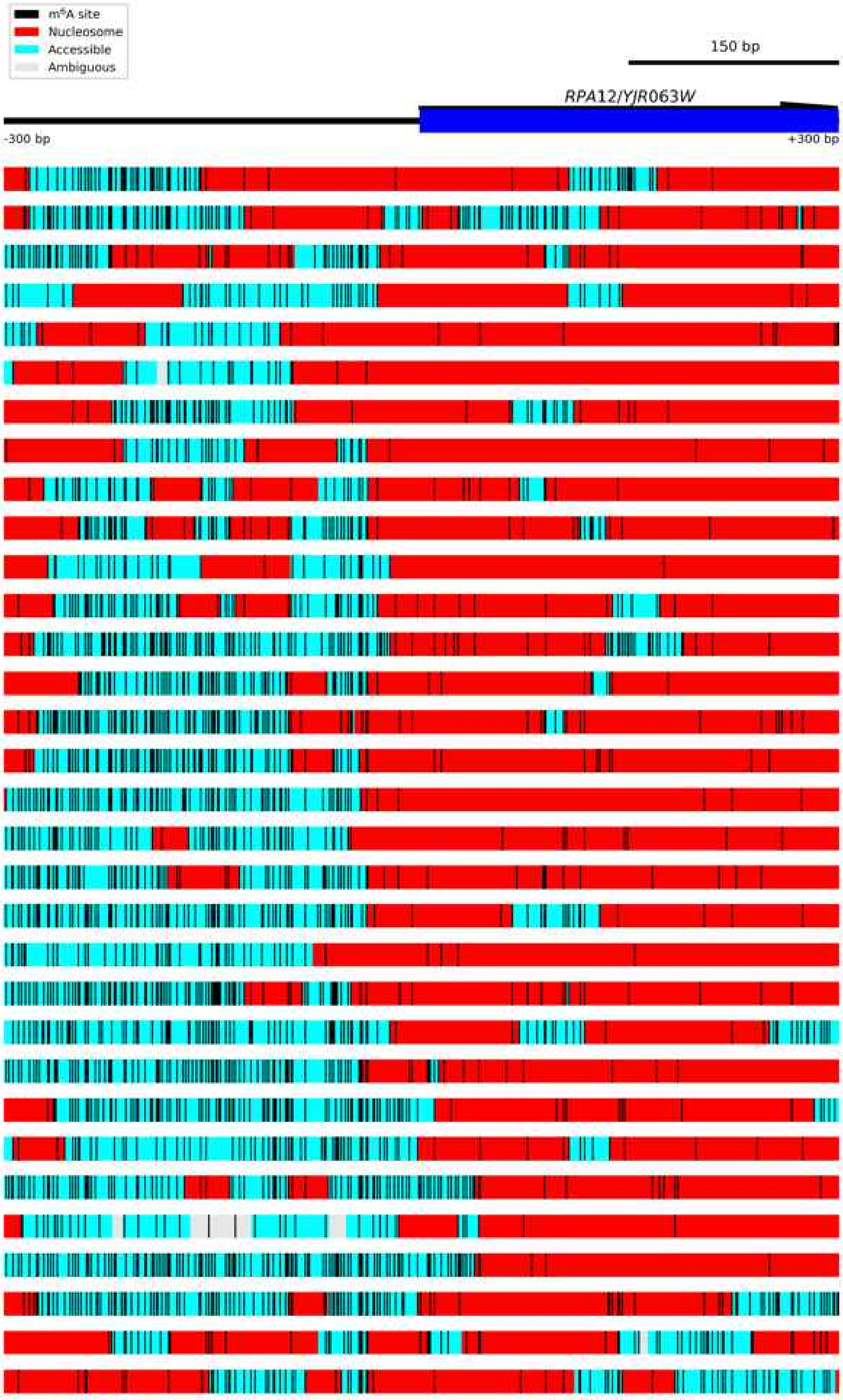

Figure S6.19 **Diagram of *RPA12/YJR063W*.**
The *RPA12/YJR063W* locus was an example of homogeneous nucleosome positioning. Each block below the gene locus represents one read. Accessible regions are indicated by cyan boxes, nucleosomes as red boxes, and ambiguous regions by grey boxes. Called $m^6A$ bases are indicated by vertical black lines.

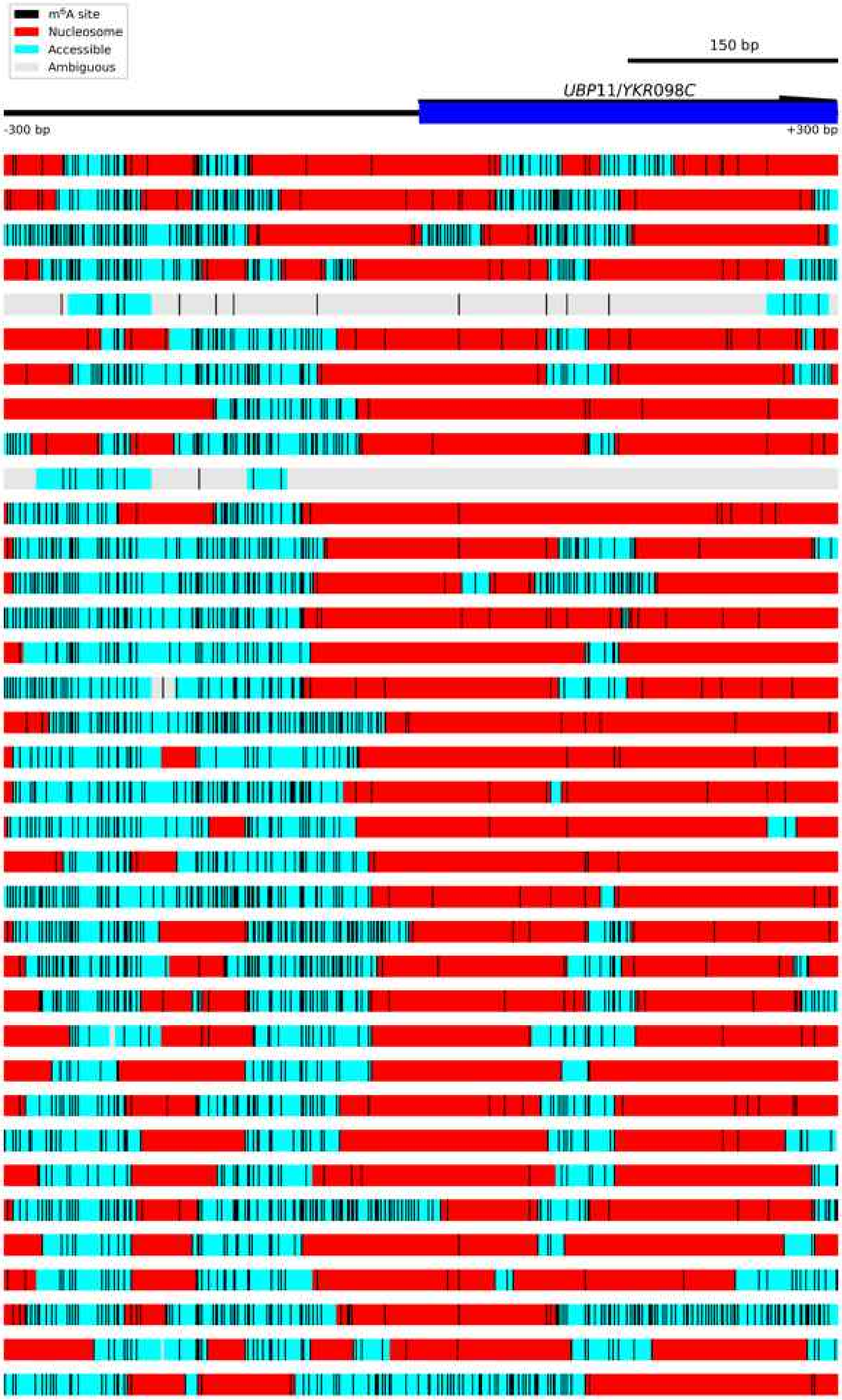

Figure S6.20 **Diagram of *UBP11/YKR098C*.**
The *UBP11/YKR098C* locus was an example of homogeneous nucleosome positioning. Each block below the gene locus represents one read. Accessible regions are indicated by cyan boxes, nucleosomes as red boxes, and ambiguous regions by grey boxes. Called m$^6$A bases are indicated by vertical black lines.

#### Supplementary Figure 7. Nucleosome positioning on the 20 most heterogeneous genes.

Nucleosome positioning on the 20 most heterogeneous genes. The genes are ordered by heterogeneity. Each block below the gene locus represents one read. Accessible regions are indicated by cyan boxes, nucleosomes as red boxes, and ambiguous regions by grey boxes. Called $m^6$A bases are indicated by vertical black lines.

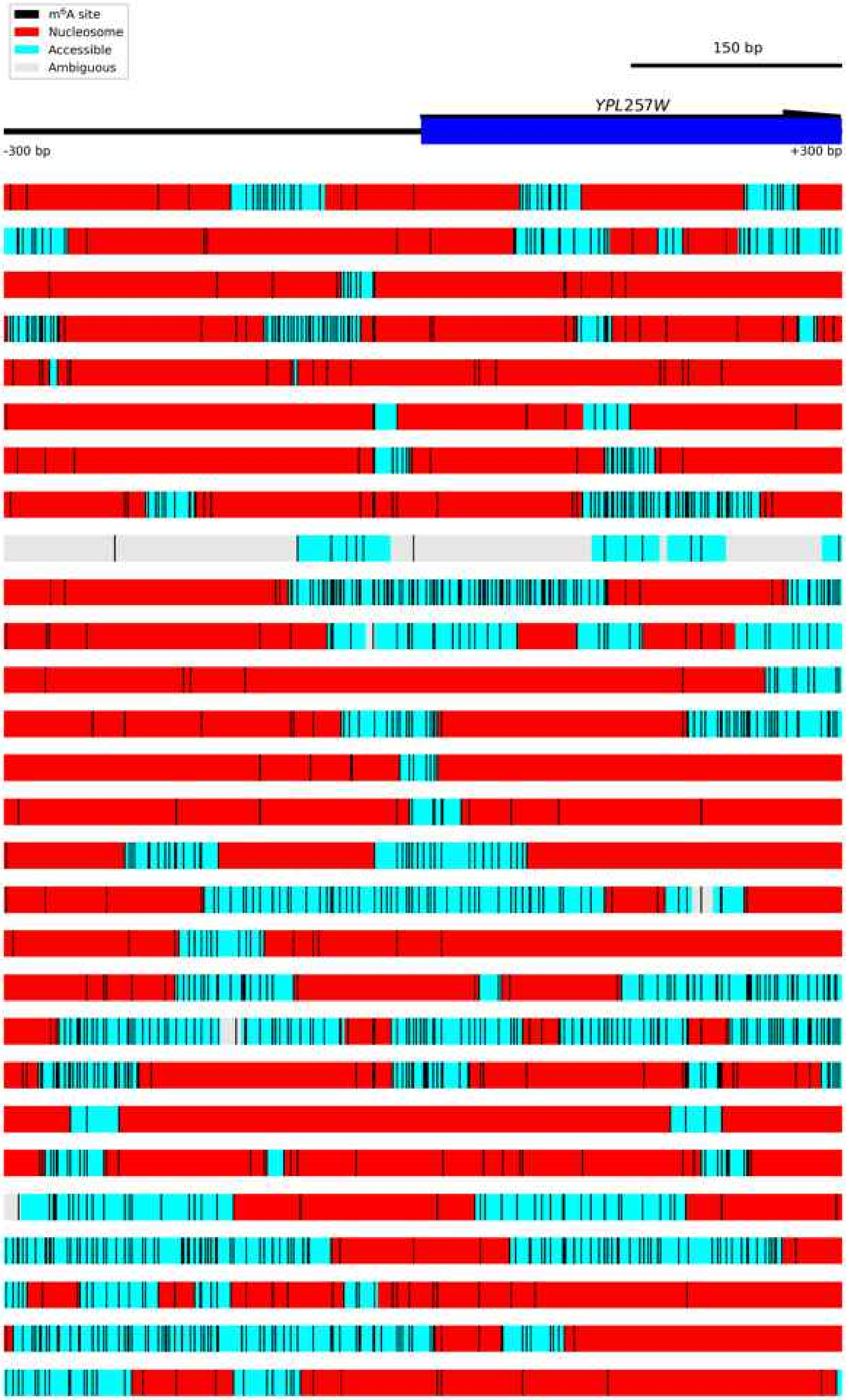

Figure S7.1 **Diagram of *YPL257W*.**

The *YPL257W* locus was an example of heterogeneous nucleosome positioning. Each block below the gene locus represents one read. Accessible regions are indicated by cyan boxes, nucleosomes as red boxes, and ambiguous regions by grey boxes. Called m$^6$A bases are indicated by vertical black lines.

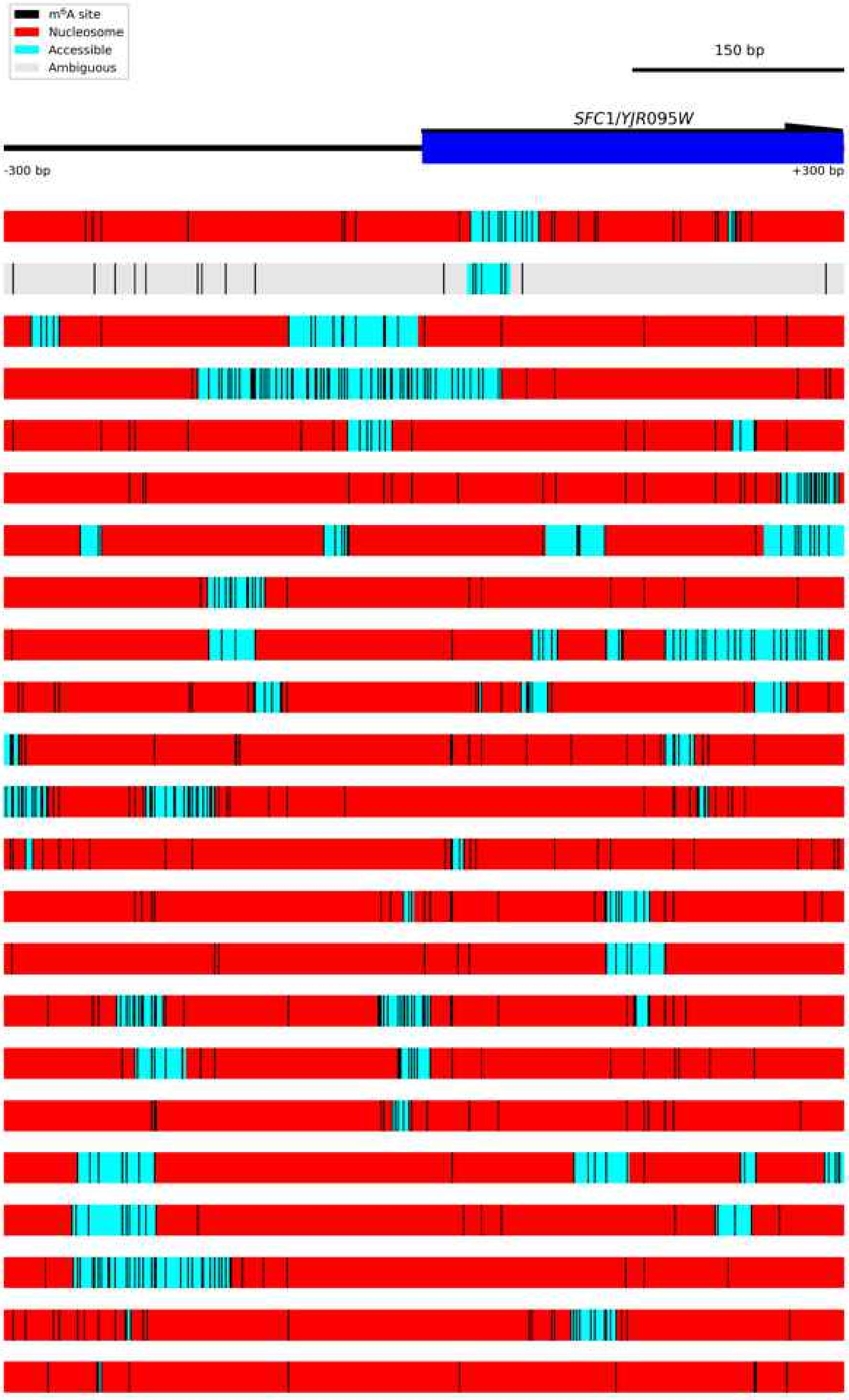

Figure S7.2 **Diagram of *SFC1/YJR095W*.**

The *SFC1/YJR095W* locus was an example of heterogeneous nucleosome positioning. Each block below the gene locus represents one read. Accessible regions are indicated by cyan boxes, nucleosomes as red boxes, and ambiguous regions by grey boxes. Called $m^6A$ bases are indicated by vertical black lines.

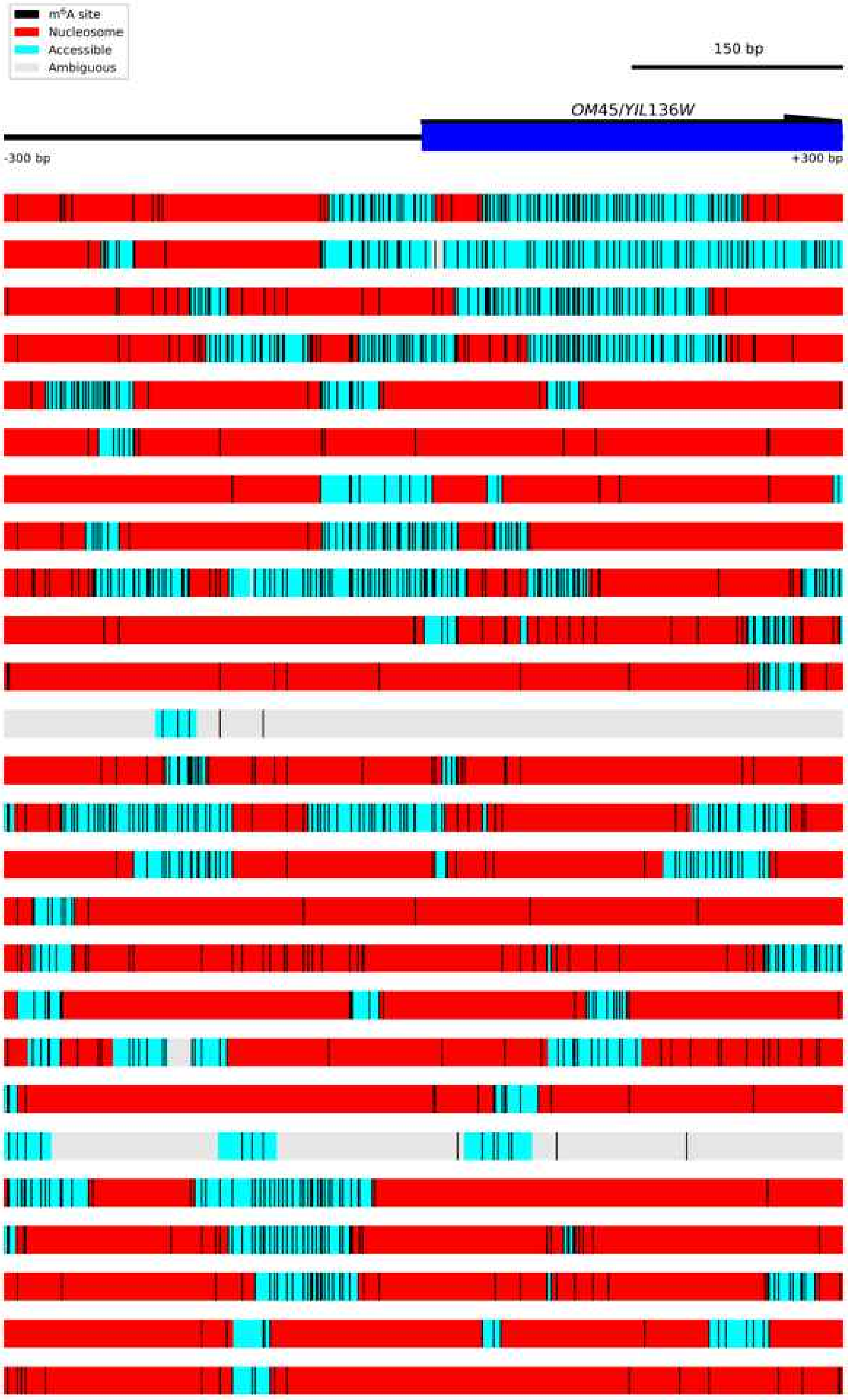

Figure S7.3 **Diagram of *OM45/YIL136W*.**
The *OM45/YIL136W* locus was an example of heterogeneous nucleosome positioning. Each block below the gene locus represents one read. Accessible regions are indicated by cyan boxes, nucleosomes as red boxes, and ambiguous regions by grey boxes. Called $m^6A$ bases are indicated by vertical black lines.

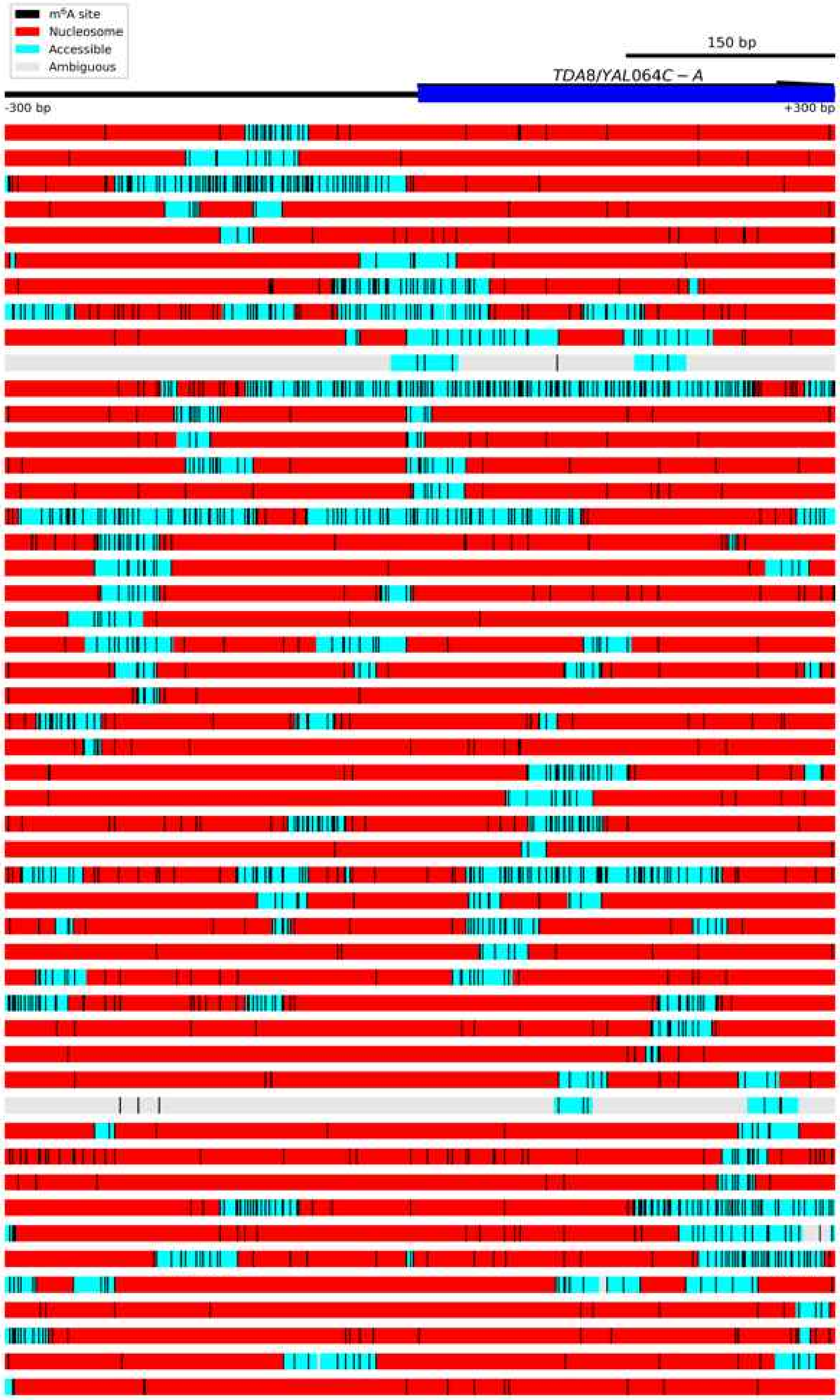

Figure S7.4 **Diagram of *TDA8/YAL064C-A*.**
The *TDA8/YAL064C-A* locus was an example of heterogeneous nucleosome positioning. Each block below the gene locus represents one read. Accessible regions are indicated by cyan boxes, nucleosomes as red boxes, and ambiguous regions by grey boxes. Called m$^6$A bases are indicated by vertical black lines.

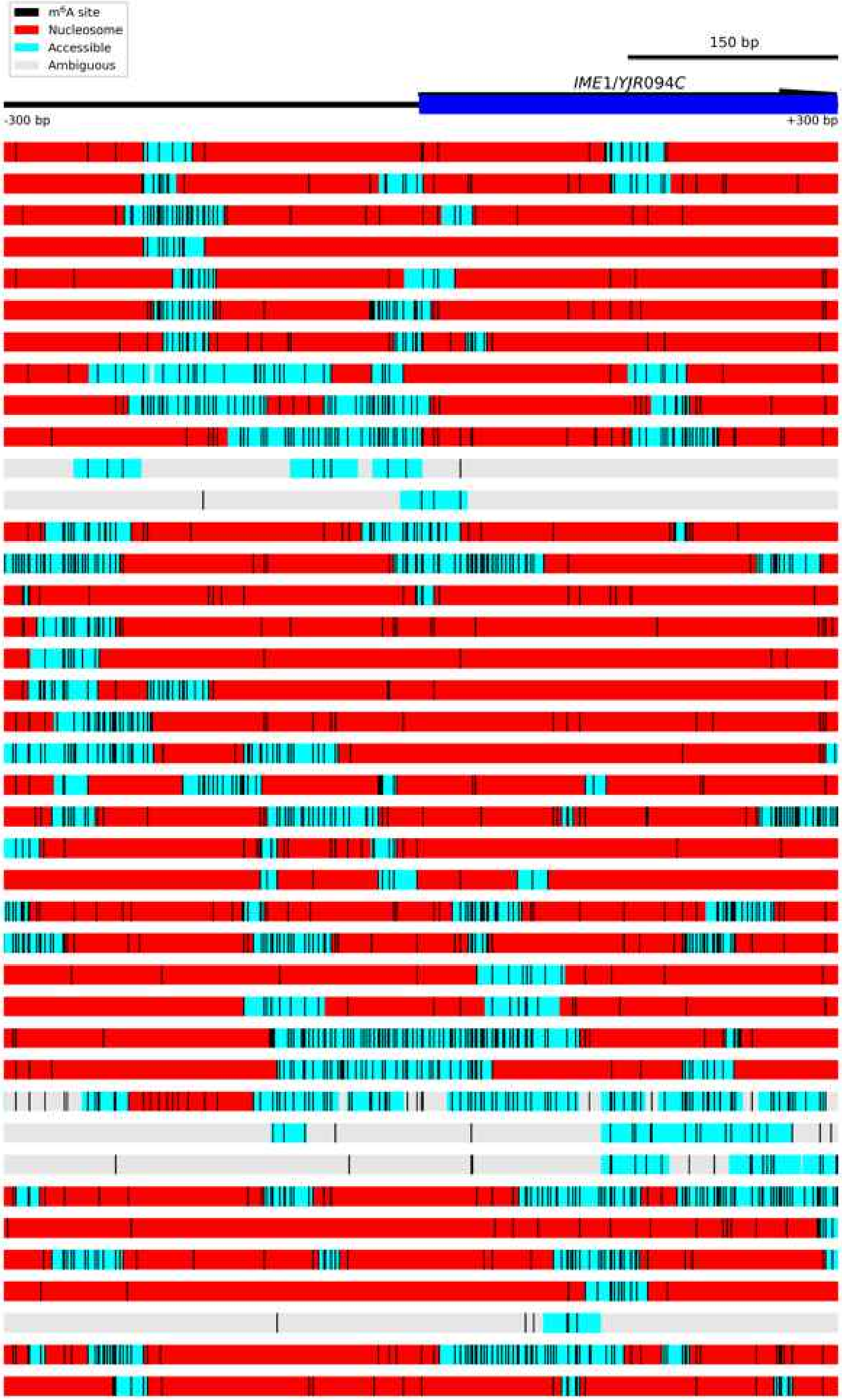

Figure S7.5 **Diagram of *IME1/YJR094C*.**

The *IME1/YJR094C* locus was an example of heterogeneous nucleosome positioning. Each block below the gene locus represents one read. Accessible regions are indicated by cyan boxes, nucleosomes as red boxes, and ambiguous regions by grey boxes. Called $m^6A$ bases are indicated by vertical black lines.

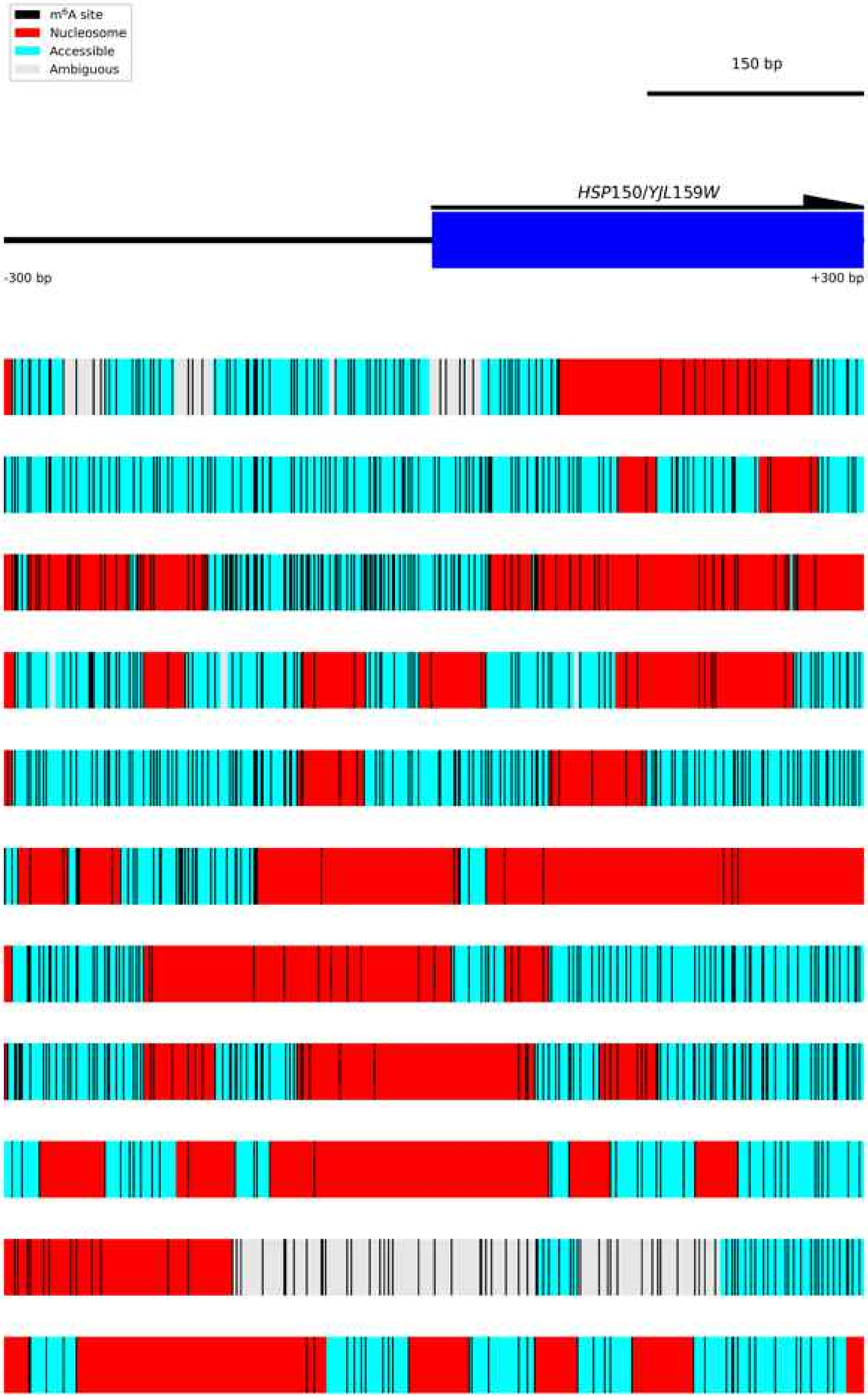

Figure S7.6 **Diagram of *HSP150/YJL159W*.**
The *HSP150/YJL159W* locus was an example of heterogeneous nucleosome positioning. Each block below the gene locus represents one read. Accessible regions are indicated by cyan boxes, nucleosomes as red boxes, and ambiguous regions by grey boxes. Called $m^6A$ bases are indicated by vertical black lines.

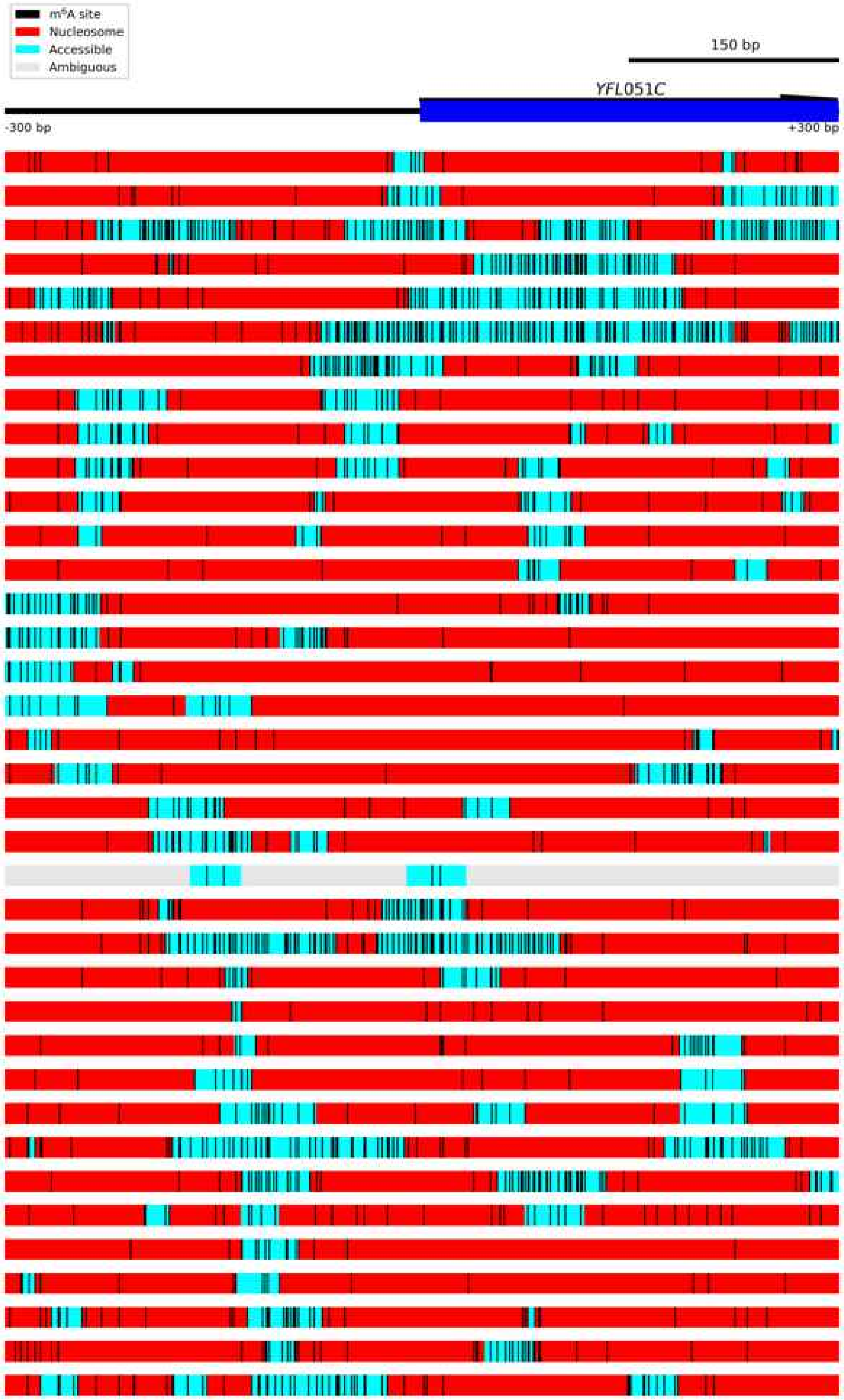

Figure S7.7 **Diagram of *YFL051C*.**

The *YFL051C* locus was an example of heterogeneous nucleosome positioning. Each block below the gene locus represents one read. Accessible regions are indicated by cyan boxes, nucleosomes as red boxes, and ambiguous regions by grey boxes. Called m$^6$A bases are indicated by vertical black lines.

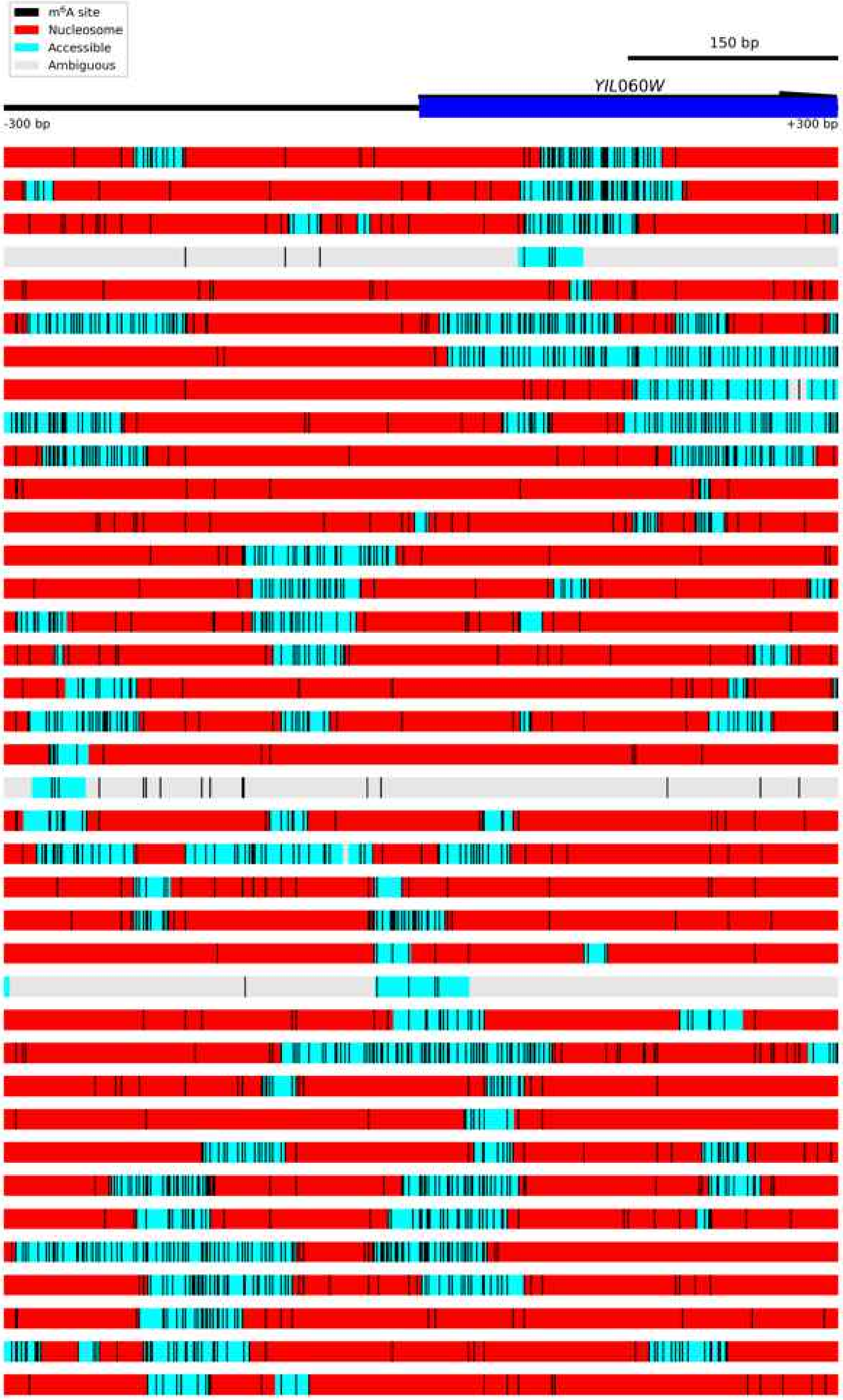

Figure S7.8 **Diagram of *YIL060W*.**

The *YIL060W* locus was an example of heterogeneous nucleosome positioning. Each block below the gene locus represents one read. Accessible regions are indicated by cyan boxes, nucleosomes as red boxes, and ambiguous regions by grey boxes. Called $m^6A$ bases are indicated by vertical black lines.

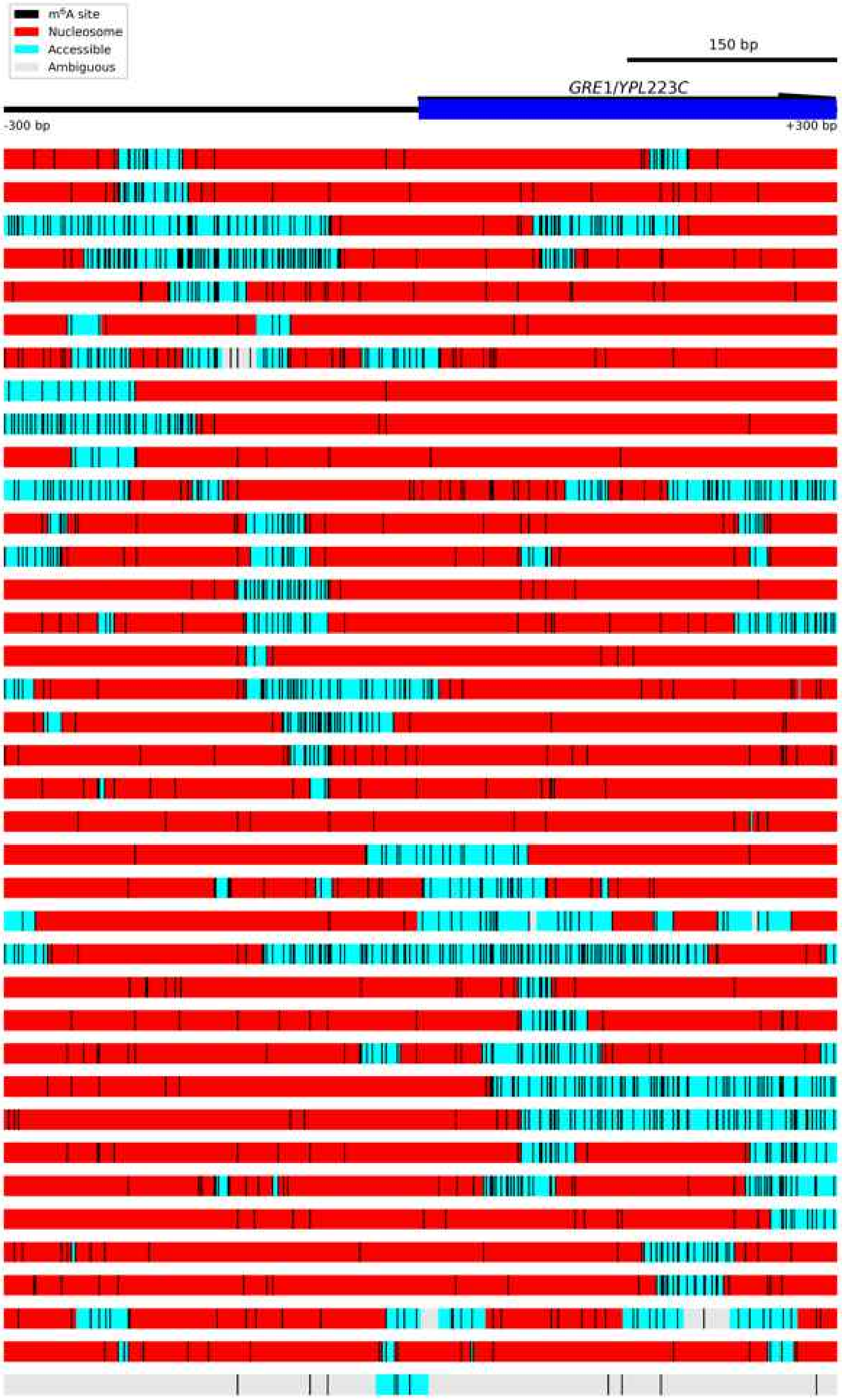

Figure S7.9 **Diagram of *GRE1/YPL223C*.**
The *GRE1/YPL223C* locus was an example of heterogeneous nucleosome positioning. Each block below the gene locus represents one read. Accessible regions are indicated by cyan boxes, nucleosomes as red boxes, and ambiguous regions by grey boxes. Called m$^6$A bases are indicated by vertical black lines.

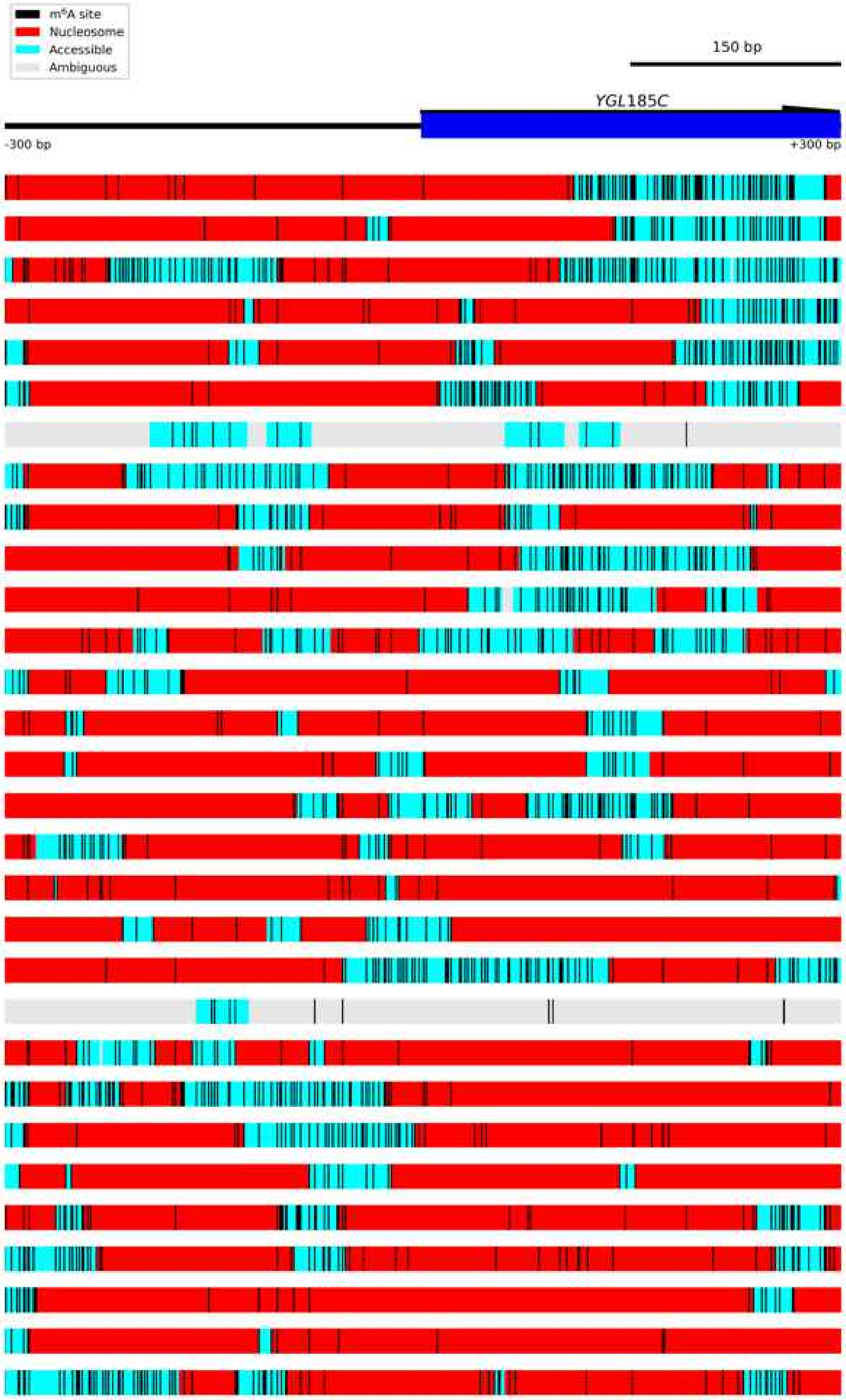

Figure S7.10 **Diagram of *YGL185C*.**
The *YGL185C* locus was an example of heterogeneous nucleosome positioning. Each block below the gene locus represents one read. Accessible regions are indicated by cyan boxes, nucleosomes as red boxes, and ambiguous regions by grey boxes. Called $m^6A$ bases are indicated by vertical black lines.

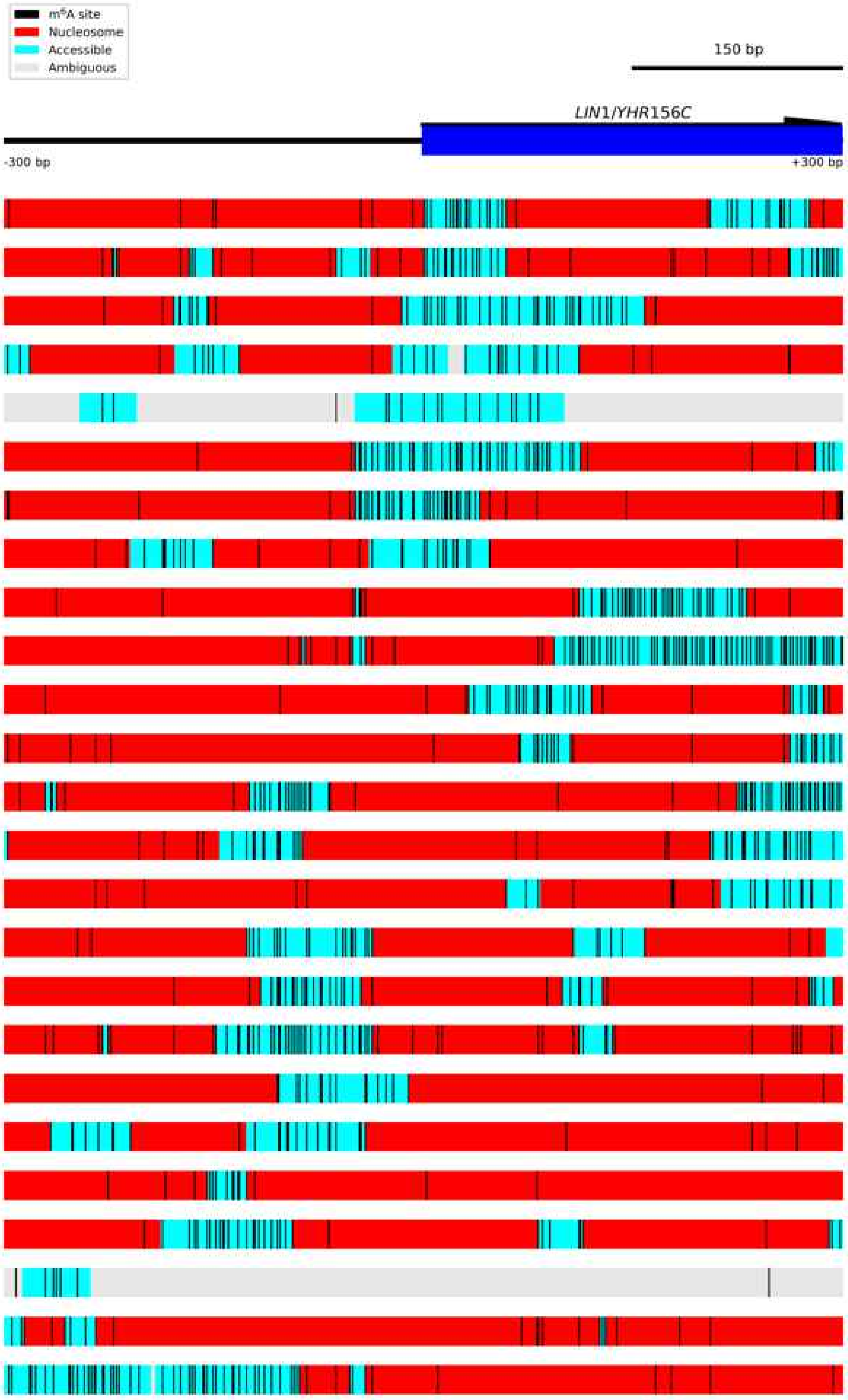

Figure S7.11 **Diagram of *LIN1/YHR156C*.**
The *LIN1/YHR156C* locus was an example of heterogeneous nucleosome positioning. Each block below the gene locus represents one read. Accessible regions are indicated by cyan boxes, nucleosomes as red boxes, and ambiguous regions by grey boxes. Called $m^6A$ bases are indicated by vertical black lines.

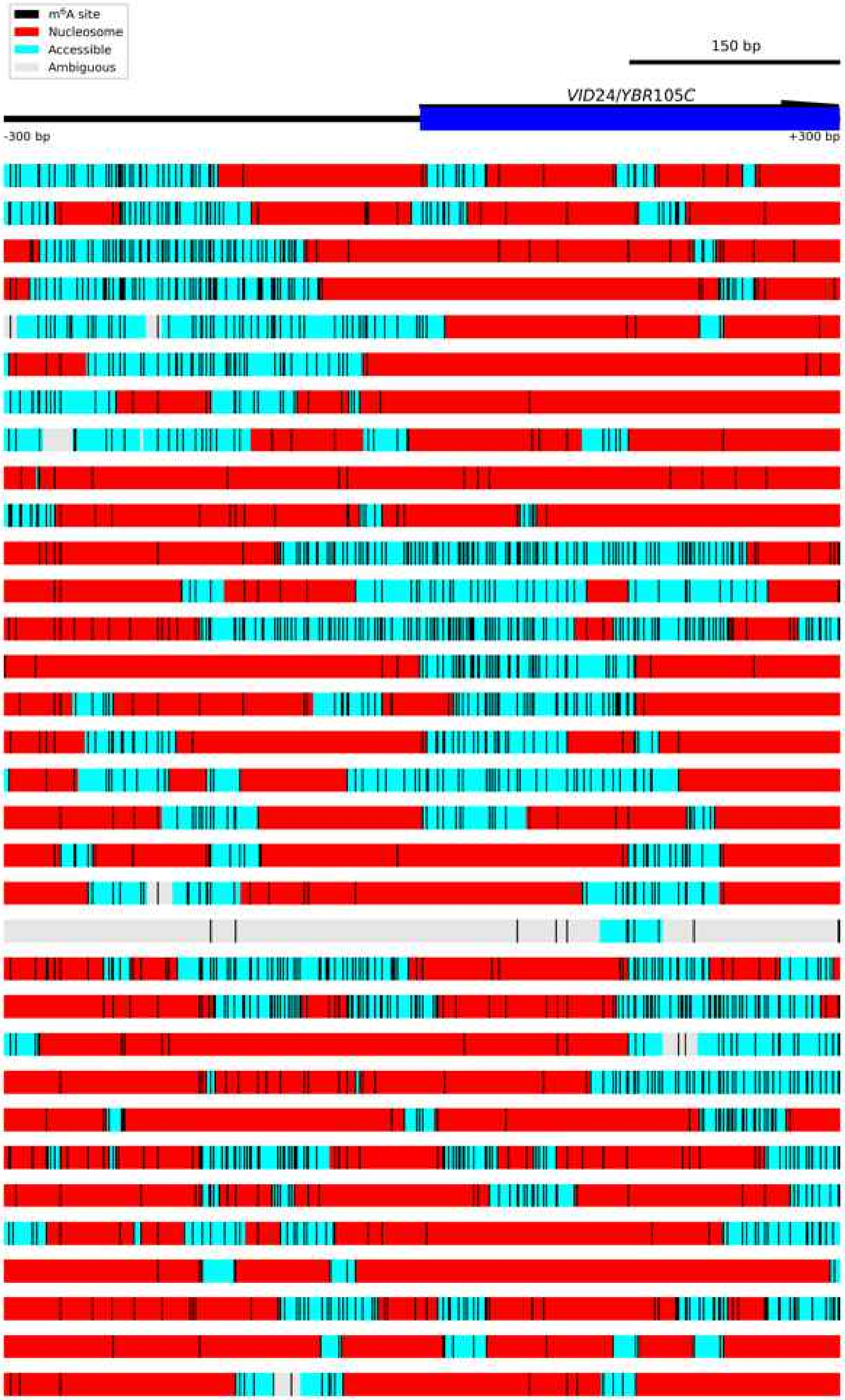

Figure S7.12 **Diagram of *VID24/YBR105C*.**
The *VID24/YBR105C* locus was an example of heterogeneous nucleosome positioning. Each block below the gene locus represents one read. Accessible regions are indicated by cyan boxes, nucleosomes as red boxes, and ambiguous regions by grey boxes. Called $m^6A$ bases are indicated by vertical black lines.

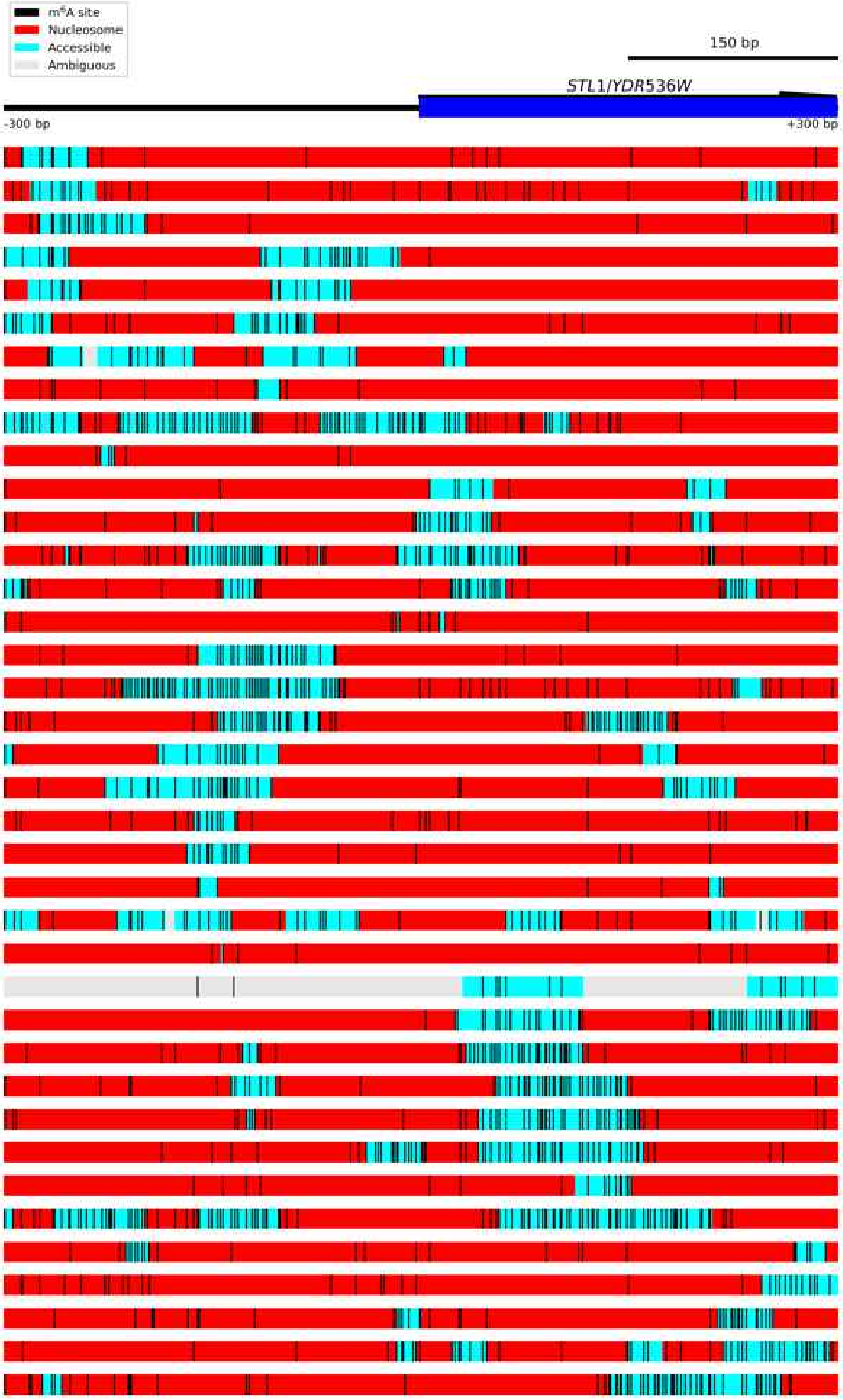

Figure S7.13 **Diagram of *STL1/YDR536W*.**
The *STL1/YDR536W* locus was an example of heterogeneous nucleosome positioning. Each block below the gene locus represents one read. Accessible regions are indicated by cyan boxes, nucleosomes as red boxes, and ambiguous regions by grey boxes. Called $m^6A$ bases are indicated by vertical black lines.

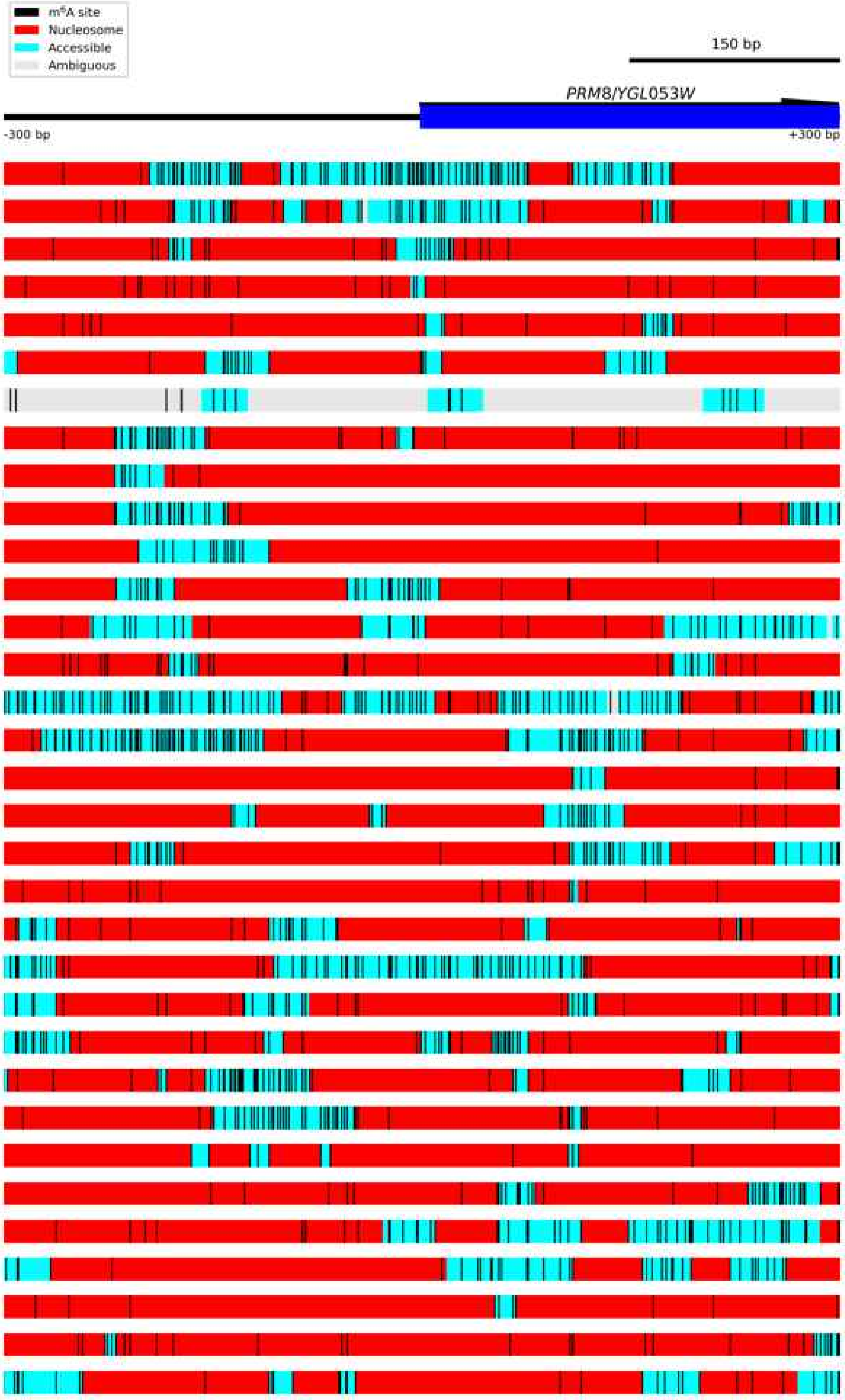

Figure S7.14 **Diagram of *PRM8/YGL053W*.**
The *PRM8/YGL053W* locus was an example of heterogeneous nucleosome positioning. Each block below the gene locus represents one read. Accessible regions are indicated by cyan boxes, nucleosomes as red boxes, and ambiguous regions by grey boxes. Called $m^6A$ bases are indicated by vertical black lines.

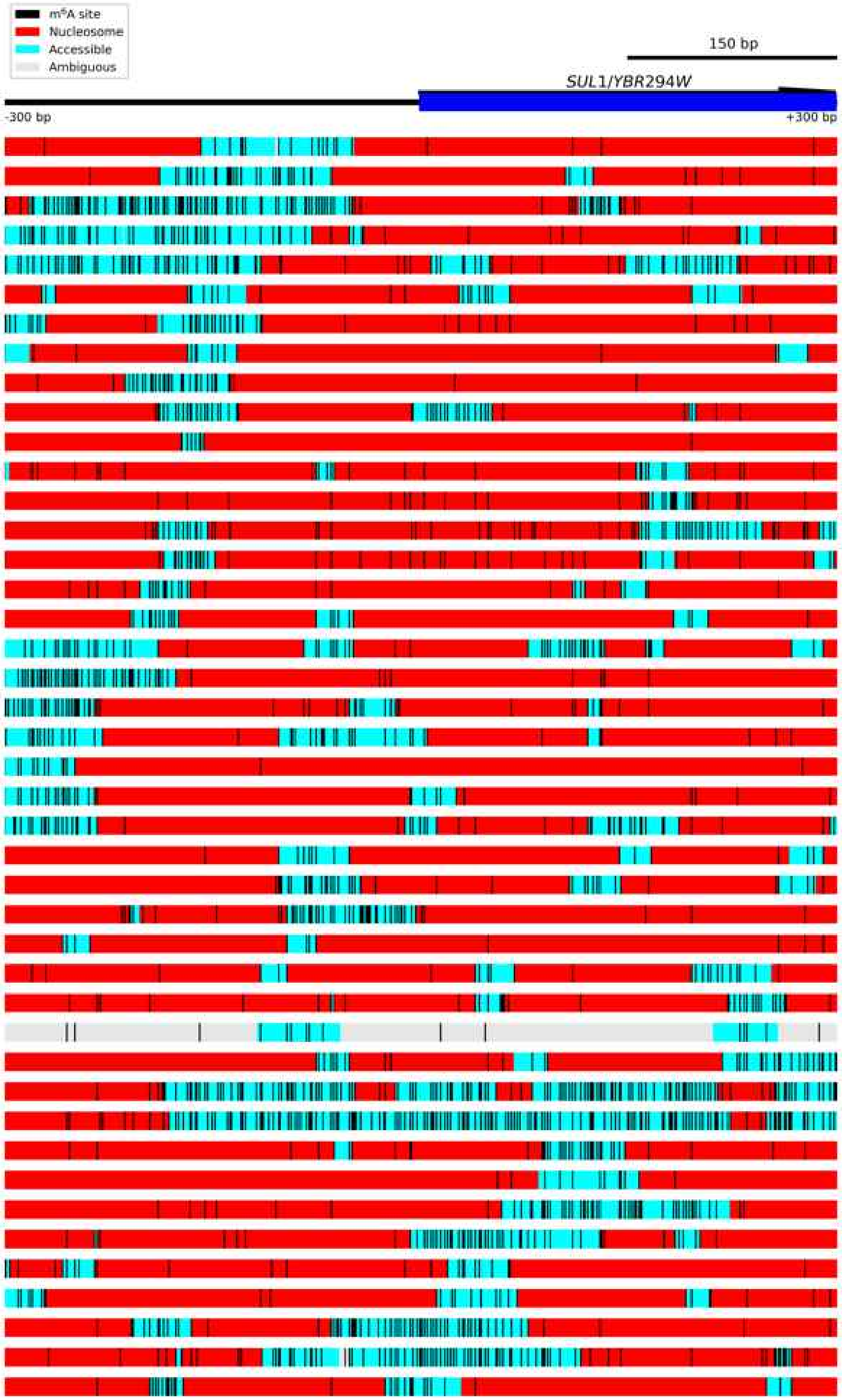

Figure S7.15 **Diagram of *SUL1/YBR294W*.**
The *SUL1/YBR294W* locus was an example of heterogeneous nucleosome positioning. Each block below the gene locus represents one read. Accessible regions are indicated by cyan boxes, nucleosomes as red boxes, and ambiguous regions by grey boxes. Called $m^6A$ bases are indicated by vertical black lines.

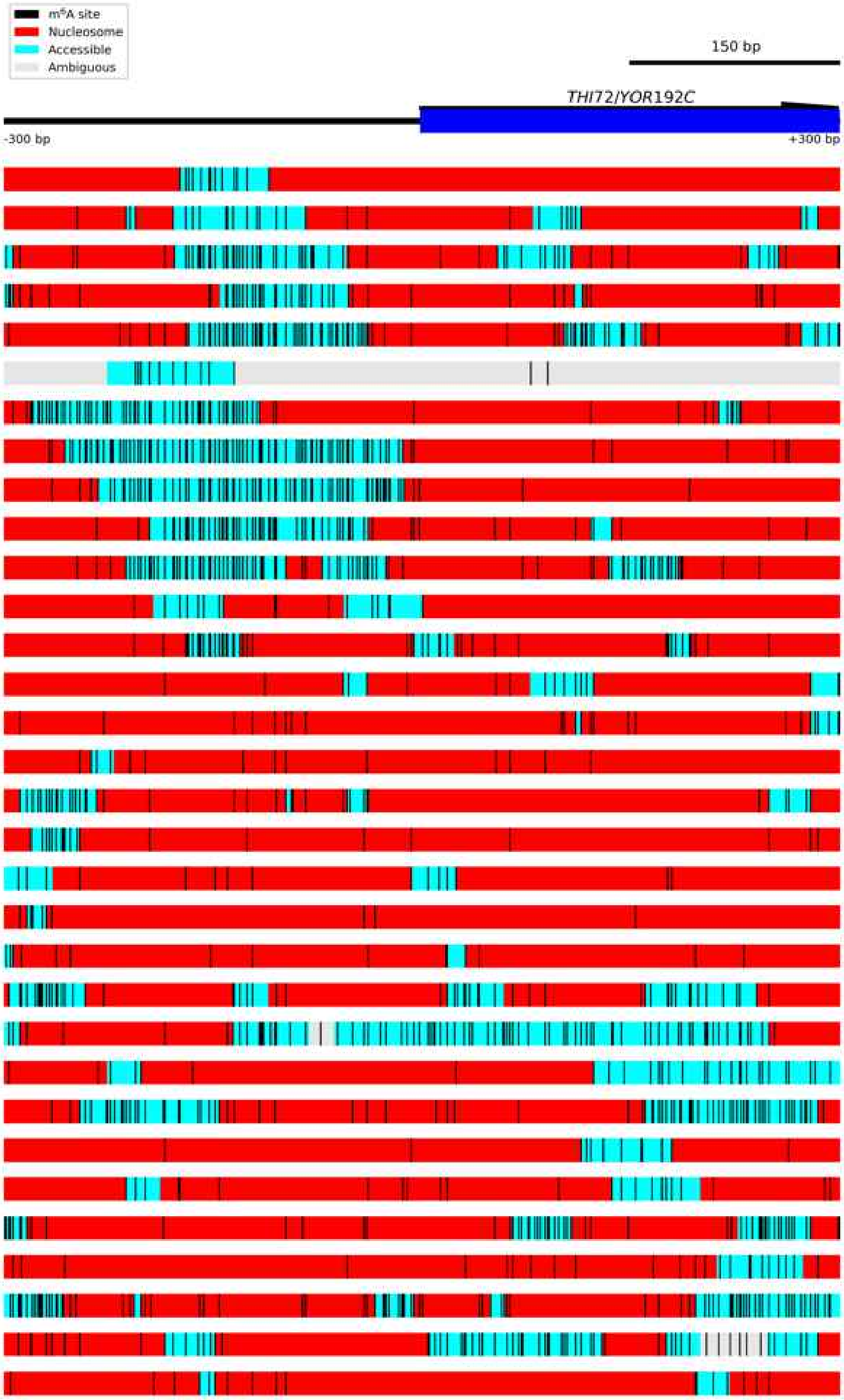

Figure S7.16 **Diagram of *THI72/YOR192C*.**
The *THI72/YOR192C* locus was an example of heterogeneous nucleosome positioning. Each block below the gene locus represents one read. Accessible regions are indicated by cyan boxes, nucleosomes as red boxes, and ambiguous regions by grey boxes. Called $m^6A$ bases are indicated by vertical black lines.

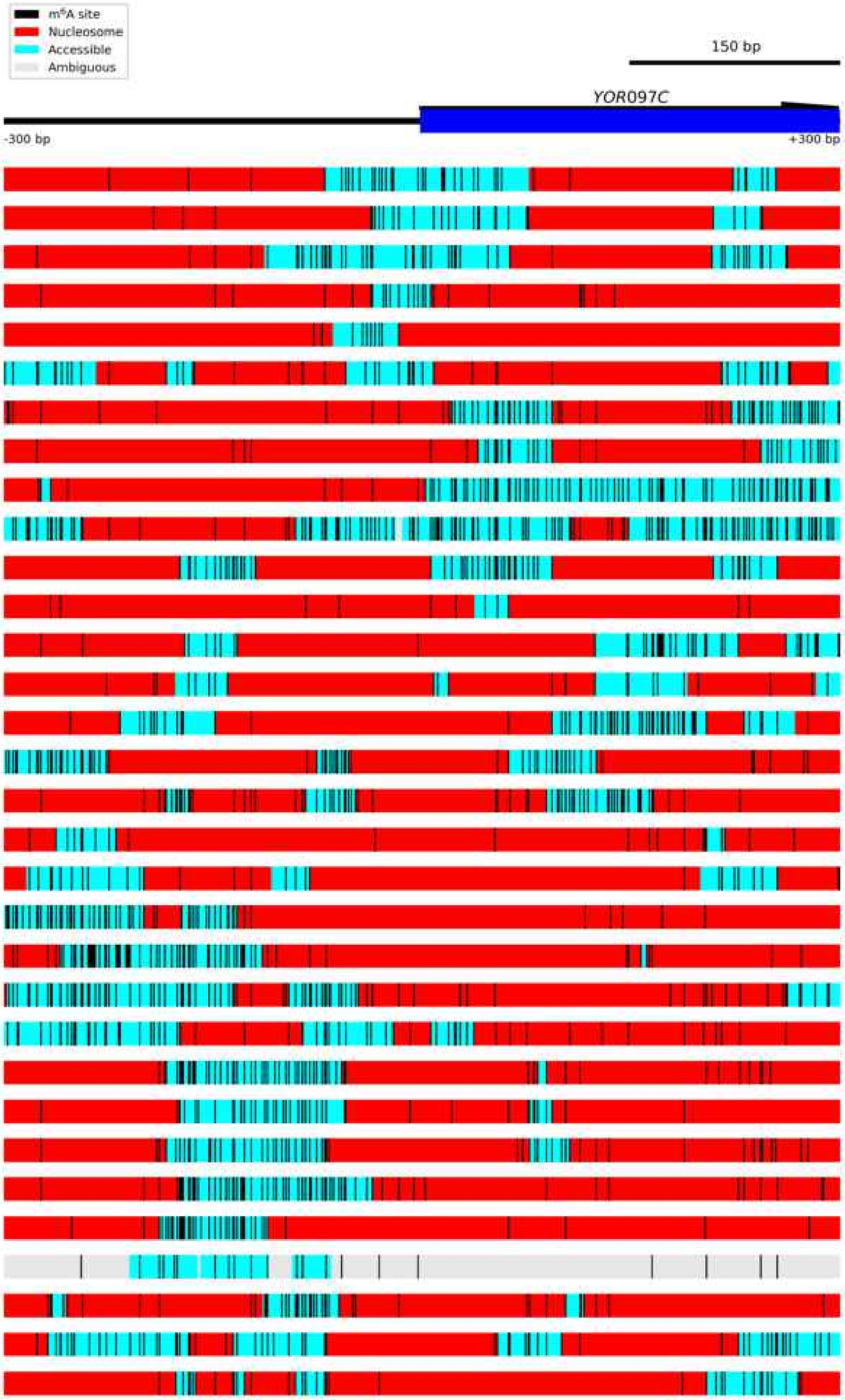

Figure S7.17 **Diagram of *YOR097C*.**
The *YOR097C* locus was an example of heterogeneous nucleosome positioning. Each block below the gene locus represents one read. Accessible regions are indicated by cyan boxes, nucleosomes as red boxes, and ambiguous regions by grey boxes. Called $m^6A$ bases are indicated by vertical black lines.

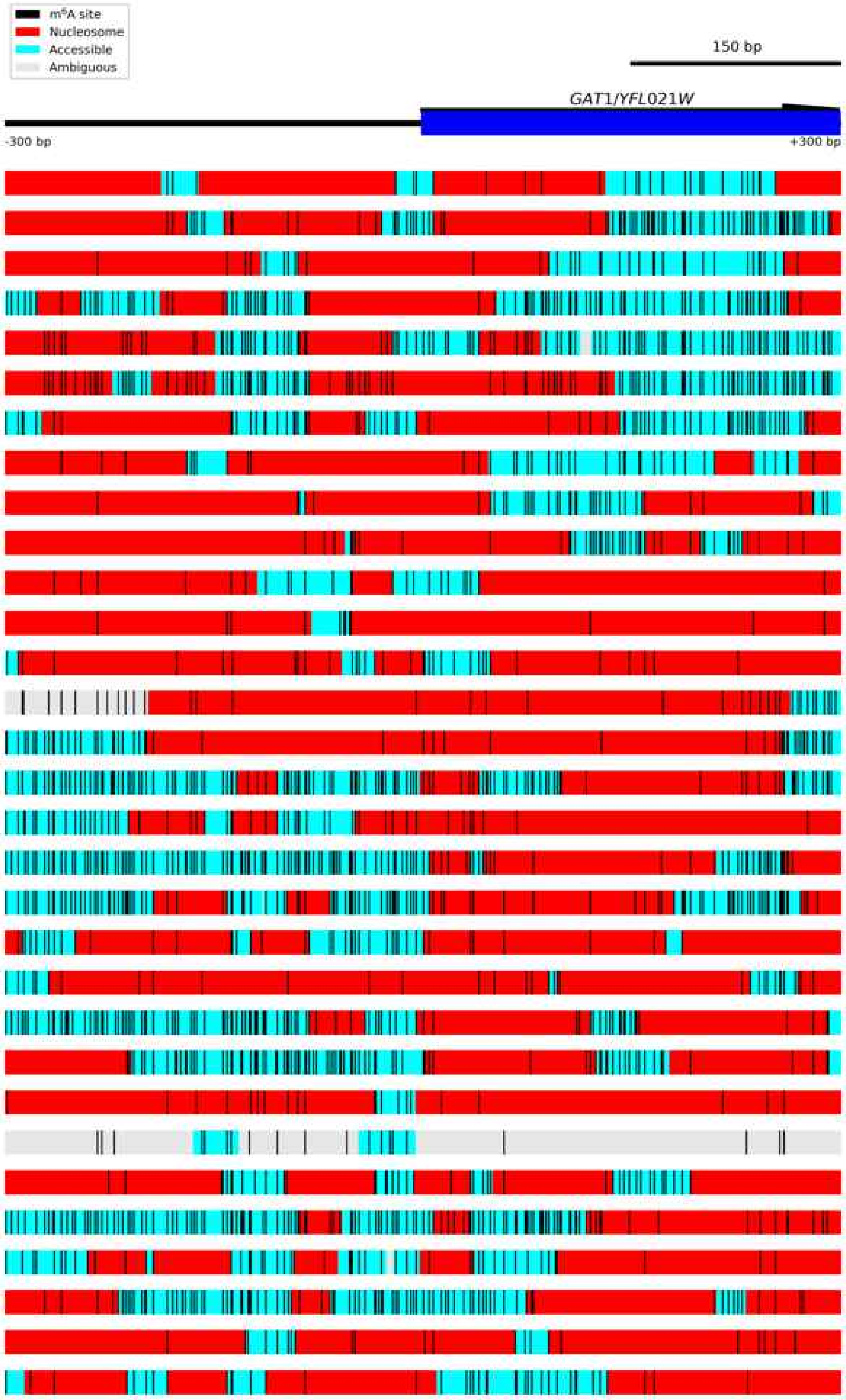

Figure S7.18 **Diagram of *GAT1/YFL021W*.**
The *GAT1/YFL021W* locus was an example of heterogeneous nucleosome positioning. Each block below the gene locus represents one read. Accessible regions are indicated by cyan boxes, nucleosomes as red boxes, and ambiguous regions by grey boxes. Called $m^6A$ bases are indicated by vertical black lines.

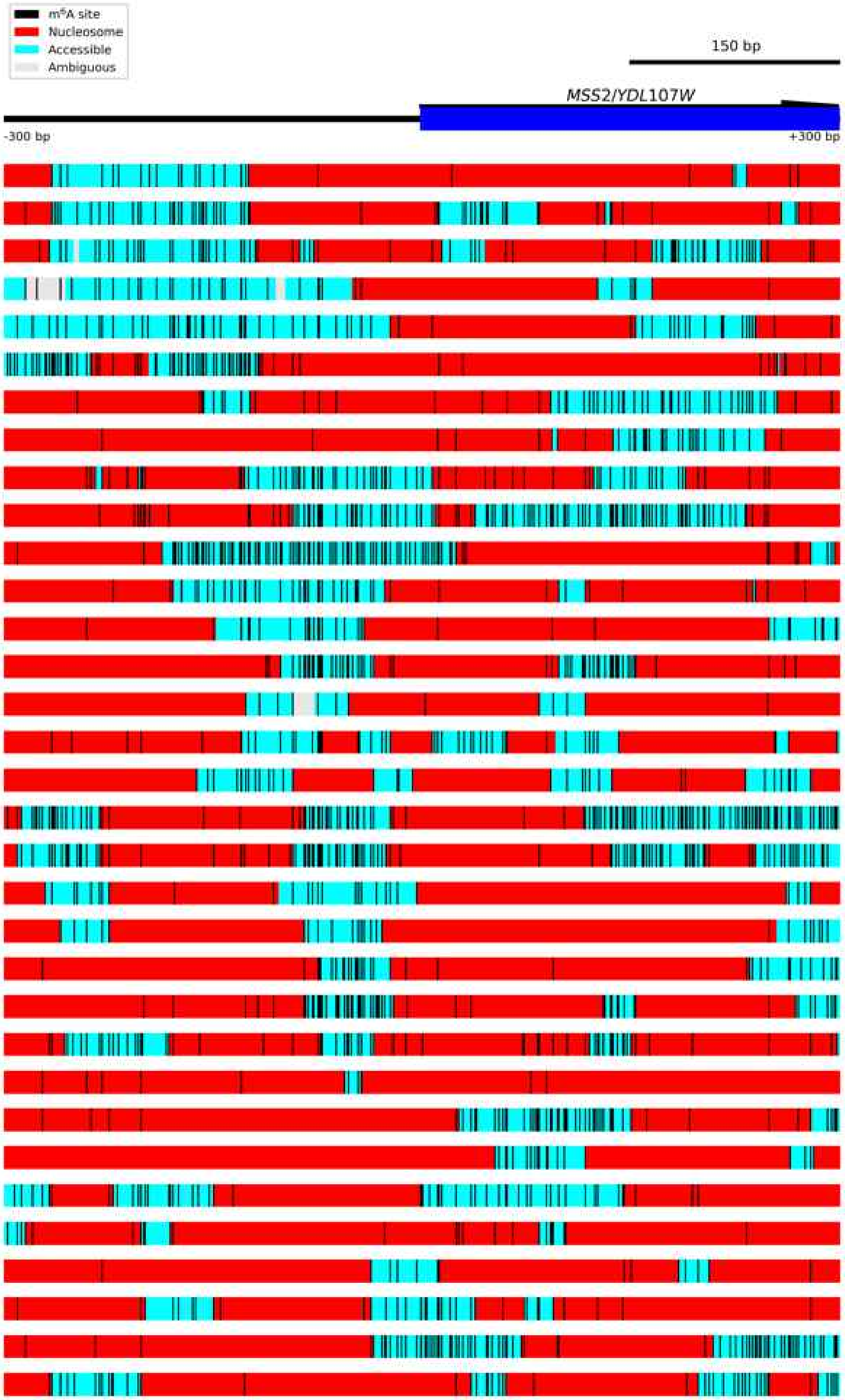

Figure S7.19 **Diagram of *MSS2/YDL107W*.**
The *MSS2/YDL107W* locus was an example of heterogeneous nucleosome positioning. Each block below the gene locus represents one read. Accessible regions are indicated by cyan boxes, nucleosomes as red boxes, and ambiguous regions by grey boxes. Called $m^6A$ bases are indicated by vertical black lines.

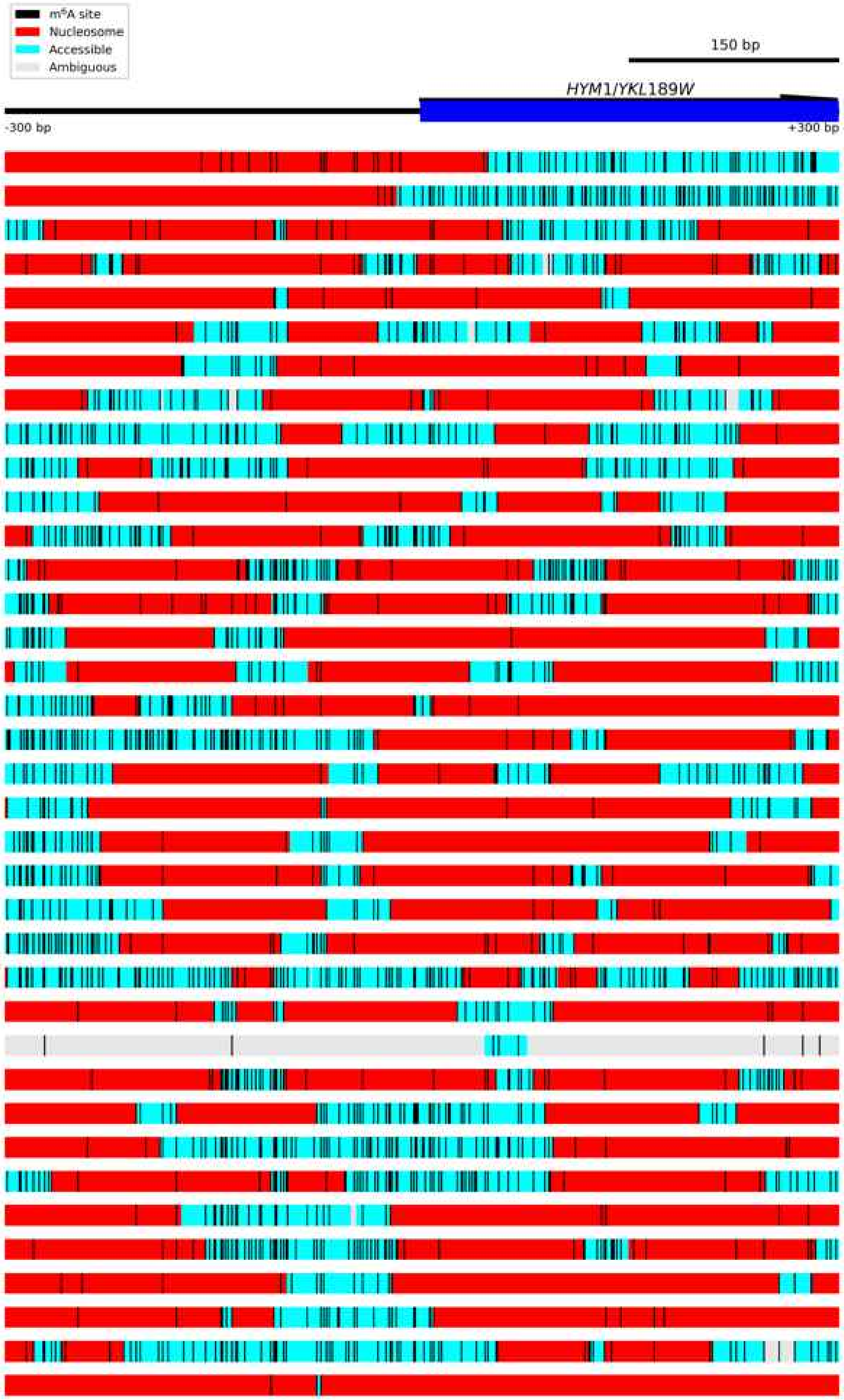

Figure S7.20 **Diagram of *HYM1/YKL189W*.**
The *HYM1/YKL189W* locus was an example of heterogeneous nucleosome positioning. Each block below the gene locus represents one read. Accessible regions are indicated by cyan boxes, nucleosomes as red boxes, and ambiguous regions by grey boxes. Called m$^6$A bases are indicated by vertical black lines.
